# Chemo-mechanotherapy of fibrosis: dynamic control of biological materials through tissue-architecture-dependent crosslink disruption

**DOI:** 10.1101/2025.09.01.673308

**Authors:** Wenyu Kong, Meiyue Song, Xiangjun Peng, Lu Bai, Jia’nan Zeng, Kaini Liang, Yuhong Jin, Jiaxin Wang, Xue Wang, Yuxuan Huang, Yudi Niu, Lyu Zhou, Xi-Qiao Feng, Chen Wang, Guy M. Genin, Jing Wang, Yanan Du

**Affiliations:** School of Biomedical Engineering, Tsinghua-Peking Joint Center for Life Sciences, Tsinghua University; Beijing, 100084, China; State Key Laboratory of Respiratory Health and Multimorbidity, Institute of Basic Medical Sciences Chinese Academy of Medical Sciences, School of Basic Medicine Peking Union Medical College; Beijing, 100005, China; National Key Laboratory of Kidney Diseases; Beijing, 100039, China; Institute of Biomechanics and Medical Engineering, Applied Mechanics Laboratory, Department of Engineering Mechanics, Tsinghua University; Beijing, 100084, China; Department of Mechanical Engineering and Materials Science, NSF Science and Technology Center for Engineering Mechanobiology, Washington University in St. Louis; St. Louis, 63130, USA; Internal Medicine, Harbin Medical University; Harbin, 150081, China; Department of Respiratory Medicine, the Second Affiliated Hospital of Harbin Medical University; Harbin, 150086, China

## Abstract

Extracellular matrix (ECM) stiffness drives cell fate, but tissues are nonlinear and difficult to modify chemically without harming patients. Here, we show that nonlinear mechanical responses of fibrous and porous tissues can be controlled by mechanically modulating crosslinks. Through integrated *in vivo, in vitro*, and computational approaches, we found that physiological dynamic stretching disrupts pathological advanced glycation end-product (AGE) crosslink-induced fibrogenesis in an architecture-dependent manner. This was effective in lung-like porous scaffolds but not in liver-like fibrous matrices, revealing a critical structure-property relationship. Computational and *in vitro* models established that this modulates cell-ECM feedback, and that controlling mechanical crosslinking modulates key nonlinearity of networked solids. This discovery inspired a non-invasive, “mechanotherapuetic” ventilation protocol, which reversed pulmonary fibrosis in mice by physically disrupting ECM pathological crosslinks. The therapeutic efficacy was further amplified when combined with pharmacological AGE inhibition. These findings establish design principles for dynamically reprogrammable biological materials and mechano-targeted therapies.

## Introduction

Controlling the mechanical properties of biological materials in living systems represents a grand challenge in materials science^1,2^. Gradual changes over time are associated with fibrosis and cancer^3^, and these changes are in general irreversible^4^. Thus, while synthetic materials can be designed with programmable properties that change in response to external stimuli, biological tissues have largely been treated as static materials whose properties are difficult to modify once formed^5^. The ability to dynamically modify tissue mechanics through external inputs would revolutionize fields from tissue engineering to regenerative medicine, but has remained elusive due in part to limited understanding of how mechanical forces interact with biological networks in realistic tissue architectures.

Crosslinking is fundamental to the mechanical behavior of biological materials, determining properties from tissue stiffness to failure mechanisms^6^. Advanced glycation end-product (AGE) crosslinking, in particular, represents a pathological modification that progressively stiffens tissues and is associated with aging and disease across multiple organ systems^7–10^ However, unlike synthetic crosslinked networks where properties are fixed during curing, biological crosslinking networks exist in dynamic mechanical environments that could potentially be harnessed to modify material properties post-formation.

The lung presents a unique materials science challenge and opportunity in this context. Unlike static organs, lung tissue experiences continuous cyclic deformation with strains up to 20% during normal respiration.^11^ This dynamic mechanical environment, combined with the lung’s distinctive porous architecture, creates conditions that are not found in other biological materials^12,13^ and that are needed for homeostasis. The lung also highlights the exquisite sensitivity of tissues to homeostatic imbalances: physiological dynamic stretch maintains lung cell identity and regulating stem cells,^14^ and abnormal tension can drive progressive lung fibrosis.^15^ We hypothesized that the combination of architecture and mechanical loading might enable unprecedented control over tissue crosslinking and mechanical properties.

This is particularly impactful in the context of fibrosis, a pathological stiffening of soft tissues that affects virtually every organ system.^16^ Here, changes to extracellular matrix (ECM) and fibroblast cells occur in tandem,^17,18^ leading to a cycle that in general cannot be reversed. Idiopathic pulmonary fibrosis exemplifies the challenge, with FDA-approved drugs only slow disease progression,^19^ and with a median survival of 3-4 years and a 5-year survival rate of 20-50%.^20,21^ This limited therapeutic success may stem from targeting individual disease components in isolation: neither reducing ECM accumulation^22–24^ nor altering cell behavior^19,25,26^ can halt or reverse disease progression. Intriguingly, idiopathic pulmonary fibrosis is associated with elevated serum AGE levels^27^ and prevalence of diabetes.^28–30^ The receptor for AGE (RAGE), while expressed more in healthy lung tissue than in other organs,^31^ paradoxically decreases in pulmonary fibrosis,^32,33^ contrasting with the increased RAGE expression in liver fibrosis.^34^ This motivated a hypothesis that tissue architecture and cyclic stretching affects AGE-mediated signaling and mechanobiological feedback loops in fibrosis.

To test this hypothesis, we developed an integrated experimental and computational approach that enabled mechanical control of AGE crosslinking in a range of biomaterials and tissues, and that revealed how dynamic control of biomaterial properties enables both control and reversal of fibrosis progression both *in vitro* and *in vivo*.

## Results

### AGE crosslinking creates tissue-specific mechanical environments in biological materials

AGE crosslinking represents a pathological modification of biological materials that fundamentally alters tissue mechanical properties in organ-specific patterns. To establish the role of AGE-mediated crosslinking in pulmonary fibrosis ECM, we first examined human lung tissue samples from both healthy individuals and patients with established disease (**Fig. 1a**). Fibrotic samples showed extensive tissue remodeling characterized by clusters of activated fibroblasts, with increased α-smooth muscle actin (α-SMA) staining, substantial collagen I (COL-1) deposition, and notably increased AGE crosslinking in fibrotic regions. Paradoxically, RAGE expression was decreased in fibrotic lung tissue, contrasting with the findings in liver fibrosis where RAGE expression increases (**Fig. 1b**).

**Fig. 1.**
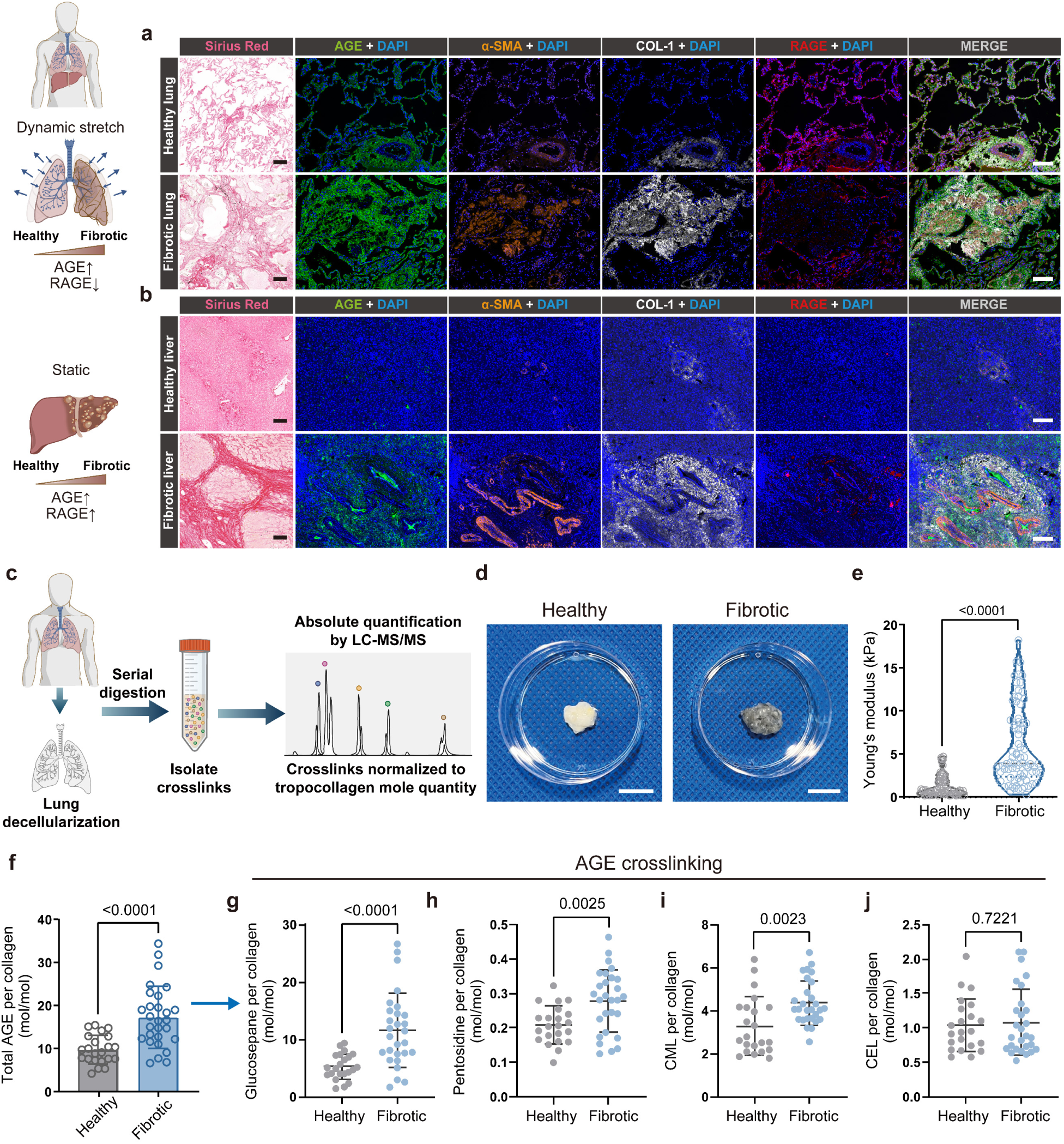
Advanced glycation end-product (AGE)-crosslinked ECM distinguishes pulmonary fibrosis with paradoxical RAGE downregulation. (**a-b**) Representative images comparing healthy and fibrotic tissue in human lung (**a**) and liver (**b**), showing distinct patterns of collagen organization (Sirius red), AGE crosslinking, collagen I (COL-1), α-smooth muscle actin (α-SMA), and receptor for advanced glycation end products (RAGE). Note the paradoxical decrease of RAGE in fibrotic lung versus increase in fibrotic liver. Scale bars, 200 μm. (**c**) Schematic of the Absolute Quantification of Matrix-specific Crosslinking (AQMC) method used to analyze crosslinking density. (**d**) Representative bright-field images of decellularized extracellular matrix (ECM) from healthy and fibrotic human lungs. Scale bars, 1 cm. (**e**) Effective Young’s moduli of decellularized human lung ECM measured by atomic force microscopy (AFM), showing significant stiffening in fibrotic tissue (*n* ≥ 63 measurements per group, randomly selected locations). (**f**) Total AGE crosslinking density in decellularized human lung ECM, revealing significant increase in fibrotic tissue (*n* ≥ 22 independent patient samples). (**g-j**) Quantification of specific AGE crosslinks in decellularized lung ECM: (**g**) glucosepane, (**h**) pentosidine, (**i**) CML (N^ε^-(1-carboxymethyl)-L-lysine), and (**j**) CEL (N^ε^ -(1-carboxyethyl)-L-lysine), all significantly elevated in lung ECM (*n* = 6 ECM samples from 5 independent mice per condition). Statistical analysis performed using two-tailed unpaired t-test, exact *p* values labeled. Results presented as mean ± S.D.

Following decellularization (**Extended Data Fig. 1a**), fibrotic ECM appeared less translucent than healthy ECM (**Fig. 1d**), and exhibited significantly increased effective modulus as measured by atomic force microscopy (AFM) (**Fig. 1e**). These changes were associated with AGE crosslinking, as measured by absolute quantification of matrix-specific crosslinking (AQMC) analysis^35^ (**Fig. 1c, Extended Data Fig. 2a**). Total AGE crosslinking was elevated by a factor of 1.77 (*p* < 0.0001) in fibrotic decellularized lung ECM (**Fig. 1f**), with significant increases in all four major types: N^ε^-carboxymethyl-lysine (CML), (N^ε^-carboxyethyl-lysine (CEL), glucosepane, and pentosidine (**Fig. 1g-j**). Importantly, AGE crosslinking levels exceeded both TGM and LOX crosslinking in fibrotic lung ECM (**Extended Data Fig. 2b**). These findings establish AGE crosslinking as a critical determinant of tissue mechanical environment and reveal organ-specific patterns that suggest different materials-based therapeutic approaches.

### Pathological crosslinking drives mechanical property changes in a controlled biological materials system

To validate the role of pathological crosslinking in controlling biological material properties, we employed a murine model system that recapitulates key mechanical and biochemical features of tissue stiffening. We induced pulmonary fibrosis in mice using bleomycin (**Extended Data Fig. 3a**), under conditions that produce persistent fibrosis without self-recovery over 8 weeks^36^; although lower doses produce fibrosis that resolves over several weeks^37^, the doses used here lead to persistent and progressive disease^38^. Bleomycin-treated mice showed characteristic signs of pulmonary fibrosis including rapid weight loss, increased ECM accumulation (confirmed by Masson staining and micro-CT, **Extended Data Fig. 3b, d, e**), and elevated blood glucose (**Extended Data Fig. 3c**). Lung function tests revealed increased inspiratory resistance and decreased compliance and capacity (**Extended Data Fig. 3f**), indicating compromised respiratory function. As with human samples, mouse tissue showed the paradoxical increase in RAGE and decreased RAGE expression (**Extended Data Fig. 3g-h**).

Following whole-lung-perfusion decellularization (**Extended Data Fig. 1b**), mouse lung tissue retained its characteristic architecture, with healthy tissue showing a porous structure a that contrasts with the previously published fibrous structure of decellularized liver ECM^35^ (**Fig. 2a-b**). Decellularized fibrotic lung ECM displayed a similar porous structure, but with irregular morphology, elevated dry weight, increased pore size, greater pore wall thickness, and increased modulus compared to healthy decellularized lung ECM, indicating enhanced ECM deposition and densification during the progression of fibrosis (**Fig. 2c-d, Extended Data Fig. 1c-d**).

**Fig. 2.**
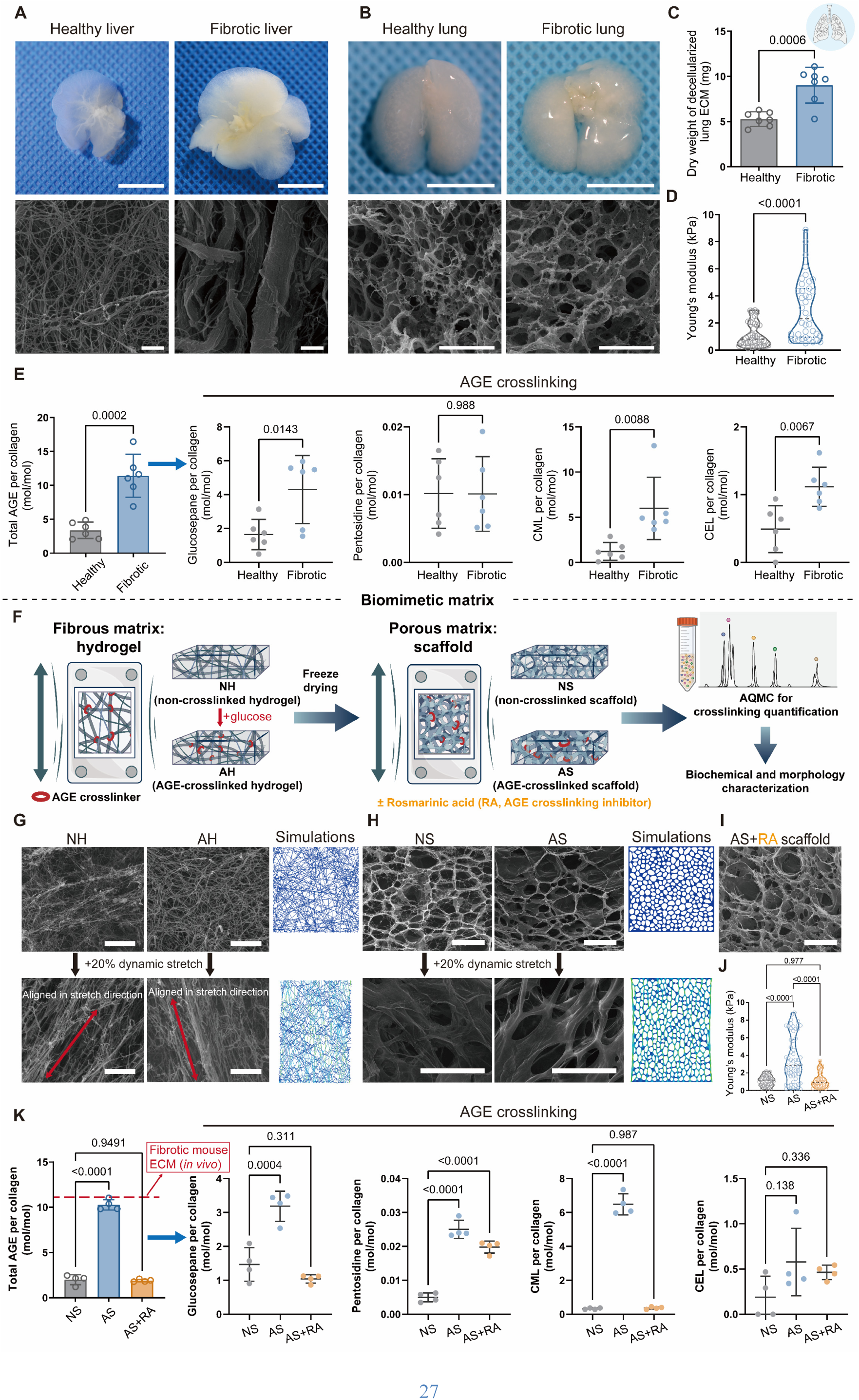
AGE crosslinking drives ECM architectural and mechanical alterations in pulmonary fibrosis that can be reversed *in vitro* by targeted interventions. (**a-b**) Comparison of decellularized ECM architecture in healthy and fibrotic tissue from liver (**a**, images adapted from *Nat. Biomed. Eng.* 7, 1437-1454 (2023)^35^) and lung (**b**), showing distinct organ-specific patterns through bright-field microscopy (scale bars, 1 cm) and scanning electron microscopy (SEM) (scale bars, 100 μm). (**c**) Total mass of decellularized lung ECM showing significant increase in fibrotic tissue (*n* = 7 independent samples). (**d**) Effective Young’s modulus measured by AFM demonstrating matrix stiffening in fibrotic lung. (**e**) AGE crosslinking analysis of healthy versus bleomycin (BLM)-treated mouse lung ECM (*n* = 6 ECM samples from 5 independent mice per condition), showing revealing 3.4-fold increase in bleomycin-treated mouse lung ECM, with breakdown by specific AGE types (CML, CEL, glucosepane, and pentosidine). (**f**) Schematic of biomimetic matrices that recapitulate both liver-like fibrous and lung-like porous ECM architectures, integrated with dynamic stretching capability. (**g**) SEM images and computational simulations showing structural changes in fibrous hydrogels (non-crosslinked: NH; AGE-crosslinked: AH) with AGE crosslinking and 20% dynamic strain. Scale bars, 200 μm. (**h-i**) SEM images and computational simulations demonstrating how AGE crosslinking alters porous scaffolds, with rosmarinic acid (RA) treatment reversing these changes, and 20% dynamic stretch altering architecture (non-crosslinked scaffold: NS; AGE-crosslinked scaffold: AS; AGE scaffold treated with rosmarinic acid: AS+RA). Scale bars, 200 μm. (**j**) AFM-based Young’s modulus measurements confirming that RA treatment restores normal mechanics in AGE-crosslinked scaffolds (*n* ≥ 36 measurements from ≥ 10 fields per sample). (**k**) Quantification of AGE crosslinking in biomimetic scaffolds (*n* = 4 independent matrix samples), with breakdown by specific AGE type, demonstrating successful recapitulation of the fibrotic phenotype and its reversibility with RA treatments. Statistical analysis performed using two-tailed unpaired t-test, and one-way ANOVA with Tukey’s post-hoc test, exact *p* values labeled. Results presented as mean ± S.D.

AGE crosslinking increased 3.4-fold in fibrotic ECM compared to healthy tissue, with CML showing the highest levels, followed by glucosepane, CEL, and pentosidine (**Fig. 2e**). Lung ECM in the bleomycin-induced pulmonary fibrosis model revealed no significant increase in TGM crosslinking, while LOX crosslinking remained low (**Extended Data Fig. 2c**). This might explain why previous attempts to treat pulmonary fibrosis by targeting LOX-mediated crosslinking have shown limited success, despite the clear role of matrix mechanics in disease progression^39^.

RNA-seq analysis of primary pulmonary fibroblasts (pPFBs) isolated from healthy and fibrotic mouse lungs revealed upregulation of fibrosis-related genes (*Acta2*, *Actg2*, *Col11a1*) in fibrotic conditions (**Extended Data Fig. 4a**). Pathway enrichment and gene set enrichment analysis (GSEA) showed enrichment for focal adhesions, ECM-receptor interactions, and the AGE-RAGE signaling pathway, the last of these being a pathway characteristic of diabetic complications (**Extended Data Fig. 4b-e**). While most AGE-RAGE pathway genes were upregulated, *Ager* was downregulated (**Extended Data Fig. 4f**), suggesting a surprising mismatch between decreased RAGE expression and increased AGE-RAGE signaling in pulmonary fibrosis.

These results suggested AGE-mediated crosslinking as a key feature of fibrotic lung ECM, contributing to both mechanical and biochemical alterations in diseased tissue. The distinct pattern of RAGE expression in pulmonary versus liver fibrosis suggested tissue-specific regulation, potentially related to the different biomechanical environments in the dynamically stretching lung versus the static liver. This controlled biological materials system demonstrates that pathological crosslinking creates measurable changes in tissue architecture and mechanical properties, providing a platform for testing materials-based interventions. This observation prompted us to investigate how mechanical stretches might influence the AGE-RAGE axis in pulmonary fibrosis.

### Biomimetic materials systems reveal architecture-dependent responses to mechanical intervention

To isolate the effects of tissue architecture from biochemical factors, we developed biomimetic materials systems that recapitulate the distinct structural features of different organ microenvironments (**Fig. 2f**). This enabled us to search for pathways to interrupt this mechanobiological feedback loop. We created non-crosslinked scaffolds (NS) and AGE-crosslinked scaffolds (AS) that mimicked the porous architecture of lung ECM, alongside a comparison case: non-crosslinked hydrogels (NH) and AGE-crosslinked hydrogels (AH) that mimicked the fibrous architecture of liver ECM. These ECM scaffolds were cast in custom molds that could be placed on a stretching device (**Supplementary Video 1**, **Extended Data Fig. 5**) to simulate human respiratory motion (20% strain applied at 15 cycles/min in 2-second stretch/contraction cycles^11^.

In fibrous, liver-like hydrogels, AGE crosslinking caused collagen to transition from the fine, uniform structure representative of healthy liver ECM to a bundled, aligned structure (scanning electron microscopy (SEM) images, **Fig. 2**), consistent with computational predictions of stretch-induced fiber alignment in a Mikado discrete network (**Fig. 2g, Supplementary Video 2**). Porous scaffolds showed lung-like porous ECM architecture (**Fig. 2h**), with AGE-crosslinking increasing pore size and wall thickness as in fibrotic lung ECM (**Extended Data Fig. 6a, b**). Consistent with computer simulations of porous 2D lattices, dynamic stretching did not alter these pores substantially (**Fig. 2h, Extended Data Fig. 6a, b, Supplementary Video 3**). AGE crosslinking significantly increased scaffold modulus as measured by AFM (**Fig. 2j**).

To test the role of AGEs in these mechanical and architectural changes to the lung scaffolds, the *in vitro* systems were treated with rosmarinic acid (RA), a specific AGE crosslinking inhibitor. RA reversed the changes associated with AGE crosslinking, restoring pore size, pore wall thickness, and modulus to levels comparable to non-crosslinked scaffolds (**Extended Data Fig. 6a, b, Fig. 2j**). AQMC analysis confirmed that AGE-crosslinked scaffolds presented crosslinking levels similar to the fibrotic lung ECM of the bleomycin model, and that RA treatment effectively reduced this crosslinking (**Fig. 2k**).

The *in vitro* system thus replicated key features of fibrotic lung ECM and physiological respiratory movements, providing a platform for investigating the role of AGE crosslinking in pulmonary fibrosis. These engineered materials systems demonstrate that both chemical and mechanical interventions can restore normal material properties, with efficacy dependent on architectural design principles.

### Computational modeling reveals mechanisms of mechanically-controlled crosslinking networks

To understand the fundamental mechanisms underlying mechanical control of crosslinking networks, we developed computational models that predict how mechanical forces alter crosslink density and network properties. A computational framework based on a Mikado discrete network model (**Fig. 3a**) revealed the well-known result that matrix stretching creates two distinct fiber populations: those which bear substantial load and align with the principal loading direction, and those which are less affected by the mechanical loads.

**Fig. 3.**
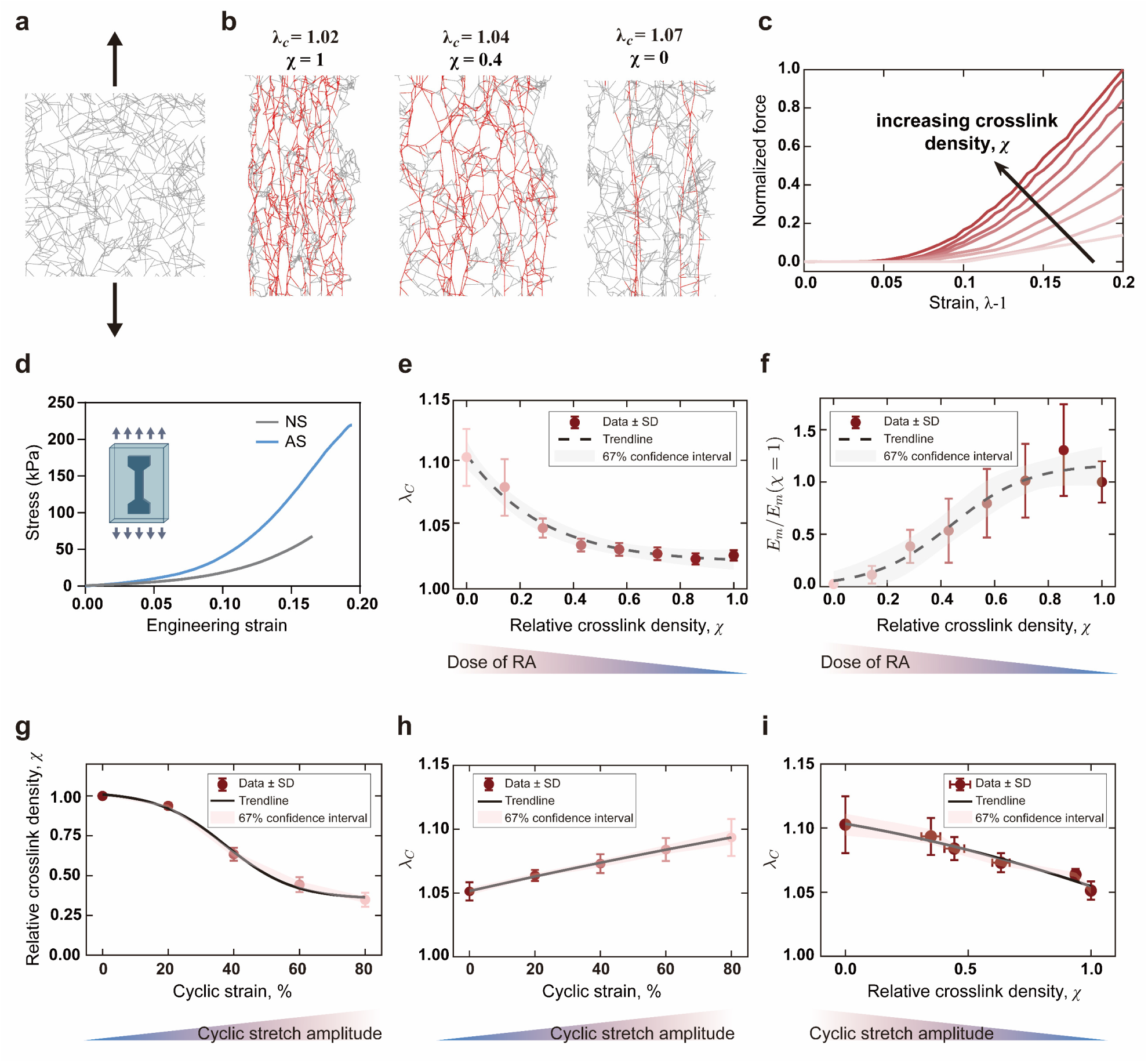
Computational modeling reveals mechanisms by which crosslink disruption can reverse fibrotic ECM mechanics. (**a**) Mikado discrete network model of collagen matrix under tension, with load-bearing fibers highlighted in red, demonstrating how crosslink density affects fiber recruitment and alignment. (**b**) Quantitative analysis showing the impact of random crosslink deletion on fiber alignment under mechanical stretch. (**c**) Stress-strain relationships predicted for collagen networks with varying crosslink densities, demonstrating how crosslinking alters the characteristic strain-stiffening transition points. (**d**) Experimental validation comparing computationally predicted and experimentally measured mechanical behavior of non-crosslinked (NS) and AGE-crosslinked (AS) scaffolds. (**e-f**) Effects of crosslink density 𝜒 on: (**e**) critical principal stretch 𝜆_c_at which strain-stiffening begins and (**f**) baseline Young’s modulus *E_m_* of the networks. (**g-h**) Effects of cyclic stain amplitude on: (**g**) crosslink density 𝜒 and (**h**) critical principal stretch 𝜆_c_. (**i**) The relationship between crosslink density 𝜒 and critical principal stretch 𝜆_c_ under different cyclic stretch amplitudes, revealing the mechanical basis for potential mechanotherapy. Statistical analysis performed using two-tailed unpaired t-test. Results presented as mean ± S.D.

To model the effect of chemical deletion of crosslinks, we deleted crosslinks at random, the proportion of load-bearing fibers decreased with random deletion of crosslinkers (**Fig. 3b**), with crosslinking density directly influencing ECM organization. Stress-strain relationships derived from our simulations exhibited a characteristic “knee” point representing strain stiffening (**Fig. 3c**), which aligned well with experimental observations (**Fig. 3d**). Increased crosslinking density shortened this transition point and enhanced the stiffening process. Quantifying this revealed a strong correlation between crosslink density 𝜒 and critical principal stretch 𝜆_c_, with randomly removed crosslinks, having a more minor effect on the baseline Young’s modulus 𝐸_m_of the ECM at stretches below 𝜆_c_ (**Fig. 3e, f**).

To model the effects of cyclic stretching on ECM mechanics, we stretched this computational model, computed the stresses in the collagen network, and deleted the crosslinks stressed beyond a critical level. This model cyclic tensile straining also reduced both 𝜒 and and 𝜆_c_, as chemical deletion of crosslinks did, but had less of an effect on the tangent modulus beyond 𝜆_c_ (**Fig. 3g-i**).

These computational insights reveal that crosslink density directly controls critical mechanical transitions in biological materials, providing design principles for mechanically-controlled therapeutic interventions. Because reduction of 𝜆_c_ has been shown to reduce the tendency of fibroblasts to become activated^40^, we next tested the hypothesis that targeting ECM crosslinkers could be an effective strategy for interrupting the mechanobiological feedback loop driving fibrosis progression.

### Mechanical forces enable tissue-specific control of biological material properties

The discovery that mechanical forces can disrupt pathological crosslinking led us to investigate whether this represents a general materials control mechanism or depends on specific tissue architectures. We assessed biological responses of pPFBs cultured on our lung-like and liver-like *in vitro* models (**Fig. 4a, Extended Data Fig. 7a**). AGE-crosslinked scaffolds consistently promoted pPFB activation, as evident from increased α-SMA expression, F-actin organization, and paxillin adhesions (**Fig. 4b**). However, the application of 20% dynamic stretch revealed a striking, matrix-specific response: 20% dynamic stretch of AGE-crosslinked porous scaffolds (AS) suppressed pPFB activation, upregulated RAGE expression, and returned pPFBs to a more quiescent state with reduced pseudopodia (**Fig. 4b-e**), but these beneficial effects were absent in fibrous hydrogels (**Extended Data Fig. 7b-e**).

**Fig. 4.**
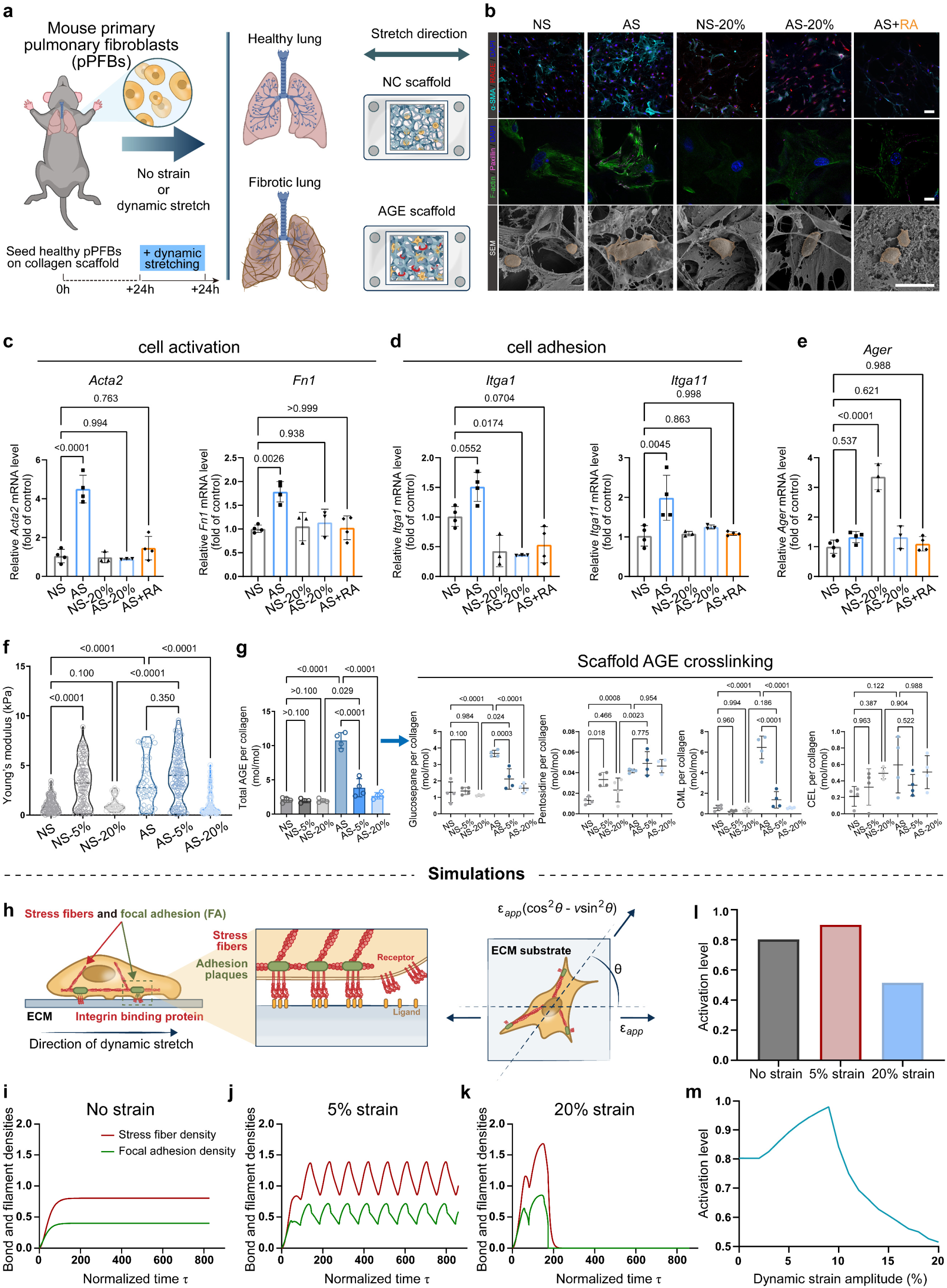
Physiological stretching reverses fibroblast activation in lung-like matrices through disruption of AGE crosslinking. (**a**) Schematic of experimental approach: primary pulmonary fibroblasts (pPFBs) isolated from lung tissue and cultured on engineered collagen scaffolds under specified conditions. (**b**) Characterization of pPFBs responses to different scaffold conditions, showing: immunofluorescence of α-SMA (activation marker, cyan), RAGE (red), and nuclei (blue) (top panels, scale bars 50 μm); F-actin cytoskeleton (green) and paxillin adhesions (amaranth) (middle panels, scale bars 10 μm); and SEM imaging of cell morphology (bottom panels, scale bars 20 μm). Note that 20% dynamic stretching reversed fibroblast activation in AGE-crosslinked scaffolds. (**c-e**) Quantification of relative mRNA expression in pPFBs showing: (**c**) activation markers (*Acta2*, *Fn1*), (**d**) adhesion molecules (*Itga1*, *Itga11*), and (**e**) AGE receptor (*Ager*) under different scaffold conditions, confirming that both mechanical stretching and rosmarinic acid treatment can reverse pro-fibrotic phenotypes (non-stretched: NS; AGE-crosslinked: AS; dynamically stretched to 20%: −20%; RA-treated: -RA). (**f**) Young’s modulus measurements demonstrating that 20% dynamic strain selectively reduces stiffness in AGE-crosslinked (AS) scaffolds but increases stiffness in non-crosslinked (NS) scaffolds at 5% strain (*n* ≥ 36 measurements from ≥10 fields per sample). (**g**) AGE crosslinking quantification showing progressive reduction with increasing strain amplitude, with 20% strain reducing crosslinking to near-normal levels (*n* = 4 independent samples). (**h**) Theoretical model of stress fiber orientation (𝜃) relative to stretch direction. (**i-k**) Computational predictions of stress fiber and focal adhesion dynamics under: (**i**) no strain (stable), (**j**) 5% strain (limit cycle response), and (**k**) 20% strain (showing depolymerization and loss of activation). (**l**) Predicted cellular activation levels under different strain conditions demonstrating biphasic response. (**m**) Phase diagram illustrating how cell activation depends on both matrix stiffness and strain amplitude, providing a mechanistic basis for mechanotherapy. Statistical analysis performed using one-way ANOVA with Tukey’s post-hoc test, exact *p* values labeled. Results presented as mean ± S.D.

The contrasting cellular responses between porous and fibrous matrices prompted investigation of their mechanical properties. 20% dynamic strain dramatically decreased the AFM indentation modulus of AGE-crosslinked porous scaffolds, while 5% dynamic strain had no effect **(Fig. 4f)**. In contrast, 5% dynamic strain increased the modulus of non-AGE-crosslinked porous scaffolds, while these changes in stiffness were lost with 20% dynamic strain (**Fig. 4f**). AGE crosslinking stiffened fibrous hydrogels, and 20% dynamic stretch reduced the AFM indentation modulus of both AGE-crosslinked and non-AGE-crosslinked fibrous hydrogels substantially (**Extended Data Fig. 7f**).

AQMC analysis showed that increased cyclic strain amplitude decreased AGE crosslinking, eventually down to levels comparable to those of non-crosslinked scaffolds **(Fig. 4g)**. AGE crosslinking decreased with 20% dynamic strain in the AGE-crosslinked fibrous hydrogels as well **(Extended Data Fig. 7g)**. Notably, these changes were specific to AGE crosslinks, dynamic stretching did not induce a significant change in the TGM crosslinking degree in neither the scaffolds nor the fibrous hydrogels **(Extended Data Fig. 8)**. This is consistent with earlier observations that imines (R2C=NR’) in glucosepane or pentosidine can undergo dynamic and reversible covalent bonding^41^.

Consistent with the fibrous hydrogels having moduli in a range in which fibroblasts are not known to be mechanosensitive (0.3-1.1 kPa)^42^ (**Extended Data Fig. 7b-f**), limited effects of cyclic stretch were observed. The hydrogel’s lower initial modulus (0.1-1.0 kPa) may explain its limited response to stretching. Additionally, fiber alignment in stretched hydrogels likely disrupted mechanotransduction, unlike the stable porous scaffold structure (**Fig. 2g-h**). The modulus of the NC-crosslinked scaffold increased after dynamic stretching to 5% strain **(Fig. 4f)**, consistent with alignment and recruitment driven strain stiffening that occurs independently of reduction in AGE crosslinking (**Fig. 2g-h**).

These effects were consistent with a model of changes to the dynamics of focal adhesions (FAs), clusters of ligand-receptor bonds that connect cells to the ECM, in response to cyclic stretch^43^. We modeled actin stress fibers attached to an elastic substrate via two FAs, with each fiber oriented at an angle *θ* to the stretch direction **(Fig. 4h)** and evaluated subsequent mechanotransduction (methods). Results aligned with experimental observations. In the absence of dynamic stretching, stress fibers and FAs stabilized (**Fig. 4i**). 5% cyclic stretching increased expression of stress fibers and FAs until a limit cycle state was reached (**Fig. 4j**), while 20% cyclic stretching elevated expression of stress fibers and FAs to higher levels that terminated with cytoskeletal depolymerization and loss of cell activation (**Fig. 4k**). Thus, dynamic stretching increased cell activation up to a maximum at ∼9% amplitude, and reduced to levels below unstretched baselines thereafter (**Fig. 4l-m**).

To verify that effects of stretch on cells could be associated with changes to ECM crosslinking, treated AGE-crosslinked scaffolds with rosmarinic acid to reduce AGE crosslinking. RA treatment, in the absence of stretch, also reversed pPFB activation markers (*Acta2*, *Fn1*), adhesion molecules genes (*Itga1*, *Itga11*), and the AGE receptor gene *Ager* (**Fig. 4c-e**). These results establish that mechanical forces can serve as precision tools for controlling biological material properties, but only when applied to tissues with appropriate architectural features.

### Mechanical and biochemical pathways independently control cellular responses to material properties

To dissect how material stiffness and crosslinking chemistry independently influence cellular behavior, we designed experiments and simulations that isolate mechanical from biochemical effects. We first studied biochemical effects of AGE crosslinking at constant matrix elasticity. Non-crosslinked scaffolds were modified with 100 µg/ml of either non-crosslinked or AGE-crosslinked collagen fibrils (**Extended Data Fig. 9a**). AFM measurements showed that pPFBs adhered significantly more strongly to AGE-crosslinked fibrils compared to non-crosslinked controls (**Extended Data Fig. 9b**). This enhanced adhesion led to increased fibroblast activation, demonstrated by higher expression of activation markers and adhesion molecules at both gene and protein levels (**Extended Data Fig. 9c-d**).

We next studied mechanical effects of matrix stiffness independent of crosslinking by comparing non-crosslinked scaffolds before and after 5% strain dynamic stretching (**Extended Data Fig. 9e**). This mechanical conditioning increased matrix stiffness without changing crosslinking levels. The stiffer substrates led to enhanced cell adhesion and activation markers, but surprisingly decreased RAGE expression (**Extended Data Fig. 9f-g**). These experiments revealed that while both AGE crosslinking and matrix stiffness promote fibroblast activation, they affect RAGE expression differently. This dual regulation creates a complex feedback loop in pulmonary fibrosis: increased AGE crosslinking and matrix stiffening reinforce each other, driving progressive fibroblast activation.

These findings suggest a counterintuitive mechanism in disease progression. Although stiffer matrices decrease RAGE expression, they enhance cell adhesion, which increases the probability of AGE-RAGE binding and promotes fibroblast activation (**Fig. 4**). This pathological state contrasts sharply with healthy lung conditions, where low AGE crosslinking, soft matrix, and high respiratory amplitude maintain high RAGE expression but low cell adhesion, keeping fibroblasts quiescent through reduced AGE-RAGE interactions. These findings reveal that biological materials can be controlled through either mechanical or chemical inputs, providing multiple pathways for therapeutic materials design.

### Restricted mechanical environment synergizes with pathological crosslinking to alter material-cell interactions

The interaction between mechanical loading conditions and crosslinking chemistry represents a critical design parameter for controlling biological material-cell interactions. To investigate how AGE crosslinking and biomechanical signals regulate pPFBs under pathological conditions, we applied 5% dynamic stretching strain to AGE-crosslinked scaffolds, mimicking the restricted respiratory movement in severe pulmonary fibrosis (**Extended Data Fig. 10a**). Pulmonary fibroblasts cultured under these pathological conditions (AGE scaffolds with 5% stretch) showed increased activation and adhesion compared to physiological conditions (NC scaffolds with 20% stretch), along with reduced RAGE expression, matching our *in vivo* observations (**Extended Data Fig. 10b**).

We found that this pathological condition activated the AGE-RAGE-MAPK signaling cascade. AGE binding to RAGE triggered mitogen-activated protein kinases (MAPKs), leading to activation and nuclear translocation of transcription factors including NF-κB. This transcriptional program upregulated target genes such as *Acta2*, *Col1a1*, *Mmp9*, *Ager*, and *IL6*^44,45^. Immunostaining revealed high nuclear levels of p-ERK1/2 and NF-κB in fibroblasts cultured on AGE-crosslinked scaffolds under 5% stretch, while cells under physiological conditions (NC scaffolds, 20% stretch) showed reduced expression and nuclear localization (**Extended Data Fig. 10b**). These results implicate MAPK signaling in stretch-modulated AGE-RAGE activation of pulmonary fibroblasts.

To comprehensively assess how well our *in vitro* model recapitulated pulmonary fibrosis, we performed RNA-Seq analysis on fibroblasts cultured under physiological and pathological conditions. Pathway enrichment analysis and GSEA showed that cells on the pathological matrix (AS-5%) displayed enriched expression of genes associated with focal adhesion, ECM-receptor interaction, MAPK signaling, and AGE-RAGE signaling pathways, matching the *in vivo* disease profile (**Extended Data Fig. 11**). We compared gene expression changes between *in vitro* conditions (NS-20% versus AS-5%) and *in vivo* fibrosis progression (Healthy versus Fibrosis), focusing on focal adhesion, ECM-receptor interaction, and AGE-RAGE signaling pathways (**Extended Data Fig. 10c**). qPCR validation (**Extended Data Fig. 10d**) confirmed that pathological conditions increased expression of fibroblast activation genes (*Acta2*, *Fn1*), cell adhesion genes (*Itga1*, *Itga11*), and MAPK pathway genes (*Mapk3*), while decreasing *Ager* expression.

Together, these results reveal how increased AGE-crosslinked ECM and restricted mechanical movement synergistically promote fibrosis through enhanced AGE-RAGE binding and MAPK-NF-κB pathway activation (**Extended Data Fig. 10e**). Importantly, physiological stretching (20% strain) can counteract these effects by reducing both AGE crosslinking and matrix elasticity, but only in porous scaffold matrices. This matrix-specific effect likely stems from differences in mechanotransduction between porous scaffolds and fibrous hydrogels. These results demonstrate that mechanical environment and crosslinking chemistry work synergistically to control cellular responses, establishing design principles for programmable biological materials.

### Chemical crosslinking inhibition reverses pathological material properties *in vivo*

Having demonstrated that AGE crosslinking can be controlled in engineered materials systems, we investigated whether chemical crosslinking inhibition could restore normal material properties in living tissues *in vivo*. We administered rosmarinic acid (RA) to mice with bleomycin-induced pulmonary fibrosis at low (LD), middle (MD), and high (HD) doses (**Fig. 5a**). Bleomycin was administered using a standard intratracheal pulmonary fibrosis model, but RA in saline was delivered daily to the stomach. RA treatment significantly improved survival rates and prevented weight loss in a dose-dependent manner (**Fig. 5b-c**).

**Fig. 5.**
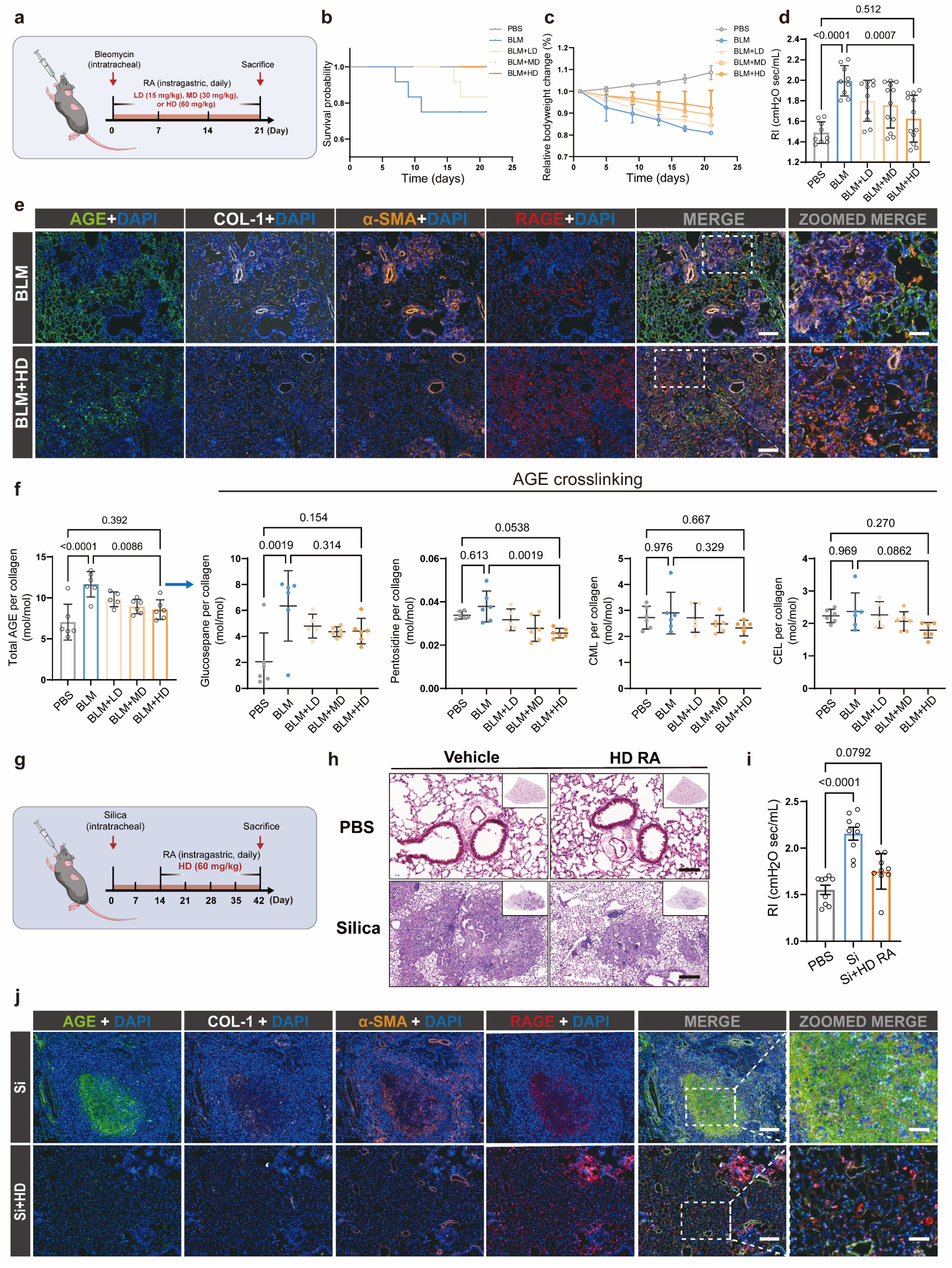
Pharmacological inhibition of AGE crosslinking in ECM reverses established pulmonary fibrosis in multiple disease models. (**a**) Treatment protocol for bleomycin (BLM)-induced pulmonary fibrosis showing BLM administration at day 0, followed by treatment with low (LD), medium (MD), or high dose (HD) rosmarinic acid (RA) (*n* = 9 mice per group). (**b**) Survival curves demonstrating dose-dependent improvement with RA treatment, indicating significant therapeutic benefit. (**c**) Body weight changes demonstrating protective effects of medium (MD) and high dose (HD) RA against BLM-induced weight loss. (**d**) Lung function (resistance index, RI) showing restoration toward normal levels with HD RA and demonstrating functional improvement; see also Figure. S12A. (**e**) Immunofluorescence of lung sections comparing vehicle versus HD RA treatment, showing normalized levels of AGE crosslinking (green), COL-1 (white), α-SMA (orange), and RAGE (red), demonstrating reversal of fibrotic phenotype (nuclei: DAPI, blue, scale bars: 200 μm, insets 50 μm). (**f**) Quantitative analysis of AGE crosslinking in decellularized lung ECM confirming significant reduction with RA treatment, supporting the mechanistic basis of therapeutic effect. Results expressed as molar ratio of crosslinks to tropocollagen molecules (*n* ≥ 5 ECM samples from ≥3 mice per group). (**g**) Treatment protocol for silica-induced pulmonary fibrosis model, testing RA in a different disease etiology. (**h**) H&E-stained lung sections from silicosis mice showing marked reduction in fibrotic changes with HD RA treatment. Scale bars: 100 μm. (**i**) Lung function improvements following HD RA treatment in the silicosis model, demonstrating generalizability across different fibrosis etiologies; see also Figure. S14D. (**j**) Immunofluorescence of silicosis model lung sections showing AGE, COL-1, α-SMA, and RAGE, confirming that the same mechanobiological pathway is targetable in multiple forms of pulmonary fibrosis (nuclei: DAPI, blue; scale bars: 200 μm, insets 50 μm). Statistical analysis performed using one-way ANOVA with Tukey’s post-hoc test, exact *p* values labeled. Results presented as mean ± S.D.

High-dose RA treatment dramatically improved lung function in treated mice (**Fig. 5d, Extended Data Fig. 12a**), with decreased inspiratory resistance and increased inspiratory capacity, dynamic lung compliance, and static lung compliance. We observed a transient elevation in blood glucose at day 21 of treatment that normalized by day 28 (**Extended Data Fig. 12b**), possibly suggesting that RA may act as a glucose breaker, releasing previously crosslinked glucose and reducing AGE formation. Histological analysis confirmed restoration of normal pulmonary architecture (**Extended Data Fig. 12c-e**). Immunofluorescence staining revealed reduced expression of fibrosis markers (COL-1, α-SMA) and AGE crosslinking, while RAGE expression increased toward normal levels (**Fig. 5e**). Quantitative AQMC analysis confirmed that RA treatment significantly reduced AGE crosslinking in lung ECM (**Fig. 5f**), but did not significantly reduce cross-linking by TGM and LOX **(Extended Data Fig. 12f-g)**. RA treatment inhibited alleviated several additional markers of pulmonary fibrosis **(Extended Data Fig. 12h-m)** and inflammatory responses (**Extended Data Fig. 13**) following BLM-induced pulmonary fibrosis.

These findings demonstrate that targeting AGE crosslinked ECM can break the mechanobiological feedback loop driving pulmonary fibrosis, leading to improvement in both tissue structure and function. This successful chemical intervention validates the therapeutic potential of targeting crosslinking networks and demonstrates that pathological material properties can be reversed through materials-based approaches.

### Crosslinking network control represents a generalizable materials intervention strategy

To test whether crosslinking network control represents a generalizable materials intervention strategy, we evaluated the approach in a different model system with distinct initial triggers. Silica-induced pulmonary fibrosis presents a mechanobiological puzzle: while initial inflammation and injury are triggered by silica particles^46^, the progressive nature of the disease persists even after particle clearance^47^, when residual silica levels fall well below the percolation threshold necessary for direct mechanical effects^48^. This persistence suggests that, similar to bleomycin-induced fibrosis, a self-sustaining cycle of cell-ECM feedback may be involved in disease progression independently of the initiating stimulus.

To test this hypothesis and evaluate the generality of AGE-mediated crosslinking in fibrosis, we administered rosmarinic acid to mice with established silica-induced pulmonary fibrosis which continued to worsen beyond six weeks following a single silica exposure^49,50^ (**Fig. 5g**). Histological analysis revealed marked reduction in fibrotic changes (**Fig. 5h, Extended Data Fig. 14a-c**). High-dose RA treatment significantly improved lung function parameters and normalized blood glucose levels (**Fig. 5i, Extended Data Fig. 14d-e**), while immunofluorescence showed decreased expression of AGE crosslinking, COL-1, and α-SMA, accompanied by recovery of RAGE expression (**Fig. 5j**). Western blot analysis confirmed reduced fibronectin (FN-1) expression, indicating decreased ECM production (**Extended Data Fig. 14i-j**). Comprehensive molecular analysis demonstrated that RA treatment reduced both fibrotic markers (**Extended Data Fig. 14f-h**) and inflammatory mediators (**Extended Data Fig. 15**).

The effectiveness of AGE inhibition in both bleomycin- and silica-induced fibrosis suggests that AGE- mediated ECM crosslinking represents a common pathway in the mechanobiological feedback loop driving disease progression. This finding is particularly significant for silica-induced fibrosis, where it helps explain how the disease can progress even after the initial mechanical stimulus is removed, and points to AGE inhibition as a broadly applicable therapeutic strategy for pulmonary fibrosis of diverse etiologies. The success across multiple disease models establishes crosslinking network control as a broadly applicable materials intervention strategy, independent of the initial tissue damage mechanism.

### Mechanotherapy achieves unprecedented dynamic control of biological material properties

The discovery that mechanical forces can disrupt pathological crosslinking inspired the development of mechanotherapy - a materials-based treatment that uses controlled mechanical forces to dynamically modify tissue properties. To determine the optimal dosage for mechanotherapy, we first applied a small animal ventilator to deliver tidal volumes of 0.3 ml, 0.6 ml, or 0.9 ml to mice with established pulmonary fibrosis. Histological analysis (**Extended Data Fig. 16a**) and pulmonary function tests (**Extended Data Fig. 16b**) revealed that low tidal volume (0.3 ml) significantly reduced fibrosis and improved lung function, while higher volumes exacerbated lung injury, consistent with known risks of ventilator-induced injury at high volumes^51^.

Based on these results, we evaluated this mechanotherapy in a controlled trial, comparing ventilation alone (0.3 ml tidal volume), RA alone, and combined treatment (**Fig. 6a**). The combined therapy showed the strongest therapeutic effect, followed by RA and ventilation monotherapies, as measured by survival (**Fig. 6b**) and prevention of disease-associated weight loss (**Fig. 6c**). Histological analysis revealed that mechanical ventilation reduced ECM deposition and tissue injury, with maximal improvement in the combination therapy group (**Extended Data Fig. 17a**). This finding was confirmed by hydroxyproline quantification (**Fig. 6d**), pulmonary function testing (**Fig. 6e, Extended Data Fig. 17b**), and molecular analysis of fibrosis markers (**Extended Data Fig. 17c-d**). Mechanical testing of mouse lung tissue revealed that mechanical ventilation reduced the mechanical strength of lung tissue, rendering it more susceptible to strain (**Fig. 6f**). Immunofluorescence analysis further demonstrated that ventilation-mediated attenuation of pulmonary fibrosis correlates with reduced AGE-mediated ECM crosslinking, suggesting a mechanochemical basis for improved tissue mechanics (**Fig. 6g**). This successful demonstration of mechanotherapy establishes a new paradigm for materials-based medicine, where mechanical forces serve as precision tools for real-time control of biological material properties in living systems.

**Fig. 6.**
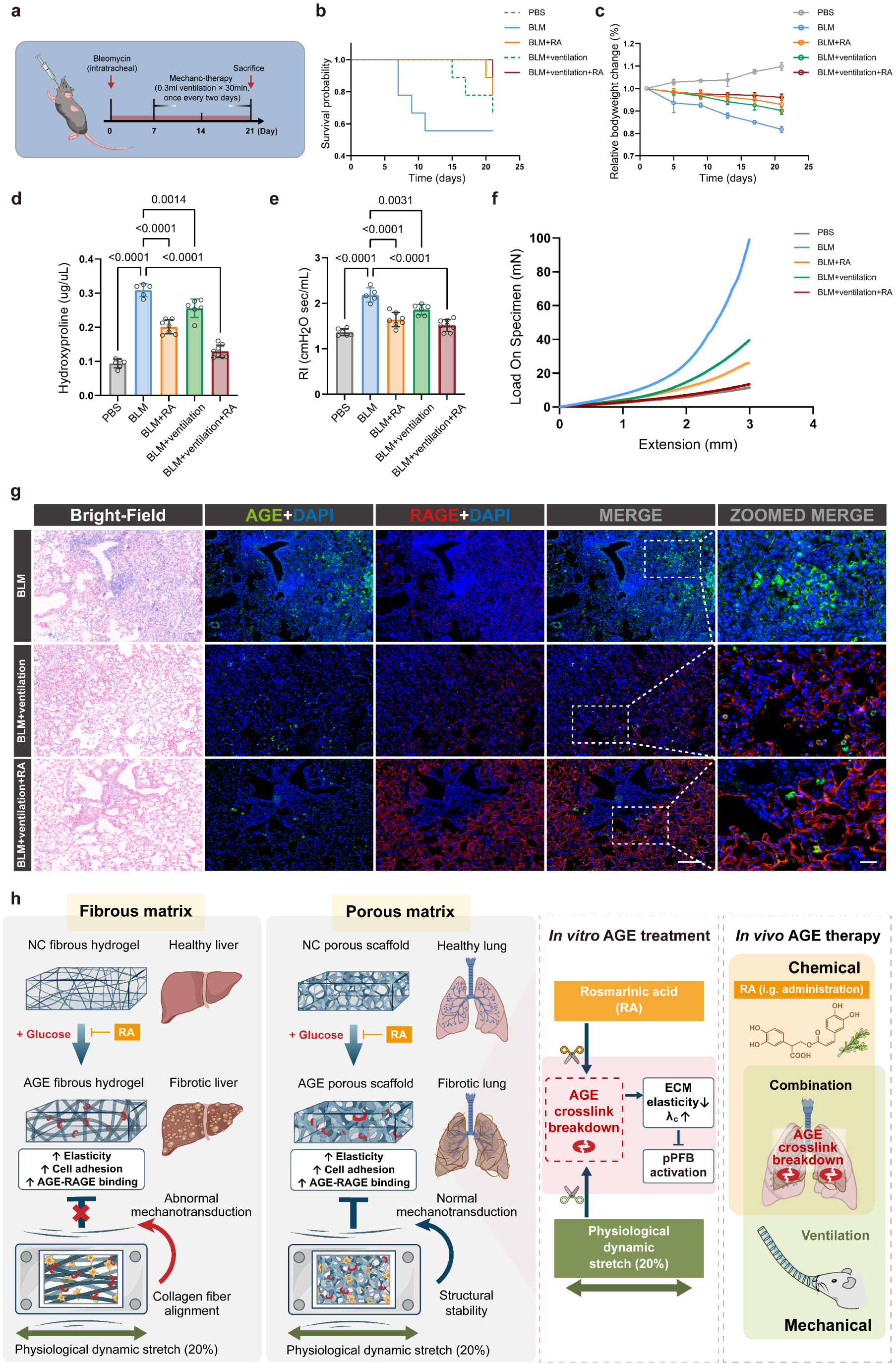
Ventilation-based mechanotherapy reverses established pulmonary fibrosis and synergizes with pharmacological AGE inhibition. (**a**) Experimental design showing bleomycin (BLM) administration at day 0, followed by treatment initiation at day 7 with either low-volume mechanical ventilation (0.3 mL tidal volume, 30 minutes every 48 hours), high-dose rosmarinic acid (RA), or combination therapy (ventilation plus RA) (*n* ≥ 5 biologically independent mice per group). (**b**) Survival analysis demonstrating enhanced survival with both single treatments and maximal benefit from combination therapy, supporting the concept of mechanochemical treatment. (**c**) Body weight trajectories showing protection against BLM-induced weight loss, with combination therapy providing the greatest benefit. (**d**) Hydroxyproline content analysis demonstrating significant reduction in collagen accumulation with all treatments, most pronounced with combination therapy (*n* ≥ 5 mice per group). (**e**) Pulmonary function measurements showing restoration of lung mechanics toward normal levels, confirming functional recovery. (**f**) Force-displacement curves demonstrating how treatments alter the mechanical properties of lung tissue, with mechanical ventilation reducing tissue stiffness. (**g**) Immunofluorescence of lung sections showing reduction in AGE crosslinking (green) and recovery of RAGE expression (red) with mechanical ventilation and combined therapy (nuclei: DAPI, blue; scale bars: 200 μm, insets 50 μm). (**h**) Mechanistic model illustrating how increased AGE crosslinking and restricted mechanical movement promote fibrosis through enhanced AGE-RAGE binding, and how targeted interventions can disrupt this pathological cycle. Statistical analysis performed using one-way ANOVA with Tukey’s post-hoc test, exact *p* values labeled. Data presented as mean ± S.D.

## Discussion

Our findings establish a new paradigm for controlling biological material properties through mechanical forces, revealing fundamental structure-property relationships that enable tissue-specific therapeutic interventions. By demonstrating that mechanical forces can selectively disrupt pathological crosslinking networks, we introduce the concept of mechanically-controlled biological materials, a breakthrough that bridges fundamental materials science with transformative medical applications.

The discovery that identical mechanical inputs produce opposite effects depending on material architecture represents a critical advance in understanding biological materials design. Our biomimetic systems reveal that porous scaffolds respond favorably to mechanical disruption while fibrous matrices remain largely unaffected, establishing architecture as a primary determinant of mechanical responsiveness. This architecture-dependent behavior explains why traditional materials interventions show varying efficacy across different tissue types and provides design principles for engineering responsive biological materials.

The mechanobiological feedback loop we identified shows that pathological crosslinking reduces a key aspect of nonlinear tissue compliance and the mechanical stretch amplitude, leading to enhanced cellular activation and further crosslinking. This represents a self-reinforcing materials failure mode previously unrecognized in biological systems. Unlike synthetic materials where crosslinking typically stabilizes mechanical properties, biological tissues exist in dynamic mechanical environments where crosslinking density can be modulated post-formation. This insight opens possibilities for developing materials that can be mechanically “reprogrammed” in real time.

Our computational modeling reveals that crosslink density directly controls critical mechanical transitions in fibrous networks, with implications extending beyond biological systems to synthetic materials design. The relationship between crosslink density and strain-stiffening behavior provides predictive frameworks for designing materials with tunable mechanical responses. The discovery that mechanical forces can selectively target specific crosslink types (AGE but not enzymatic crosslinks) suggests opportunities for developing materials with hierarchical mechanical control systems.

The successful demonstration of controlled mechanical forces as therapeutic tools (“mechanotherapy”) establishes the mechanical environment as a designable parameter in materials-based medicine. Unlike static interventions that rely on passive material properties, mechanotherapy enables dynamic, real-time modification of tissue mechanics. This approach is particularly powerful when combined with chemical interventions, creating “mechanochemical” control systems that achieve unprecedented therapeutic outcomes.

The tissue-specific nature of our findings has broader implications for materials design. The lung’s unique combination of porous architecture and cyclic mechanical loading creates conditions not found in other biological systems, enabling mechanical interventions impossible elsewhere. This observation suggests that understanding organ-specific mechanical environments could inform the design of next-generation biomaterials and tissue engineering scaffolds optimized for specific anatomical locations.

Beyond the immediate therapeutic applications, our work establishes principles for programmable biological materials. The demonstration that mechanical forces can serve as precision control inputs for modifying crosslinking networks opens new avenues for developing smart materials that respond to mechanical stimuli. This could enable materials that self-heal under mechanical stress, or scaffolds that adapt their properties in response to tissue loading patterns.

The clinical success of our mechanochemical approach across multiple disease models validates the generalizability of crosslinking network control as a materials intervention strategy. This suggests that the principles we’ve identified could be applied to a broad range of conditions where pathological crosslinking contributes to tissue dysfunction, potentially revolutionizing treatment approaches for age-related diseases and fibrotic conditions.

Looking forward, our findings point toward new “mechanically-controlled therapeutic materials” where physical forces are used as precision tools for modifying biological material properties. The integration of mechanical and chemical control mechanisms provides a powerful platform for developing next-generation materials that can be dynamically programmed for specific applications. This work establishes the foundation for materials systems that combine the precision of mechanical control with the specificity of chemical targeting, opening unprecedented possibilities for both therapeutic applications and fundamental materials research.

## Supporting information

Supplementary information including Sup Note, Sup Figures 1-2, Sup Tables 1-4

## Methods

### Human lung samples

Fresh lung tissue samples from patients (as shown in Supplementary Table 1) were obtained from China-Japan Friendship Hospital (Beijing, China). All subjects agreed to participate in the study after receiving detailed information, and all research protocols were consistent with the guidelines outlined in the 2013 Declaration of Helsinki, and were approved by the Human Studies Committee of Peking Union Medical College.

### Human liver samples

Fresh liver tissues from patients with established liver cirrhosis were provided by the Department of Hepatology at Tsinghua Changgung Hospital, as approved by the Institutional Review Board of Tsinghua University. Informed consent was obtained from participants. Paraffin-embedded liver tissues from human organ donors were purchased from OriGene Technologies.

### Animal models of lung and liver fibrosis

C57BL/6J mice weighing between 25-30 g and aged 8 weeks were accommodated in sterile cages under standard conditions of 24-26°C temperature, 60-70% humidity, and a 12-hour light/12-hour dark cycle. These mice were housed in a pathogen-free controlled facility with unrestricted access to food and water. All animal study protocols were reviewed and approved by the Institutional Animal Care and Use Committee at Peking Union Medical College. For the bleomycin (BLM)-induced pulmonary fibrosis model, 8-week-old male C57BL/6 mice were anesthetized with tribromoethanol (1.2 mL/100 g, intraperitoneal) and received a single intratracheal instillation of BLM (1.5-2 U/kg body weight). For the silicosis model, mice received a single intratracheal instillation of silica (300 mg/kg body weight). Sham-operated control groups received an equal volume of PBS intratracheally. The BLM-induced chronic lung fibrosis models were evaluated at 3 weeks post-instillation, while the silica-induced silicosis models were assessed at 6 weeks post-exposure. For the CCl_4_-induced late-stage liver fibrosis model, 8-week-old male C57BL/6 mice received an intraperitoneal injection of CCl_4_ (2.5 ml/kg body weight; 1:4 diluted with olive oil) twice weekly for 15 weeks.

### Lung function tests

Mice were induced anesthesia with pentobarbital (intraperitoneal, 0.2 ml/100 g, Sigma-Aldrich) and subsequently tracheotomized. A modified needle was securely fixed to the trachea for cannulation. The mice were then connected to a FlexiVent respirator (SCIREQ, FlexiVent, Montreal, Canada). Standard spirometry techniques were employed to measure several pulmonary function parameters, including inspiratory capacity (IC), airway resistance (RI), dynamic compliance (Cdyn), and chord compliance (Cchord). Each measurement was based on the mean value derived from three individual replicates.

### Histological staining

For immunofluorescence staining, frozen tissue sections were fixed with 4% paraformaldehyde, followed by permeabilization and blocking for 1 h. Sections were incubated with primary antibodies at 4 °C overnight, then with fluorescent secondary antibodies at 37 °C for 1 h. Nuclei were stained with Hoechst-33432 (Beyotime, 1:2000). For immunohistochemistry, tissue sections were dried at 60 °C for 30 min, followed by deparaffinization and rehydration. Antigen retrieval was performed in sodium citrate buffer using an autoclave for 20 min. Endogenous peroxidase activity was inhibited with 3% hydrogen peroxide. Sections were permeabilized and blocked for 1 h, followed by primary antibody incubation at 4 °C overnight. Biotinylated secondary antibodies were applied at room temperature for 1 h, and visualization was developed using 3,3′-diaminobenzidine. The following primary antibodies were used: anti-RAGE (Abcam, ab216329), anti-AGE (Abcam, ab23722), anti-COL-1 (Abcam, ab6308), and anti-α-SMA (Abcam, E184). Sirius red staining was performed according to the manufacturer’s protocol (Huayueyang Bio-Technology, GH6044s). Hematoxylin and eosin (HE) staining, following the Szapiel method^52^, was conducted to evaluate the level of lung inflammation. Masson’s trichrome staining was performed to assess the extent of fibrosis.^53^ Histological images were captured using a 3DHISTECH “Pannoramic” SCAN system (3DHISTECH, Budapest, Hungary).

### Lung tissue decellularization

To obtain decellularized ECM from mouse lungs, an *in-situ* perfusion protocol was followed (**Extended Data Fig. 1b**). Briefly, mice with established pulmonary fibrosis were anesthetized with 2.5% avertin, and the inferior vena cava was cannulated with a polyethylene tube connected to a peristaltic pump. The flow rate was set to 3 mL/min, and the left auricle was severed to allow buffer outflow. The perfusion sequence consisted of 1.9% EGTA solution for 20 min to remove blood, followed by sequential perfusion with 0.01%, 0.05%, and 0.1% sodium dodecyl sulfate (SDS) for 12 h each, continuing until the lung tissue became translucent and the perfusate tested negative for protein and DNA. The tissue was then perfused with deionized water for 12 h to thoroughly remove buffer residues. The decellularized ECM was harvested and preserved in a vacuum drying chamber after CO_2_ critical point drying. Complete decellularization was verified by hematoxylin and eosin staining to confirm the absence of cellular components.

For human pulmonary fibrosis samples, lung tissue specimens were first processed by thorough washing and vertexing in 1% SDS buffer for 10 min, followed by sonication for 10 min. Tissue was then centrifuged at 12,000 rpm for 15 min at 4 °C. The pellet was collected and this process was repeated until the supernatant became clear and white, fibrous, decellularized ECM was obtained. Human lung decellularized ECM was washed and preserved using the method described for mouse tissue.

### Sample hydrolysis

Dried samples were weighed and placed in glass hydrolysis vessels for treatment with 6 mol/L HCl. The vessels were sealed and incubated in a drying oven at 110 °C for 48 h. Samples were then dried using a RapidVap Vacuum Evaporation System (LABCONCO, USA). For crosslink quantification by LC-MS/MS, samples were dissolved in 80% methanol. For hydroxyproline content quantification, samples were dissolved in deionized water, and the pH was adjusted to 7.4 using 10 mol/L NaOH.

### Synthesis of glucosepane

Glucosepane was synthesized according to a modification of a protocol described previously^45^. Briefly, lysine (0.8771 g), arginine (0.6968 g), and D-glucose (0.3603 g) were dissolved in 10 mL of deionized water. The reaction mixture was filtered through a 0.22-μm filter and incubated at 50°C for 21 days. The glucosepane fractions were separated by preparative high-performance liquid chromatography (HPLC) using a Waters MS-directed Auto-Purification system (USA) equipped with an automatic collector/preparative liquid/mass spectrometer (models 2545/2767/QDa, Waters, USA). The separation was performed on an Agilent Eclipse XDB-C18 column (4.6 × 250 mm, 5 μm; Agilent, USA). The mobile phase consisted of water containing 10 mM ammonium formate (eluent A) and methanol (eluent B). The flow rate was 1 mL/min, with isocratic conditions of 1% solvent B maintained for 18 minutes to ensure retention time stability. Fractions with retention times (RTs) of 2.1-3.0 min (electrospray ionization detection, mass-to-charge ratio (m/z) 429 for the protonated molecule [M+H]+) were collected and lyophilized. The purity of the obtained product was determined by liquid chromatography-mass spectrometry (LC-MS, column B, gradient B). The collected fraction contained three byproducts with RTs of 1.8, 9.35, and 19.96 min. Glucosepane had an RT of 19.96 min with 57.5% purity. The purity analysis was performed using an UHPLC-Q-Orbitrap LC-MS system consisting of an Ultimate 3000 Ultra High Performance Liquid Chromatography unit (UHPLC, Dionex, USA) and Q-Exactive Plus High-Resolution Mass Spectrometer (Thermo Fisher Scientific, USA) in positive ionization mode. The LC separation was performed on an ACQUITY UPLC BEH Amide column (2.1 × 100 mm, 1.7 μm; Waters, USA). The mobile phase consisted of acetonitrile containing 0.1% formic acid (eluent A) and water containing 10 mM ammonium acetate and 0.1% formic acid (eluent B). Gradient elution was performed at a flow rate of 0.3 mL/min with a column temperature of 25°C using the following program: 0% B for 6 minutes, 25% B for 10 minutes, 50% B for 10 minutes, and 0% B for 4 minutes. UV detection was performed at 254 nm.

### Isolation of crosslinking products

The dried sample was placed in a tube for enzymatic digestion. The following proteases were added stepwise every 24 h: (1) 2 mg/mL collagenase I (GIBCO, 17100-017) in 1 mL TRIS-HCl buffer (pH, 8); (2) 0.8 units of pronase (Roche, 11459643001); (3) 0.8 units of pronase and 0.5 mg papain (Sigma, P4762); (4) 0.8 units of aminopeptidase M (Shanghai Yuanye, S25360), 1 unit prolidase (Sigma, P6675) and 10 mM Mg^2+^/Mn^2+^ to activate the enzyme; (5) 0.8 units of aminopeptidase M; (6) 0.65 units of carboxypeptidase (Sigma, SAE0046). Thymol (1 mg; Macklin, T818893) was added to inhibit bacterial growth. Samples were incubated in a shaking incubator at 37 °C. The enzyme was inactivated by vortexing and sonication at 100 °C for 10 min after each step. Quality control was performed to evaluate the efficiency of enzymatic digestion by comparing the hydroxyproline content in samples treated by enzymatic digestion versus acid hydrolysis using HPLC-tandem mass spectrometry (HPLC-MS/MS). After the final step, enzyme residues were precipitated with cold methanol for 1 hour and removed after centrifugation at 13,000 *G* (*G* = 9.81 m/s^2^) for 15 min. Crosslinks in the supernatants were collected and dried using a RapidVap Vacuum Evaporation System (LABCONCO, USA). Samples were dissolved in 80 μL of 80% methanol for detection of γ-glu-ε-lys, glucosepane, and pentosidine by HPLC-MS/MS. Samples were further hydrolyzed with 6 mol/L HCl for an additional 48 h as described in the sample hydrolysis section for the detection of pyridinoline (PYD), carboxymethyl-lysine (CML), and carboxyethyl-lysine (CEL).

### High performance liquid chromatography-tandem mass spectrometry/mass spectrometry (HPLC-MS/MS)

HPLC was performed on a Jasso (Japan) system consisting of a five-way online degasser (ExionLC Degasser-5036645, Sciex, USA), an ultra-efficient binary pump (ExionLC AD Pump-5036653, Sciex, USA) with a solvent mixer (20 μL Micro Mixer-45054489), a column thermostat (ExionLC AD Oven-5036656, Sciex, USA), and an autosampler (ExionLC AD Autosampler-5036654). Mass detection was performed on an AB Sciex API 5500 Qtrap LC/MS/MS system (Sciex, USA) triple quadrupole mass spectrometer equipped with a turbo ion spray source, using positive electrospray ionization ion mode. The MS parameters were set as follows: ESI ion source temperature of 500 °C, air curtain pressure of 30 psi (207 kPa), collision-activated dissociation gas set to medium, ion spray voltage of 5500 V, and ion gas 1 and 2 pressures of 50 psi (345 kPa). Chromatographic separation was performed on a Luna NH_2_ analytical column (2 × 100 mm, 3 μm, Phenomenex, USA) equipped with Exion LC AD system (Sciex, USA) infinite binary pump (ExionLc AD Pump-5036653, Sciex, USA). The mobile phase consisted of eluent A (acetonitrile containing 20 mM ammonium acetate and 0.2% formic acid) and eluent B (water containing 20 mM ammonium acetate and 0.2% formic acid). Gradient elution was performed at a flow rate of 0.4 mL/min with a column temperature of 25 °C using the following program: 10% B for 8 min, 80% B for 2 min, and 10% B for 4 min. Samples were maintained at 10 °C, and the injection volume was 5 μL. Mass spectrometry detection was performed in multiple reaction monitoring mode, using collision-induced dissociation (CID) of protonated molecules, with compound-specific declustering potential (DP) and fragment-specific collision energy (CE). The MS_2_ fragmentation spectra of the crosslinking products used for detection are listed in Supplementary Table 2.

### Determination of crosslinking degree

MS data were analyzed using AB Sciex Analyst 1.7.1 software (Sciex, USA). Quantification was performed using external standard methods. Standard crosslinking compound mixtures of increasing concentrations were analyzed by HPLC-MS/MS using the methods described in the HPLC-MS/MS section. Standard curves of detector response versus concentration were used to determine the concentration of corresponding crosslinking products in each sample. Background signals from the isolation and preparation processes were subtracted before determining the final content of crosslinking products. The crosslinking degree mediated by a specific crosslinking mechanism in a sample was defined as the molar ratio (mol/mol) of crosslinking products to tropocollagen molecules detected in the sample, as illustrated in **Extended Data Fig. 2a**. The degrees of lysyl oxidase (LOX) and transglutaminase (TGM) crosslinking were determined by measuring pyridinoline and γ-glutamyl-ε-lysine, respectively. The total degree of advanced glycation end-product (AGE) crosslinking was determined by the molar sum of carboxymethyl-lysine (CML), carboxyethyl-lysine (CEL), glucosepane, and pentosidine. The molar quantity of triple-helix tropocollagen molecules was determined by hydroxyproline content, assuming 300 hydroxyproline residues per tropocollagen molecule^54^. Authentic standards used for determination include pyridinoline (Toronto Research Chemicals, H954036), γ-Glutamyl-ε-Lysine (γ-glu-ε-lys, Sigma, G5136), N^ε^-(1-carboxymethyl)-L-lysine (CML, Cayman Chemical, 16483), N^ε^-(1-carboxyethyl)-L-lysine (CEL, Cayman Chemical, 25333), pentosidine (PYD, Cayman Chemical, 10010254) and hydroxyproline (HYP, Solarbio, SH8250).

### Construction of crosslinked collagen matrices

The pre-mixed solution was prepared by combining calculated volumes of reagents according to the following formulas: PBS (10×) volume was calculated as final volume/10; collagen I volume as (final volume × final collagen I concentration)/stock concentration; 1M NaOH as 0.023 × collagen I volume; transglutaminase (TGM) volume as (final volume × final TGM concentration)/stock concentration; and water volume as final volume minus the sum of all other reagent volumes. For non-crosslinked collagen matrix preparation, collagen I solution (rat tail Collagen I, 354236, Corning, USA) was neutralized to pH 7.4 using 1M NaOH in the presence of 10X PBS. The final concentrations were adjusted to 2 mg/mL collagen and 10 mg/mL TGM (BioBomei, BC5582) in the premix. The solution was gently mixed with a pipette and immediately transferred (500 μL per well) into a 12-well culture plate. All steps were performed on ice. Gelation was allowed to proceed at 37°C for 5 hours in a humidified atmosphere, after which PBS buffer was added gently to cover the collagen matrix surface for subsequent assays. For AGE-crosslinked collagen matrix preparation, glucose solution (Sigma, G8769) was added to the neutralized collagen I mixture to achieve a final concentration of 250 mM. The covering buffer also contained 250 mM glucose and was replaced every three days for 4 weeks to allow AGE crosslinking reactions to occur. For AGE+RA collagen matrix preparation, the pre-mixed solution and covering buffer were supplemented with 100 μM rosmarinic acid (RA) (Shanghai Yuanye, S31633) during AGE-crosslinked matrix preparation. RA solutions were prepared immediately before use, and the covering buffer was replaced every three days to maintain inhibitory effects. All crosslinking degrees were verified quantitatively using the AQMC method as illustrated in **Extended Data Fig. 2a**. After hydrogel matrix preparation, samples were thoroughly washed with endotoxin-free water, excess moisture was removed, and matrices were placed in a −80°C freezer overnight. Matrices were then freeze-dried for 12 h to obtain the final scaffold matrices.

### Micro-nano tensile testing

Collagen scaffold matrices were prepared as dog-bone-shaped specimens with a gauge width of 5 mm for standard tensile testing. Specimen thickness, measured using a caliper, averaged approximately 2 mm. Specimens were aligned parallel to the loading direction of a tensile testing machine (Agilent Technologies T150 UTM, USA). Specimens were loaded at a velocity of 0.1 mm/s (approximate strain rate, 0.005/s). Stress-strain data were collected after a 10 μN tension trigger was tripped. Four scaffold matrix specimens were tested for each condition.

### SEM sample preparation

Lung decellularized extracellular matrix (ECM) and *in vitro* collagen matrix samples were washed thoroughly three times with distilled water for 4 h each. Samples were dehydrated through a graded ethanol series (50%, 70%, 80%, 90%, 100%) for 10 min at each concentration, followed by overnight incubation in 100% ethanol. CO_2_ critical point drying was performed using a Leica EM CPD300 system. The samples were sputter-coated with 10 nm platinum/palladium (80/20) using a Leica EM ACE600 system. Images were acquired using a Zeiss Gemini SEM 500 scanning electron microscope.

### Mouse primary pulmonary fibroblast (pPFB) isolation

Mouse pPFBs were prepared from healthy male mice (8-12 weeks old C57BL/6) as described previously^55^. Briefly, lungs were removed from the thoracic cavity, placed into a 100 mm tissue culture petri dish, and minced into very small pieces using a razor blade or surgical scissors. The minced tissue was transferred into a T25 cell culture flask, and 3 mL of digestion medium (DMEM containing 0.1% collagenase I, 0.1% collagenase IV, 0.01% deoxyribonuclease, and 0.01% hyaluronidase) was added. The tissue was incubated at 37 °C for 6-8 h. After incubation, the cell suspension was filtered through a 40 μm cell strainer to remove debris and centrifuged at 1,000 *G* (*G* = 9.81 m/s^2^) for 5 min at 24 °C. The supernatant was carefully decanted and discarded. The cell pellet was washed with 1 mL of Hanks’ Balanced Salt Solution (HBSS) and centrifuged again at 1,000 *G* for 5 min at 24 °C. After discarding the supernatant, cells were resuspended in DMEM supplemented with 10% FBS and 1% penicillin-streptomycin solution (Gibco) and distributed into 100-mm petri dishes. Cells were incubated at 37 °C in 5% CO_2_. After 24 h, the medium was removed gently, and dishes were washed 2-3 times with 10 mL PBS to remove red blood cells, dead cells, and debris. Fresh fibroblast culture medium was then added to each dish.

### Growth of pPFBs on crosslinked collagen matrices

The prepared crosslinked collagen matrices were thoroughly washed with PBS to remove reagent residues. pPFBs were collected and suspended in DMEM supplemented with 10% FBS and 1% penicillin-streptomycin solution. Cell suspensions were applied to the collagen matrices at a concentration of 3 × 10^4^ cells/mL. The matrices were incubated for 24 h without disturbance to allow cell adhesion. After this period, the medium was discarded and carefully replaced with fresh medium. The cells were then cultured on the matrices for an additional 24 h. RNA and cells were isolated 24 h after induction as illustrated in **Fig. 4**, **Extended Data Fig. 7, 9-10, 19-20**.

### RNA isolation and real-time qPCR analysis

Total RNA was isolated using RNA extraction reagent (Vazyme, R401-01). For cells grown in collagen matrices, the matrices were first dehydrated, then treated with RNA extraction reagent and vortexed for 10 minutes. Complementary DNA (cDNA) was synthesized using a 5× Hiscript II qRT SuperMix Kit (V) (Vazyme, R222-01-AB). Real-time quantitative PCR (qPCR) was performed using 2× AceQ qPCR SYBR Green Master Mix (Vazyme, Q121-02-AA) in a CFX96 Real-Time PCR Detection System (Bio-Rad). The primers and their corresponding sequences used for real-time qPCR analysis are listed in Supplementary Table 3, with *bactin* serving as the housekeeping gene.

### RNA sequencing

RNA sequencing analysis was conducted on two sets of samples: (1) pPFBs isolated from healthy and fibrotic mouse lungs *in vivo* (*n* = 3 per condition), and (2) pPFBs cultured *in vitro* on non-crosslinked scaffolds with 20% strain (NS-20%) and AGE-crosslinked scaffolds with 5% strain (AS-5%) (*n* = 3 per condition). Total RNA was extracted using RNA extraction reagent (Vazyme, R401-01). RNA-seq libraries were prepared from 500-1000 ng total RNA using the VAHTS Universal V8 RNA-seq Library Prep Kit for Illumina (Vazyme, NR605). The protocol included mRNA enrichment using VAHTS mRNA Capture Beads (Vazyme, N401), RNA fragmentation, and first and second strand cDNA synthesis. The resulting double-stranded cDNA was PCR-amplified to generate the final libraries. Library quality control included concentration measurement using the Equalbit 1 × dsDNA HS Assay Kit (Vazyme, EQ121) and insert size analysis using NanoQC with Miseq (Illumina). Libraries were sequenced on a NovaSeq 6000 using an S4 flow cell (Illumina), generating 40-48 million paired-end reads per sample.

For bioinformatic analysis, sequence reads were mapped to the Mus musculus transcriptome (Ensembl release 90, GRCm38) using HISAT2 version 2.0.5. Gene expression was quantified as Fragments Per Kilobase Million mapped reads (FPKM) using HTSeq-count version 0.6.0. Differential expression analysis was performed using DESeq2 version 1.6.3, with significantly differentially expressed genes (DEGs) defined by false discovery rate (FDR) < 0.05 and |log_2_(Fold Change)| > 1.

Data visualization was performed in R version 4.1.2, with heatmaps generated using the pheatmap package version 1.0.12. To compare *in vitro* and *in vivo* pPFB responses, we analyzed genes showing significant differential expression across all conditions (NS-20% vs AS-5% and healthy vs fibrotic). Gene Ontology (GO) annotation and Kyoto Encyclopedia of Genes and Genomes (KEGG) pathway enrichment analyses were performed using clusterProfiler version 3.19. Gene Set Enrichment Analysis (GSEA) was used to identify functional pathways differentially regulated between healthy and fibrotic conditions, with significance defined as FDR < 0.05. Results focusing on cell adhesion and AGE-RAGE signaling pathways are presented in **Extended Data Fig. 4, 10-11**.

### Immunofluorescence staining of cell culture samples

Cells grown on collagen matrices were fixed with 4% paraformaldehyde in microtubule-stabilizing buffer (100 mM PIPES, 5 mM EGTA, 2 mM MgCl_2_, pH 6.8) for 30 min as described previously^56^. Samples were washed three times with PBS. Cells were permeabilized and blocked for 1 h with gentle shaking in PBS containing 0.3% Triton X-100 (Sigma) and 3% bovine serum albumin (BSA, Amresco). Primary antibodies, diluted in 1% BSA, were applied overnight at 4°C, followed by three PBS washes. The primary antibodies used were anti-α-SMA (Abcam, E184), anti-RAGE (Abcam, ab216329), anti-paxillin (Abcam, ab32115), anti-phospho-ERK1/2 (Abmart, T40072) and anti-NF-κB (Abmart, T55034F) Cells were then incubated with corresponding secondary antibodies for 1 h. Nuclei were stained with Hoechst 33342 (Beyotime, 1:2000) for 15 min. Filamentous actin (F-actin) was labeled using Acti-stain 488 phalloidin or Acti-stain 555 phalloidin (1:200; Cytoskeleton, USA).

### Confocal imaging and analysis

Confocal images were acquired using either an Olympus FV3000 or a Leica Stellaris8 confocal microscope. Images were captured using a 100× oil objective. For analysis of α-SMA, RAGE, F-actin, paxillin, phospho-ERK1/2, and NF-κB expression levels, *z*-stack images were acquired with a 1 μm step size. Imaging conditions were kept constant across experiments for comparative analysis. All quantitative analyses were performed using Imaris 10.2 with consistent parameters.

### Rosmarinic acid treatment

Mice were randomized into different treatment groups and sustained for 21 days after bleomycin treatment, or for 42 days after silicosis induction. Rosmarinic acid was dissolved in saline and administered by oral gavage every 24 h at low dose (LD, 15 mg/kg), medium dose (MD, 30 mg/kg), or high dose (HD, 60 mg/kg). Treatment was maintained for 21 days (days 1-21) in the bleomycin treated mice. To simulate the effects of treatment on a fully developed case of fibrosis, high dose (HD, 60 mg/kg) treatment continued for 28 days (days 14-42) in the silicosis model. Solutions were freshly prepared and sonicated to ensure complete dissolution before administration. For the bleomycin-treated mice, rosmarinic acid was first administered 1 h after bleomycin injection. Mice were euthanized 24 h after the final treatment, and lung samples were collected for subsequent analyses.

### Ventilation-based mechanotherapy

To evaluate mechanical ventilation as a therapeutic intervention, we applied mechanotherapy to 8-week-old male C57BL/6J mice with bleomycin-induced pulmonary fibrosis. Mice received a single intratracheal instillation of bleomycin (BLM, 1.5-2 U/kg body weight) following anesthesia with tribromoethanol (1.2 mL/100 g, intraperitoneal). Mechanical ventilation therapy was initiated seven days post-BLM administration using a RWD VentStar small-animal ventilator (Model R415). The protocol consisted of 30-minute ventilation sessions administered at 48-hour intervals from day 7 to day 21. During each session, mice were ventilated with tidal volumes of either 0.3, 0.6, or 0.9 mL while maintaining a positive end-expiratory pressure (PEEP) of 2 cm H₂O. The respiratory rate was maintained at 120 breaths per minute with an inspiration-to-expiration ratio of 1:2. Throughout each ventilation session, mice were kept under light anesthesia on a heated pad to maintain body temperature, with continuous monitoring of vital signs. At the experimental endpoint (day 21), mice were euthanized and lung tissues were harvested for comprehensive histopathological and biochemical analyses as described above to evaluate therapeutic outcomes.

### Bronchoalveolar lavage fluid collection

Each mouse underwent bronchoalveolar lavage twice using 0.5 mL of PBS to obtain bronchoalveolar lavage fluid (BALF). BALF samples were centrifuged at 60 *G* (*G* = 9.81 m/s^2^, Eppendorf, Centrifuge 5424R) for 5 minutes at 4°C. The resulting supernatants were then preserved at −80°C until further analysis.

### Hydroxyproline assay

The quantification of hydroxyproline (HYP) content in murine lung tissues was performed using an HYP measurement kit (NBP2-59747, Novus Biologicals, Littleton, CO, USA), according to the provided instructions by the manufacturer. Each analysis utilized approximately 30 mg of lung tissue from individual mice.

### Enzyme-linked immunosorbent assay

We utilized several enzyme-linked immunosorbent assay (ELISA) kits to assess the protein levels of IL-1β (MLB00C, RD), IL-6 (M6000B, RD), TGF-β (DB100C, RD).

### Western blot

Fresh lung tissues were lysed using RIPA lysis buffer (Beyotime, P0013b, Shanghai, China) to extract total protein. The protein concentration was determined with the BCA Protein Analysis kit (Thermo Fisher Scientific, 23225, Waltham, MA, USA). Protein samples were separated on 8%-12% SDS-polyacrylamide gels. Membranes were blocked with a 5% skimmed milk solution at room temperature for 1 hour. Following this, the membranes were incubated with primary antibodies overnight at 4℃, and then with secondary antibodies for 1 hour at room temperature. Protein signals were visualized using a Tanon automated chemiluminescence fluorescence image analysis system (5200, Tanon, Shanghai, China), and image analysis was performed using Image J software (Image J 1.53a, National Institutes of Health, USA).

### Collagen fiber network simulation

A Mikado-type collagen fibril network^57^ was modeled within a square domain with randomly oriented and distributed fibers. Each fiber, with a diameter of 100 nm and a length of 7.5 µm, was simulated as a truss element within a box of dimensions 50 × 50 µm. Each model had 600 fibers. Intersections between fibers were treated as crosslinks and modeled as moment-free nodes. Elements were pinned bottom edge and a top edge subject to pulling in the *y*-direction, while free to contract in the *x*-direction. The left edge was fixed in the *x*-direction but free to displace in the *y-*direction. Simulations were conducted using the dynamic explicit method in the Abaqus environment (Simulia, France), and a large deformation formulation was chosen. To simulate the effect of attenuating AGE crosslinking, intersections between fibers were removed at random, and the four fibers meeting at a node replaced with two overlapping, contiguous fibers. A relative crosslink density was defined as 𝜒 = 𝑁/𝑁_m_, where *N* is the number of crosslinks in a model and 𝑁_m_ is the total number of fiber crossings. This deletion process was divided into eight groups, each representing a specific fraction of crosslink density, to observe the impact of reducing crosslink density on the network’s mechanical response.

### Collagen scaffold simulation

Collagen scaffolds were modeled as porous elastic materials. 2D finite element models were created using Voronoi construction techniques. This process began by randomly placing *N* nuclei in a 2D space, ensuring that the distance between any two nuclei exceeds a minimum threshold. The space is then divided into *N* closed cells based on Delaunay triangulation^58^. To achieve curved edges, a piecewise quadratic spline function was applied, assuming a uniform cell wall thickness of h. Finite element simulations demonstrated that the pore morphology remained largely unchanged compared to the collagen fiber network results. This stability is mainly due to the significantly thicker cell walls in the collagen scaffold, which increase bending stiffness, allowing the scaffold to bear greater moments and resist collapse.

### Theoretical modeling

Cells within pores of porous scaffolds were modeled as adherent to planar substrates with Poisson ratio 𝜈 that were strained cyclically with an amplitude 𝜀*_a_*(𝑡) = (𝜀*_max_*/2) (1 + sin 𝜔𝑡) (**Fig. 4h**). For affine deformation, the axial strain on actin stress fibers oriented an angle 𝜃 from the direction of stretch was:

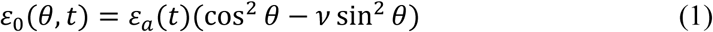

We modeled cell responses to cyclic stretch using a Kaunas-type kinetics model^59,60^. We tracked the evolution over time *t* of the areal density 𝐶*_b_* of integrin-RGD peptide (arginine-glycine-aspartic acid) bonds within each focal adhesion, and of the areal density 𝐶*_f_* of contractile actomyosin units that accumulate:

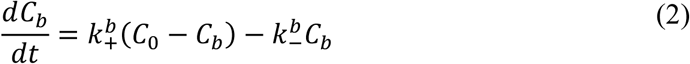

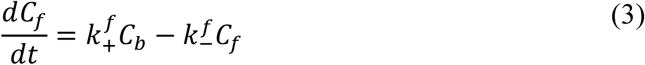

where 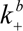 and 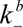 are the association and disassociation rates of integrin-RGD bonds, 𝐶_O_ is the saturation value for 𝐶*_b_*, and 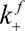 and 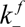 are the association and disassociation rates of contractile units. Following Qian, et al.,^61^ Eq. (3) modeled the tendency of focal adhesions to recruit contractile actomyosin machinery. Following a Bell-type model, as employed by Qian, et al.,^61^ focal adhesion mechanosensitivity arose from force-dependent integrin-RGD dissociation according to:

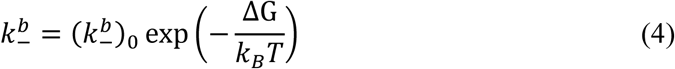

in which 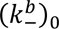 is the baseline dissociation rate, 𝑘*_B_*𝑇 is thermal energy at absolute temperature *T*, and the force-dependent energy release Δ𝐺(𝑓) associated with a single integrin-RGD bond is:

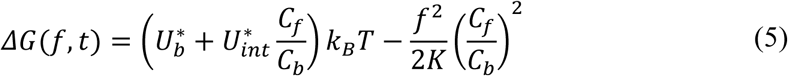

Here, 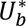 is the dimensionless energy release associated with a single integrin-RGD bond, 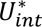 is the dimensionless additional energy associated with actomyosin recruitment to the growing focal adhesion, 𝐶*_f_*/𝐶*_b_* represents the number of actomyosin contractile units per integrin-RGD bond, and the final term is a Bell-like elastic barrier to bonding. In the last term, 𝐾(𝜀*_a_*) is the secant stiffness per bond of the ECM (assuming the ECM to be less stiff than the integrin-RGD bond), written according to a power law that accounts for the strain-stiffening associated with fibrous materials:^62,63^

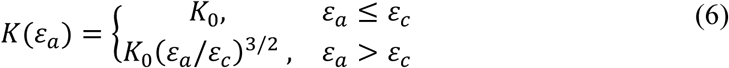

The force per actomyosin contractile unit followed a Hill-type model (**Extended Data Fig. 18**), with a nominal contractile stress 𝜎_O_ in parallel with a passive Maxwell element of modulus 𝐸*_f_* and viscosity 𝜂*_f_*, so that:

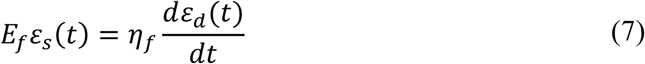

Noting that the strain 𝜀_s_ in the spring and the strain 𝜀*_d_* in the dashpot must equal that of the stress fiber (𝜀_O_(𝜃, 𝑡) = 𝜀*_s_*(𝑡) + 𝜀*_d_*(𝑡)) and enforcing equilibrium (the stress in a stress fiber, 𝜎(𝑡) = 𝜎_O_ + 𝐸*_f_*𝜀*_s_*(𝑡)) enables solution of Eqs. (1) - (7). In dimensionless form, with 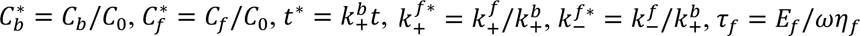 and 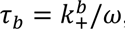, the governing equations can be rewritten:

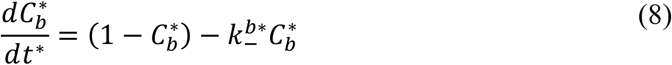

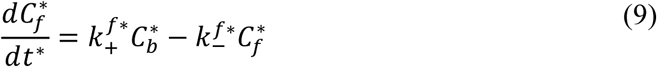

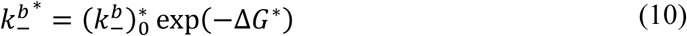

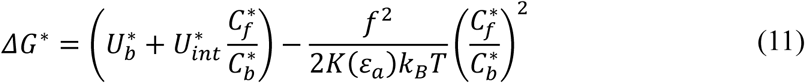

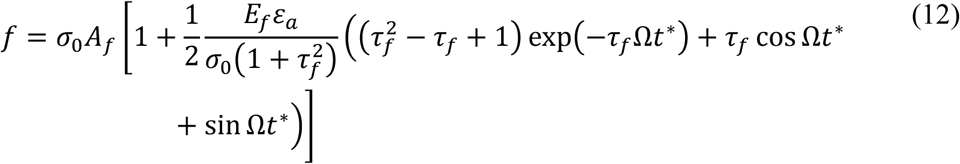

Here, 𝐴*_f_* is the nominal cross-sectional area of an actomyosin stress fiber in its reference configuration. Simulation parameters ans initial conditions are listed in Supplementary Table 4.

### Statistics

Statistical analyses were performed using GraphPad Prism version 9.0.0. Prior to analysis, potential outliers were identified and removed using the ROUT method with Q = 1%. Data normality was evaluated using two approaches: the D’Agostino-Pearson omnibus test for larger datasets and the Shapiro-Wilk test when sample sizes were small. For comparisons between two independent groups, we employed either unpaired two-tailed t-tests (for normally distributed data) or Mann-Whitney tests (for non-normally distributed data). When analyzing three or more groups, we used one-way ANOVA followed by Tukey’s multiple comparisons test if the data were normally distributed. For non-normally distributed data involving multiple groups, we utilized the Kruskal-Wallis test followed by Dunn’s multiple comparisons test. Throughout the manuscript, all data are presented as mean ± standard deviation (S.D.). Exact *p* values are provided in each figure to facilitate interpretation of statistical significance. Detailed information regarding sample sizes, specific statistical tests used, and experimental reproducibility is available in the corresponding figure legends.

### Data availability

Data supporting the findings of this paper are available within the article and its Supplementary Information files. The data can also be obtained from the corresponding authors upon request.

## Acknowledgments

We acknowledge Weihua Wang, Fang Wei at Center of Pharmaceutical Technology, Tsinghua University for assistance of HPLC/MS. We thank Bingyu Liu at Imaging Core Facility, Technology Center for Protein Sciences Tsinghua University for assistance of using Imaris 10.2. and confocal microscopy. We thank Dr. Canhong Xiang and Dr. Wei Yang of the Department of Hepatobiliary Surgery, Beijing Tsinghua Changgung Hospital for their important help in the clinical liver sample study. We thank Dr. Yan Zhang of Tsinghua University for guidance on tissue decellularization. We thank Dr. Hanxun Jin of Washington University in St. Louis for guidance on simulations. This work was funded by the National Natural Science Foundation of China (82125018, 32430058, 12402364), the Human Frontier Science Program (HFSP - RGP016/2024) and the US NSF through the NSF Science and Technology Center for Engineering Mechano-Biology (grant CMMI 1548571).

## Author contributions

W.K. and Y.D. conceived of and designed the research. W.K. characterized tissue samples and performed the in vitro work. M.S., J.W., L.B., X.W., and W.K. performed the animal experiments with guidance from C.W., J.W., and Y.D. X.P., and Y.H. performed computational simulations with guidance from G.M.G. and X.F. J.Z., Y.J. performed AFM characterization. C.W. and J.W. provided clinical tissue samples and clinical consultation. K.L. prepared the illustrations. Y.N. performed RNA-seq data analysis. L.Z. helped establish the in vitro stretching system. W.K., G.M.G., and Y.D. wrote the manuscript. Y.D., J.W., and G.M.G. are the principal investigators of the supporting grants.

## Competing interests

Authors declare that they have no competing interests.

### Additional information

**Extended Data Figures 1-18.**

**Supplementary information** includes supplementary note, supplementary figures 1-2, supplementary tables 1-4, supplementary videos 1-3 legends.

**Supplementary videos 1-3.**

### Extended Data Figures

**Extended Data Fig. 1.**
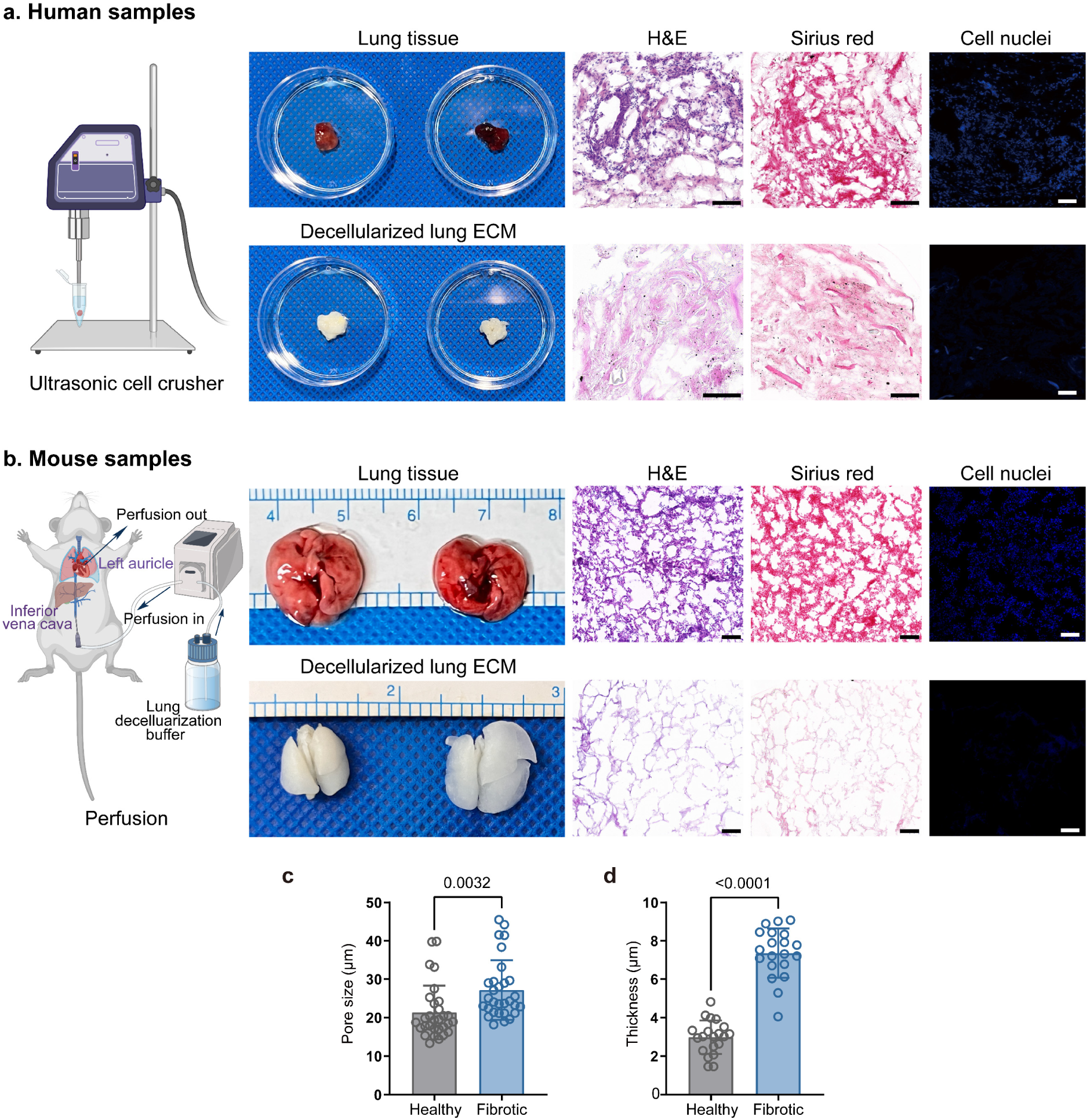
Species-specific decellularization protocols preserve ECM architecture critical for evaluating crosslinking mechanisms in pulmonary fibrosis. (**a**) Human lung decellularization via ultrasonic disruption, validated through multiple staining methods: H&E for tissue architecture, Sirius red for collagen distribution, and DAPI confirming removal of cell nuclei. This technique enables isolation of native ECM while preserving architectural features essential for accurate analysis of crosslinking patterns. Scale bars: 100 μm. (**b**) Mouse lung decellularization via whole-organ perfusion, with corresponding validation of ECM preservation and cell removal. This approach maintains the 3D porous architecture unique to lung tissue, allowing direct assessment of how crosslinking affects mechanical properties. Scale bars: 100 μm. (**c**, **d**) Quantitative comparison of ECM architecture in healthy versus fibrotic tissue: (**c**) pore size and (**d**) wall thickness measured from SEM images (*n* = 30 measurements from ≥5 independent fields per condition). These architectural alterations represent structural changes driven by AGE-mediated crosslinking that contribute to the restricted mechanical movement and altered cell-ECM interactions associated with disease progression. Statistical analysis performed using two-tailed unpaired t-test, exact *p* values labeled. Results presented as mean ± S.D.

**Extended Data Fig. 2.**
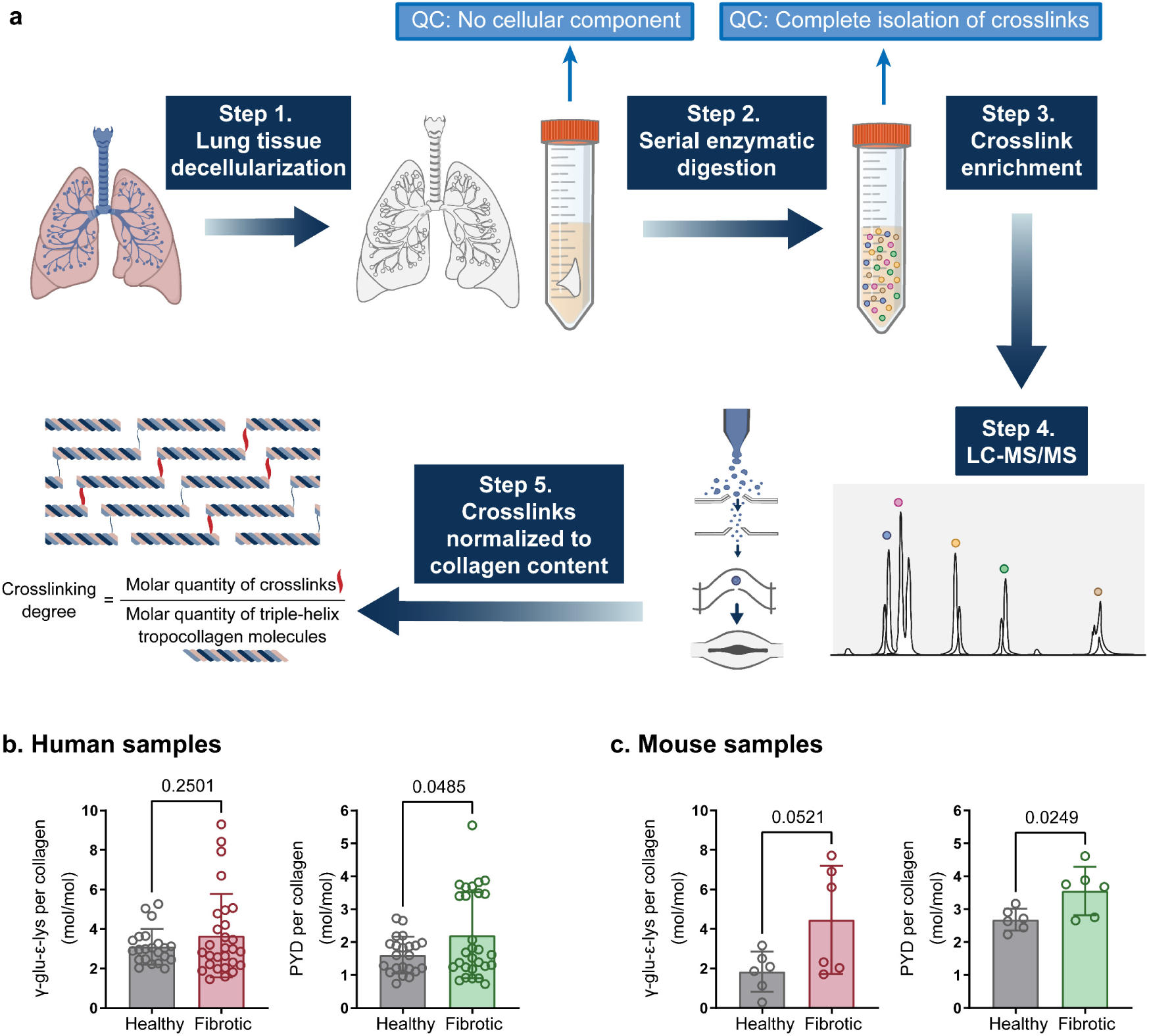
Quantification of matrix crosslinking reveals AGE-mediated crosslinking as a dominant mechanism in pulmonary fibrosis. (**a**) Absolute Quantification of Matrix-specific Crosslinking (AQMC) method. Steps: 1) Harvest lung tissue and verify complete decellularization; 2) Enzymatically digest ECM to dipeptides and amino acids; 3) Enrich crosslinked products; 4) Quantify crosslinks and hydroxyproline using LC-MS/MS; 5) Calculate crosslinking density as molar ratio of crosslinks per tropocollagen molecule. This analytical approach enables precise quantification of the molecular mechanisms driving ECM remodeling in fibrotic disease. (**b**, **c**) Enzymatic crosslinking levels in (**b**) human and (**c**) mouse lung ECM (*n* ≥ 6 ECM samples from 5 independent mice per condition), showing both transglutaminase-mediated crosslinking (TGM, measured via γ-glutamyl-ε-lysine) and lysyl oxidase-mediated crosslinking (LOX, measured via pyridinoline). The significantly higher levels of AGE crosslinking compared to enzymatic crosslinking provide mechanistic insight into why previous therapeutic approaches targeting LOX have shown limited efficacy in pulmonary fibrosis, supporting our focus on AGE-mediated mechanisms. Statistical analysis performed using two-tailed unpaired t-test, exact *p* values labeled. Results presented as mean ± S.D.

**Extended Data Fig. 3.**
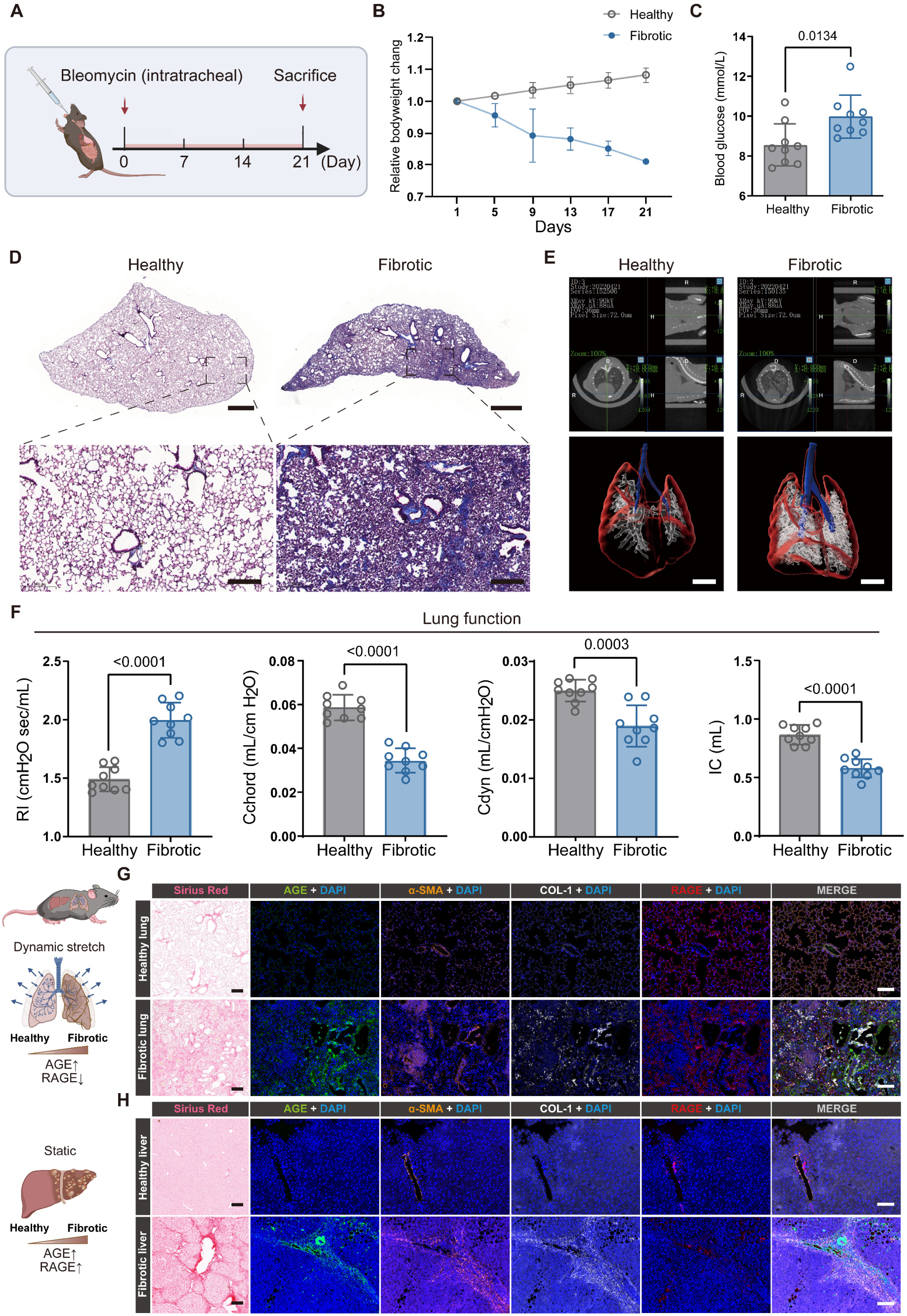
Multi-scale characterization of bleomycin-induced pulmonary fibrosis model demonstrates altered mechanics, matrix remodeling, and paradoxical RAGE expression. (**a**) Experimental timeline for bleomycin (BLM) administration and analysis at doses sufficient to establish a model of persistent fibrosis without self-recovery. (**b**) Progressive weight loss following BLM treatment (*n* = 9 mice per group), confirming systemic impact of disease. (**c**) Elevated blood glucose levels in fibrotic versus healthy mice (*n* = 9 mice per group), highlighting metabolic dysregulation that may contribute to AGE formation and crosslinking. (**d**) Masson’s trichrome staining revealing increased collagen deposition (blue) in BLM-treated lungs, a hallmark of fibrotic remodeling. Scale bars: 1000 μm overview, 200 μm detail. (**e**) Micro-CT reconstruction showing increased ECM density (white) in fibrotic lungs, providing 3D confirmation of tissue remodeling. Scale bars: 4000 μm. (**f**) Comprehensive lung function analysis showing BLM-induced changes in: resistance index (RI), chord compliance (Cchord), dynamic compliance (Cdyn), and inspiratory capacity (IC) (*n* = 9 mice per group), establishing functional consequences of the observed structural remodeling. (**g**, **h**) Comparison of fibrotic remodeling in human (**g**) lung and (**h**) liver tissue, visualized through Sirius red staining and immunofluorescence for AGE crosslinking, collagen I (COL-1), α-smooth muscle actin (α-SMA), and receptor for advanced glycation end products (RAGE). This organ-specific comparison reveals a paradoxical decrease in RAGE expression in fibrotic lungs compared to the increase in fibrotic liver, suggesting that unique mechanobiological feedback mechanisms driving pulmonary fibrosis. Scale bars: 200 μm. Statistical analysis performed using two-tailed unpaired t-test, exact *p* values labeled. Results presented as mean ± S.D.

**Extended Data Fig. 4.**
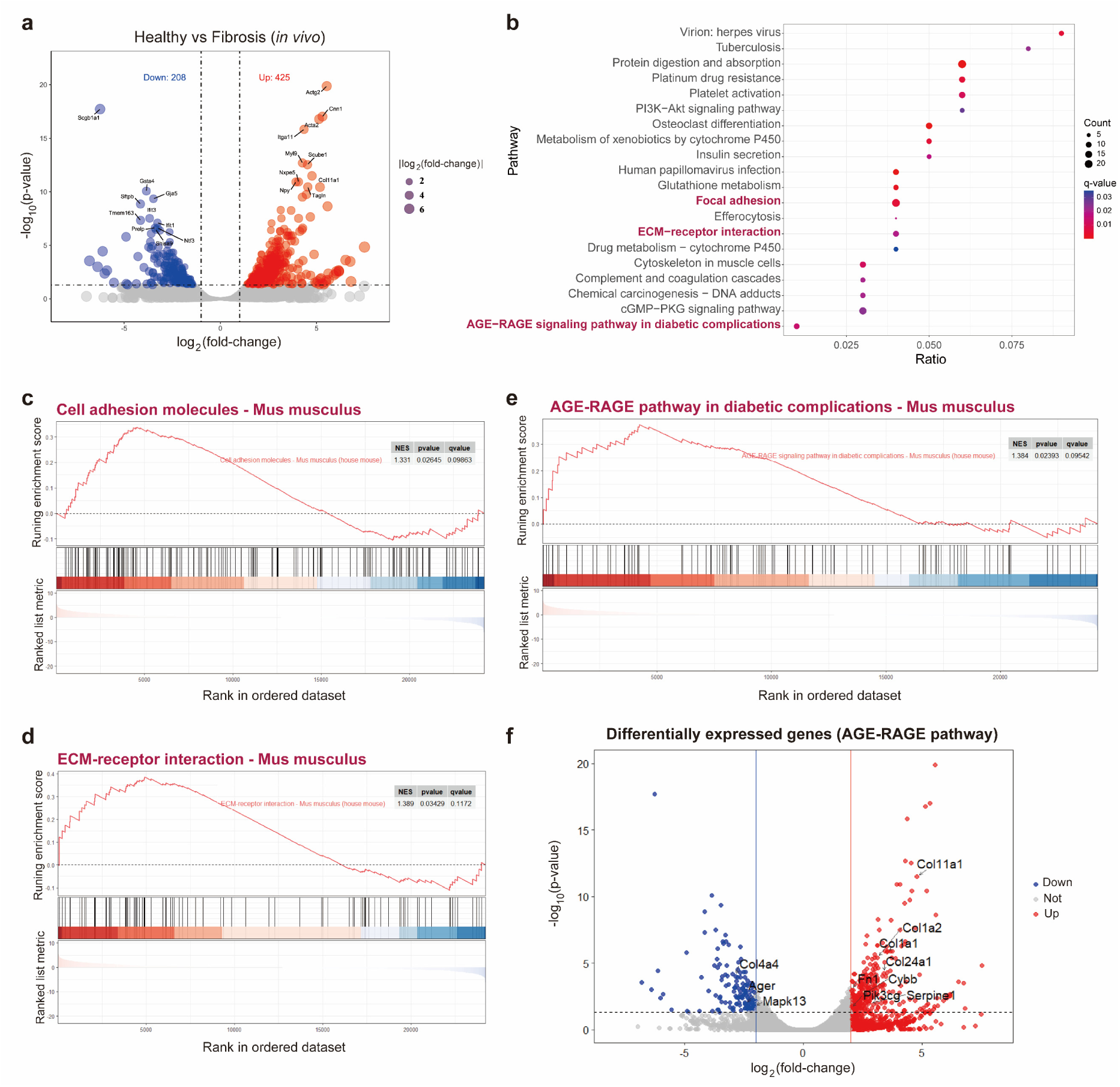
RNA-seq transcriptional analysis reveals activation of AGE-RAGE and adhesion pathways in fibrotic lung fibroblasts, establishing a molecular basis for mechanobiological feedback. (**a**) Volcano plot comparing gene expression in primary pulmonary fibroblasts (pPFBs) isolated from healthy versus fibrotic mice. Significant differential expression defined as adjusted *p*-value < 0.05 and |log_2_FC| > 1, calculated using the Wald test with Benjamini-Hochberg correction. This transcriptional profiling identified molecular signatures of activated fibroblasts in the disease state. (**b**) Pathway enrichment analysis identifying upregulation of AGE-RAGE signaling, focal adhesion, ECM-receptor interaction, and PI3K-Akt signaling pathways in fibrotic pPFB (adjusted *p*-value < 0.05). Enrichment of these mechanosensitive and AGE-related pathways provides molecular support for our hypothesized mechanobiological feedback in fibrosis progression. (**c-e**) Gene Set Enrichment Analysis (GSEA) showing enrichment of (**c**) cell adhesion molecules, (**d**) ECM-receptor interaction, and (**e**) AGE-RAGE signaling pathways in fibrotic versus healthy pPFB, with normalized enrichment scores (NES) calculated by permutation test. These pathway changes demonstrate that fibroblasts couple increased mechanical sensing with AGE-RAGE signaling to perpetuate the fibrotic state. (**f**) Detailed expression changes of individual genes within the AGE-RAGE signaling pathway showing paradoxical downregulation of RAGE (*Ager*) despite pathway activation, highlighting unique regulation of this pathway in pulmonary fibrosis. NES, normalized enrichment score; *p*-values calculated with permutation tests.

**Extended Data Fig. 5.**
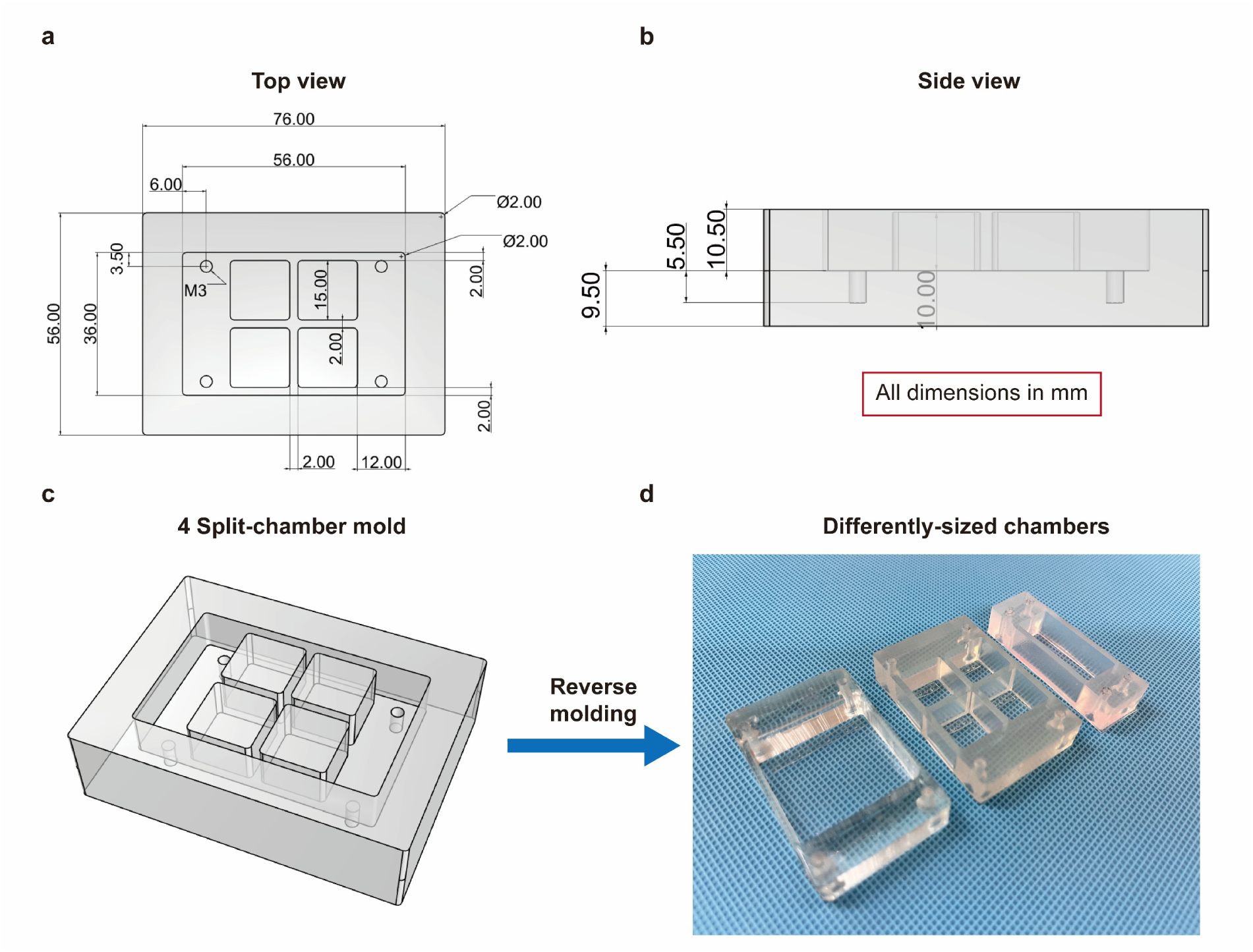
Design and fabrication of multi-part stretching chambers. (**a**) Engineering drawing: top view of four-part stretching chamber mold showing critical dimensions and assembly details. (**b**) Side-view engineering drawing detailing vertical dimensions and mold interfaces. (**c**) Three-dimensional assembly diagram illustrating component relationships and mold separation planes. (**d**) Photographs of final molded chambers at different scales demonstrating scalability of the design. All dimensions in mm.

**Extended Data Fig. 6.**
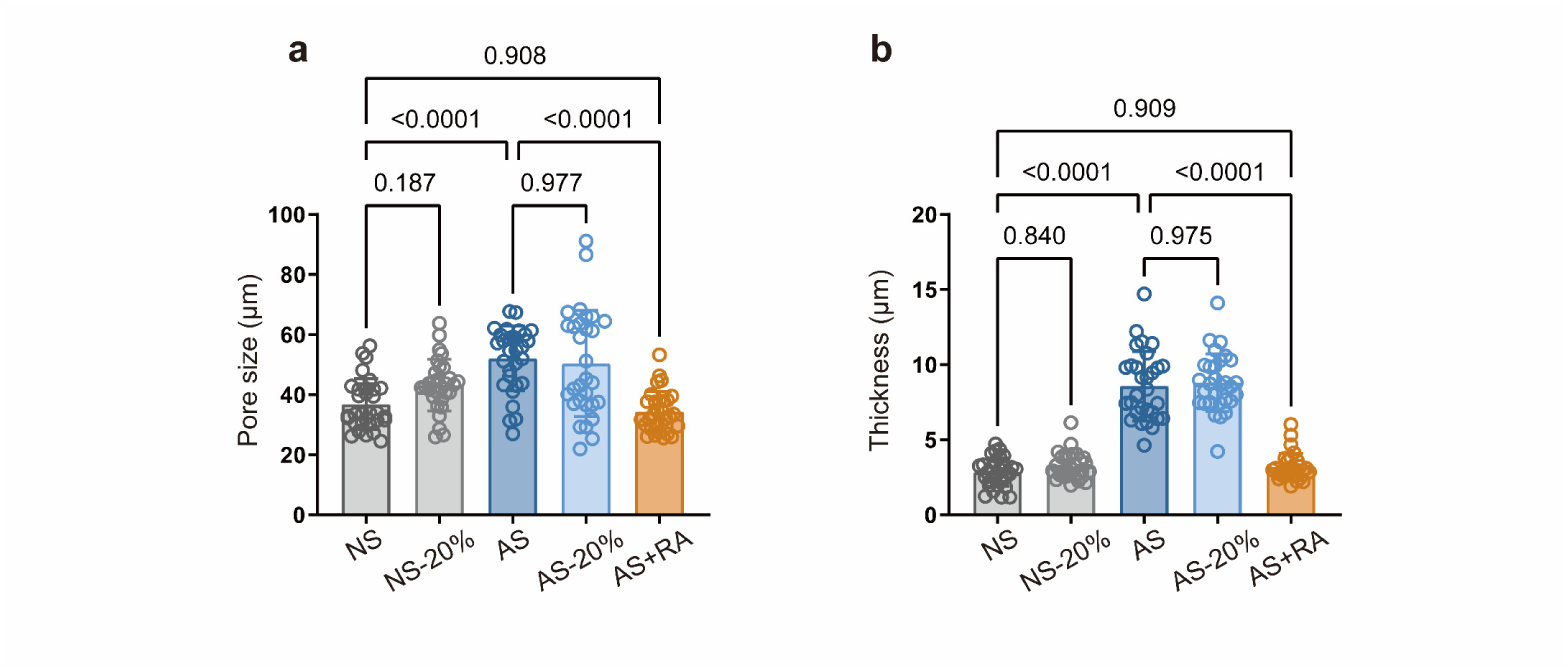
Quantitative analysis of scaffold architecture under different treatments. (**a**) Pore size and (**b**) wall thickness measurements from SEM images of non-crosslinked (NC), AGE-crosslinked (AGE), rosmarinic acid-treated (AGE+RA), and dynamically stretched (20% strain) scaffolds (*n* = 30 measurements from ≥5 independent SEM fields per condition, corresponding to images in Fig. 3c-d). Statistical analysis performed using one-way ANOVA with Tukey’s post-hoc test, exact *p* values labeled. Results presented as mean ± S.D.

**Extended Data Fig. 7.**
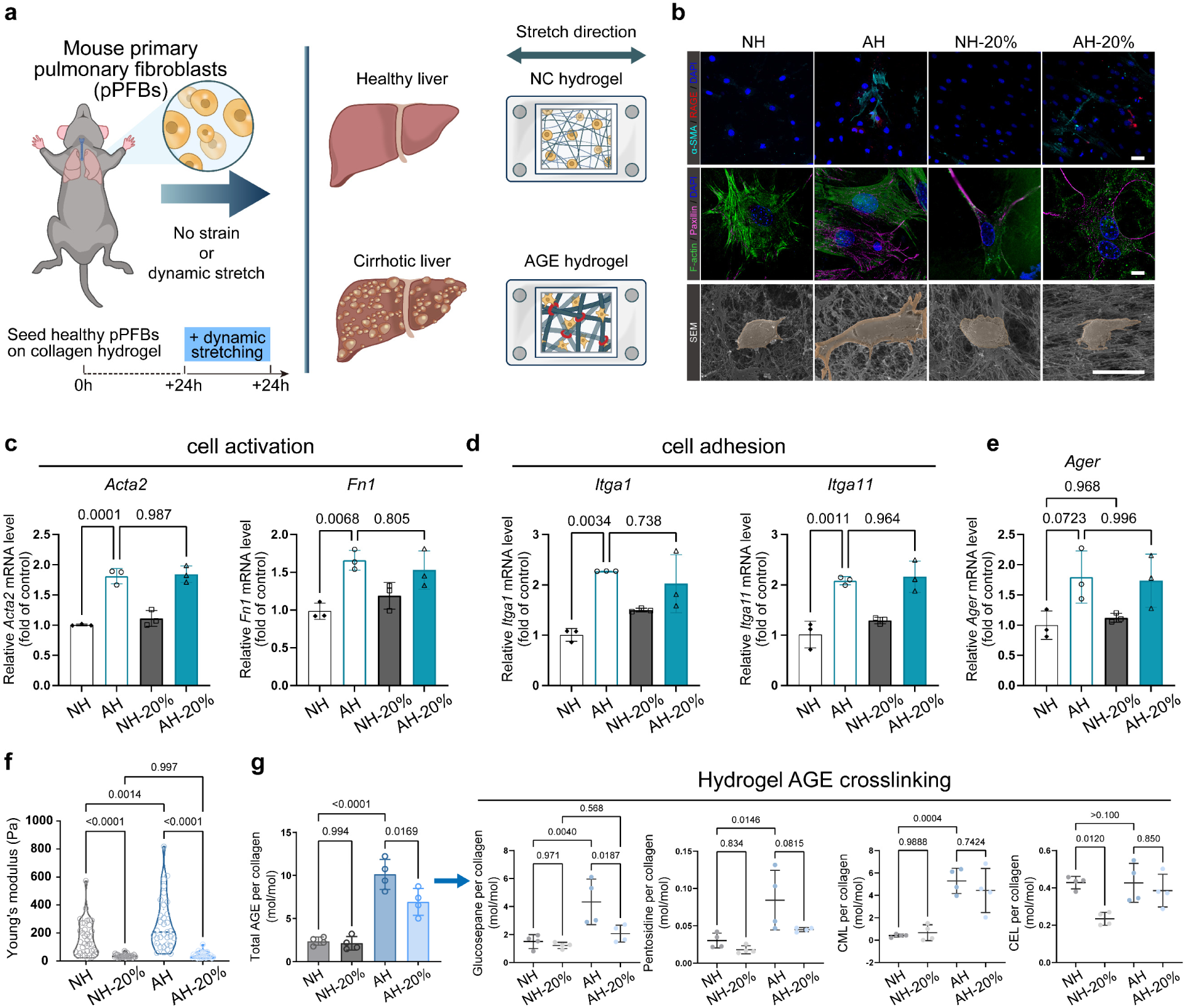
Physiological dynamic stretching cannot resist AGE-crosslinked collagen matrix induced pPFB activation on fibrous hydrogel matrix, demonstrating matrix architecture-specific mechanosensitivity. (**a**) Schematic of isolating primary pulmonary fibroblasts from lung tissues for seeding on collagen hydrogel matrix, enabling direct comparison with the porous scaffold system. (**b**) Representative images of α-SMA, RAGE, paxillin, F-actin staining and SEM of pPFBs grown on non-crosslinked hydrogel (NH), AGE-crosslinked hydrogel (AH), and hydrogel with 20% dynamic stretching strain (NH-20%, AH-20%). Top panels, detailed view showing the expression of α-SMA (cyan), RAGE (red) and cell nuclear (blue), Scale bars, 50 μm. Middle panels, detailed view showing the expression of F-actin (green) and paxillin (amaranth), Scale bars, 10 μm. Bottom panels, SEM images of pPFBs morphology. Scale bars, 20 μm. This analysis reveals a critical distinction between organ-specific ECM architectures: unlike in porous scaffolds, dynamic stretching fails to reverse fibroblast activation in fibrous hydrogels. (**c-e**) The relative mRNA expression of *Acta2*, *Fn1*, *Ager*, *Itga1*, I*tga11* in non-crosslinked and AGE-crosslinked collagen hydrogel, confirming persistent activation despite mechanical stretching. (**f**) Young’s modulus of NC hydrogel, AGE hydrogel, and hydrogel with 5%, 20% dynamic stretching strain measured by AFM (n ≥ 36, points of measurement randomly selected from at least 10 fields per sample), showing that dynamic stretching reduces stiffness regardless of crosslinking state, unlike the differential effects seen in porous scaffolds. (**g**) Quantification of AGE crosslinking degree in reconstructed collagen hydrogel with dynamic stretching (*n* = 4, independent pieces of collagen matrix), demonstrating that mechanical reduction of crosslinking alone is insufficient to reverse fibroblast activation without the porous architecture characteristic of lung ECM. Statistical analysis performed using one-way ANOVA with Tukey’s post-hoc test, exact *p* values labeled. Results are presented as mean ± S.D.

**Extended Data Fig. 8.**
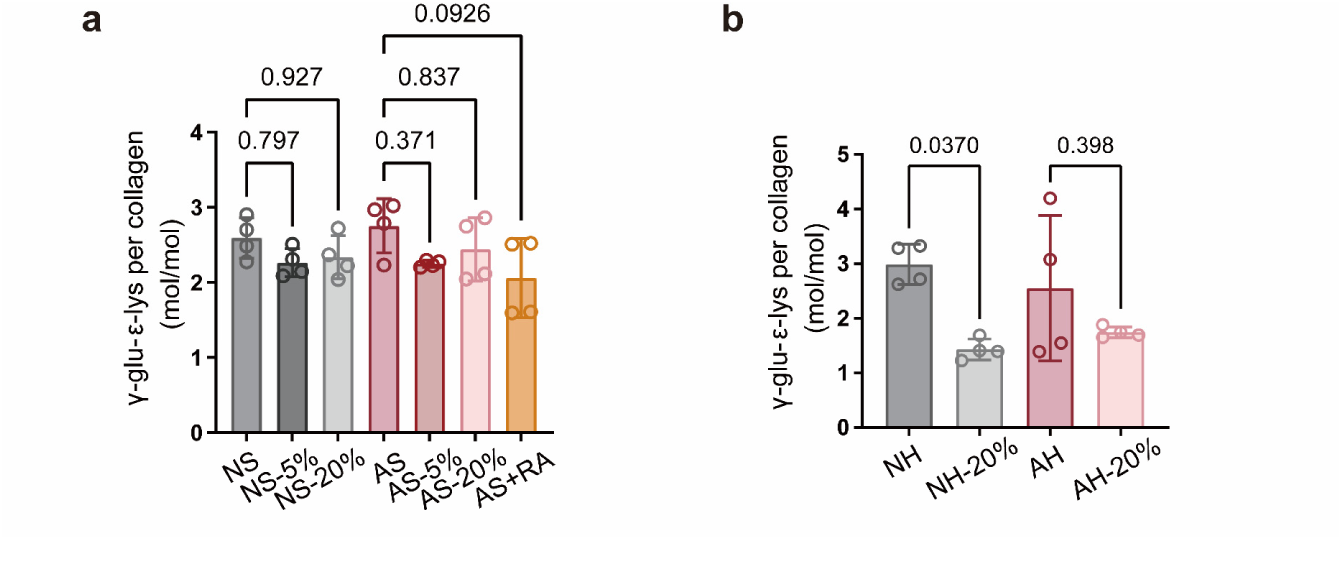
Dynamic stretch had no significant effect on TGM-crosslinking degree, demonstrating specificity of mechanical disruption for AGE crosslinks. (**a**) Quantification of TGM crosslinking degree in reconstructed collagen scaffold with dynamic stretching or RA treatment (n = 4, independent pieces of collagen matrix). This selective resistance of TGM crosslinks to mechanical disruption confirms that the therapeutic effects of dynamic stretching are specific to AGE-mediated crosslinking, suggesting mechanisms underlying why physiological breathing can selectively target pathological matrix modifications while preserving normal ECM structure. (**b**) Quantification of TGM crosslinking degree in reconstructed collagen hydrogel with dynamic stretching (n = 4, independent pieces of collagen matrix), further demonstrating that crosslink specificity is maintained across different ECM architectures. The stability of TGM crosslinks under the dame stretching regimes that disrupt AGE crosslinks highlights fundamental susceptibility of AGE’s the stretch and suggests mechanisms for how the lung’s mechanical microenvironment selectively modulates pathological but not physiological crosslinking. Statistical analysis performed using one-way ANOVA with Tukey’s post-hoc test, exact *p* values labeled. Results are presented as mean ± S.D.

**Extended Data Fig. 9.**
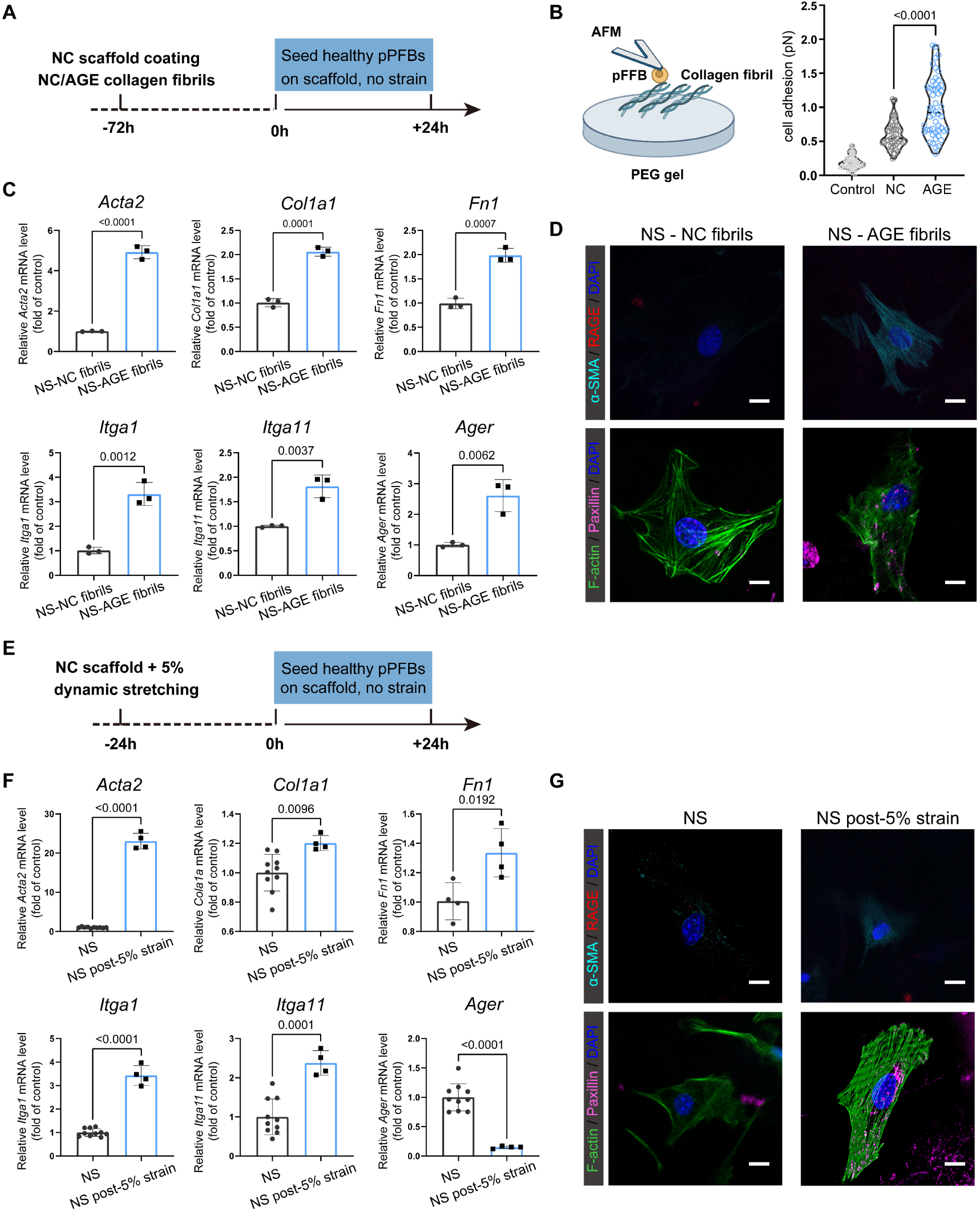
AGE crosslinking and matrix elasticity regulate RAGE expression and cell adhesion via distinct mechanisms, revealing separate components of the mechanobiological feedback loop in pulmonary fibrosis. (**a**) Schematic of pPFBs characterization on non-crosslinked scaffolds with NC collagen fibrils (NS-NC fibrils) or AGE-crosslinked collagen fibrils (NS-AGE fibrils) to isolate the biochemical effects of AGE crosslinking from those of mechanical stiffness changes. (**b**) Characterization and statistical analysis of the cell adhesion between pPFBs and collagen fibrils using AFM (n ≥ 45, points of measurement randomly selected from at least 10 fields per sample), revealing significantly enhanced fibroblast adhesion to AGE-crosslinked fibrils independent of substrate stiffness. (**c**) The relative mRNA expression of *Acta2*, *Col1a1*, *Fn1*, *Itga1*, *Itga11* and *Ager* in pPFBs grown on NS-NC fibrils and NS-AGE fibrils collagen scaffolds, demonstrating that AGE crosslinking alone can drive fibroblast activation and modulate RAGE expression. (**d**) Representative images of α-SMA, RAGE, paxillin and F-actin staining of pPFBs grown on NS-NC fibrils and NS-AGE fibrils collagen scaffold. Top panels show detailed views of α-SMA (cyan), RAGE (red) and cell nuclei (blue), Scale bars, 10 μm. Bottom panels show detailed views of F-actin (green), paxillin (amaranth) and cell nuclei (blue), Scale bars, 10 μm. (**e**) Schematic of pPFBs characterization on NC scaffolds to isolate mechanical effects from biochemical AGE signaling. (**f**) Relative mRNA expression of *Acta2*, *Col1a1*, *Fn1*, *Itga1*, *Itga11* and *Ager* in pPFBs grown on NC scaffold and NC scaffold post-5% dynamic stretch strain with a higher elasticity, demonstrating that matrix stiffening in the absence of AGE crosslinking drives opposing effects on RAGE expression compared to AGE- mediated effects. (**g**) Representative images of α-SMA, RAGE, paxillin and F-actin staining of pPFBs grown on NC scaffold and NC scaffold post-5% dynamic stretch strain. Top panels show detailed views of α-SMA (cyan), RAGE (red) and cell nuclei (blue), Scale bars, 10 μm. Bottom panels show detailed views of F-actin (green), paxillin (amaranth) and cell nuclei (blue), Scale bars, 10 μm. This decoupling of mechanical and biochemical effects demonstrates bidirectional regulation of fibroblast behavior underlying the self-reinforcing cycle of pulmonary fibrosis. Statistical analysis performed using two-tailed unpaired t-test, exact *p* values labeled. Results presented as mean ± S.D.

**Extended Data Fig. 10.**
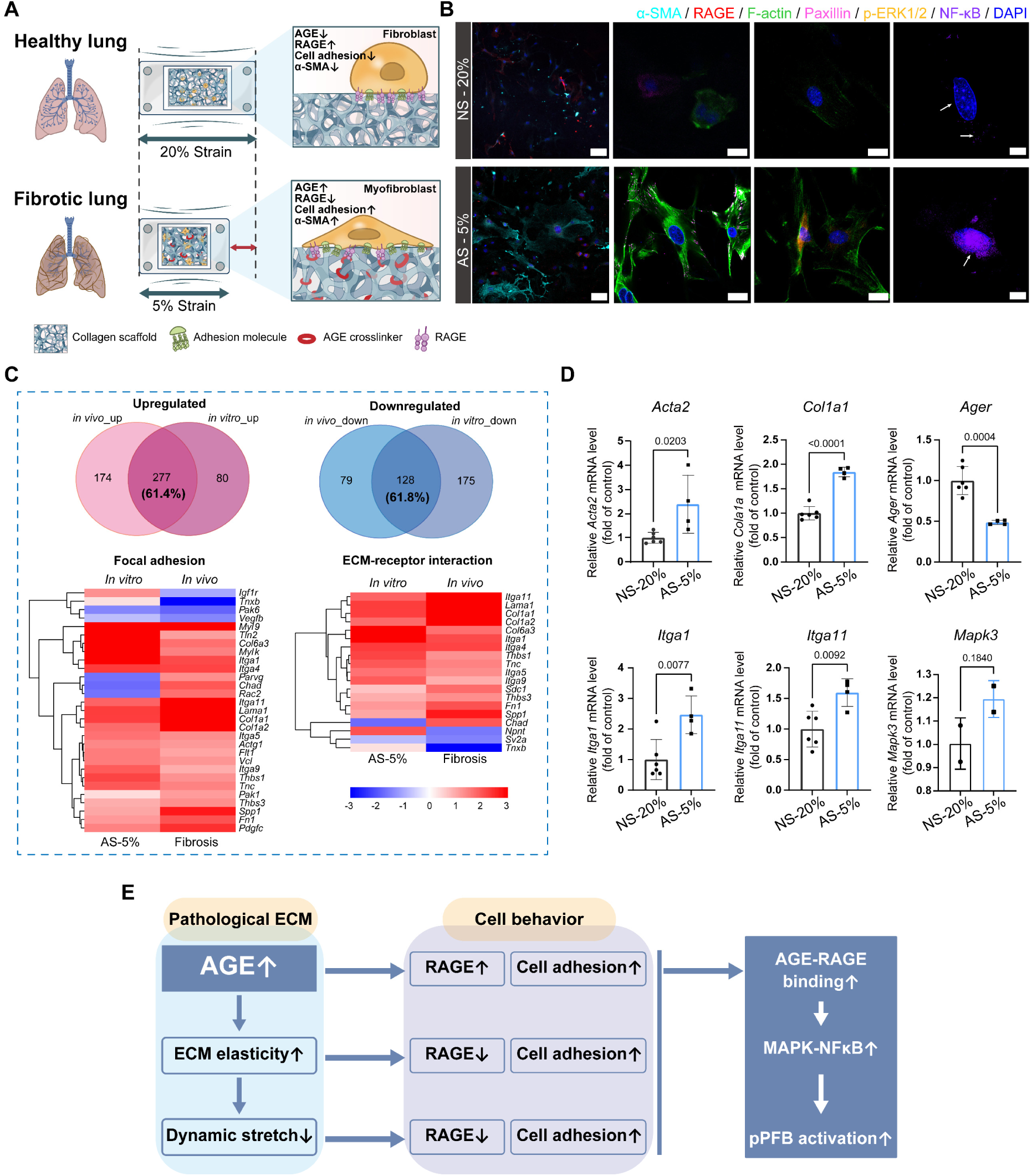
AGE crosslinking and restricted mechanical stretching cooperatively activate the AGE-RAGE-MAPK pathway in pulmonary fibrosis, establishing the molecular mechanisms underlying the mechanobiological feedback loop. (**a**) Schematic comparing pPFB responses on non-crosslinked scaffolds under 20% strain (NS-20%, mimicking healthy conditions) versus AGE-crosslinked scaffolds under 5% strain (AS-5%, mimicking fibrotic conditions). This experimental design recapitulates the dual pathological changes in fibrotic lungs: increased matrix crosslinking and restricted mechanical stretch. (**b**) Immunofluorescence showing protein expression and localization in pPFBs under healthy versus fibrotic conditions: α-SMA (cyan), RAGE (red), nuclei (blue) (left panels, scale bars 50 μm); F-actin (green), paxillin (amaranth), nuclei (blue) (second panels, scale bars 20 μm); p-ERK1/2 (orange), nuclei (blue) (third panels, scale bars 20 μm); and NF-κB (purple), nuclei (blue) (right panels, scale bars 10 μm). These images reveal that pathological conditions induce not only fibroblast activation and altered adhesion but also increased nuclear translocation of key signaling molecules in the MAPK pathway. (**c**) Heat map comparing differentially expressed genes between *in vitro* conditions (NS-20% vs AS-5%) and *in vivo* conditions (healthy vs. fibrotic) for focal adhesion, ECM-receptor interaction, and AGE-RAGE signaling pathways (*q*-value < 0.05, log₂FC > 1 or < −1). The concordance between *in vitro* and *in vivo* expression patterns validates the biomimetic model and confirms the central role of these pathways in disease progression. (**d**) Relative mRNA expression of activation markers (*Acta2*, *Col1a1*), adhesion molecules (*Itga1*, *Itga11*), MAPK pathway (*Mapk3*), and *Ager* in pPFBs under healthy vs. fibrotic conditions. (**e**) Schematic of how pathological ECM remodeling drives aberrant cellular behaviors and activates downstream signaling pathways, collectively perpetuating fibrotic progression. This molecular characterization establishes how increased AGE crosslinking and decreased mechanical stretching synergistically activate the MAPK-NF-κB pathway through altered AGE-RAGE interactions, providing a molecular basis for understanding the mechanobiological feedback loop driving pulmonary fibrosis.

**Extended Data Fig. 11.**
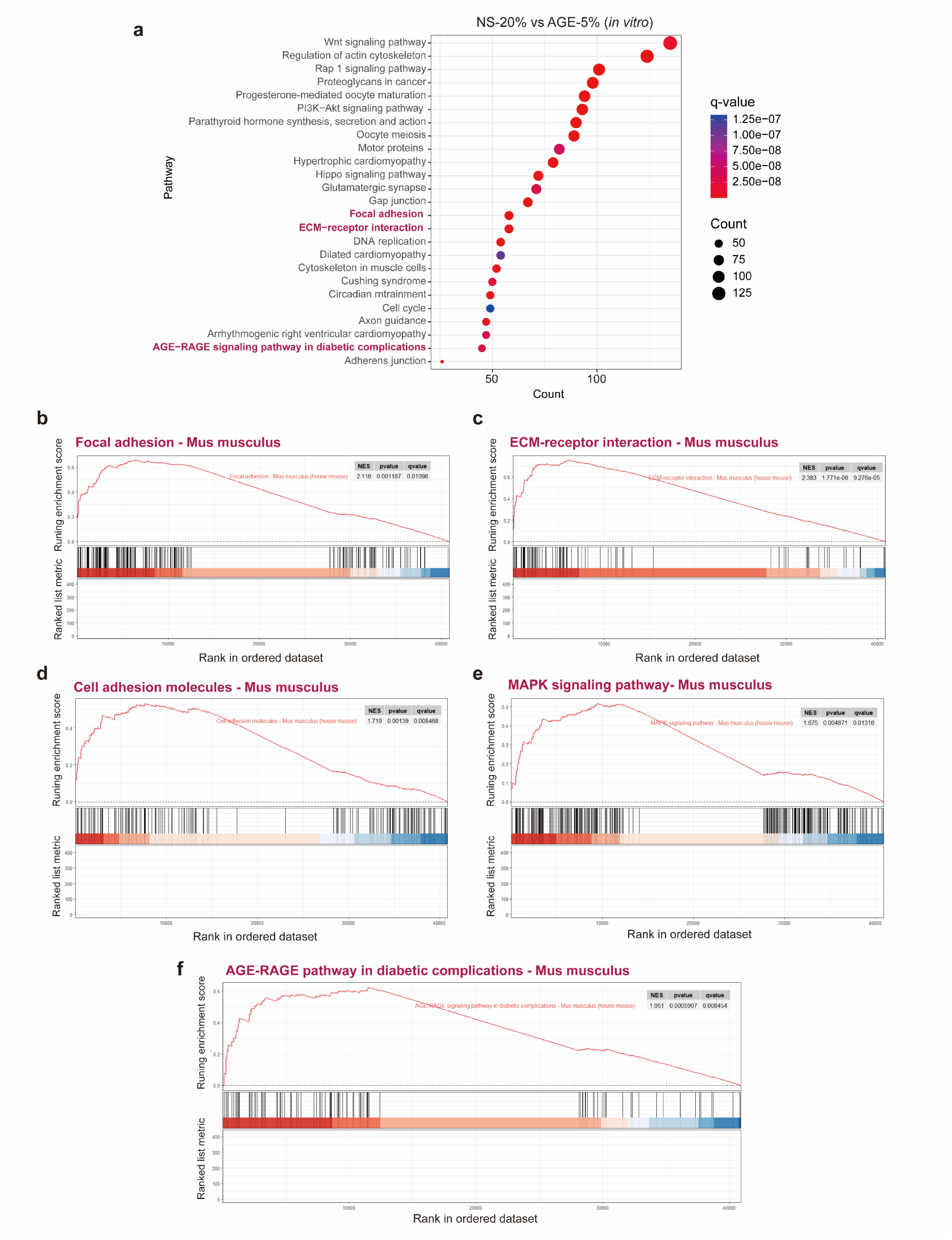
RNA-seq analysis of pPFBs cultured on scaffold matrix *in vitro* validates the mechanobiological model and reveals comprehensive pathway dysregulation mirroring pulmonary fibrosis *in vivo*. (**a**) Pathway enrichment analysis of pPFBs cultured on NS-20% and AS-5% matrix genes reveals an enrichment of genes associated with AGE-RAGE signaling pathway in diabetes complication, focal adhesion, ECM-receptor interaction (adjusted *p*-value < 0.05 and log_2_FC>1 or <-1, NS-20% versus AS-5%). This global pathway analysis confirms that the combined effects of AGE crosslinking and restricted mechanical stretching recapitulate the *in vivo* molecular signature of pulmonary fibrosis. (**b-f**) GSEA to assess focal adhesion (**b**), ECM-receptor interaction (**c**), cell adhesion molecules (**d**), MAPK signaling pathway (**e**) and AGE-RAGE signaling pathway (**f**) in pPFBs cultured on NS-20% and AS-5% matrix. These pathway analyses reveal coordinated activation of mechanosensing, adhesion, and signaling networks that link ECM properties to cellular behavior, providing molecular support for the mechanobiological feedback mechanism. Enrichment of the MAPK and AGE-RAGE signaling pathways under fibrotic conditions confirms these as key downstream effectors that translate mechanical and biochemical cues into altered fibroblast phenotypes, establishing potential therapeutic targets for breaking the self-reinforcing cycle of disease progression. NES: normalized enrichment score; *p*-values calculated with permutation tests.

**Extended Data Fig. 12.**
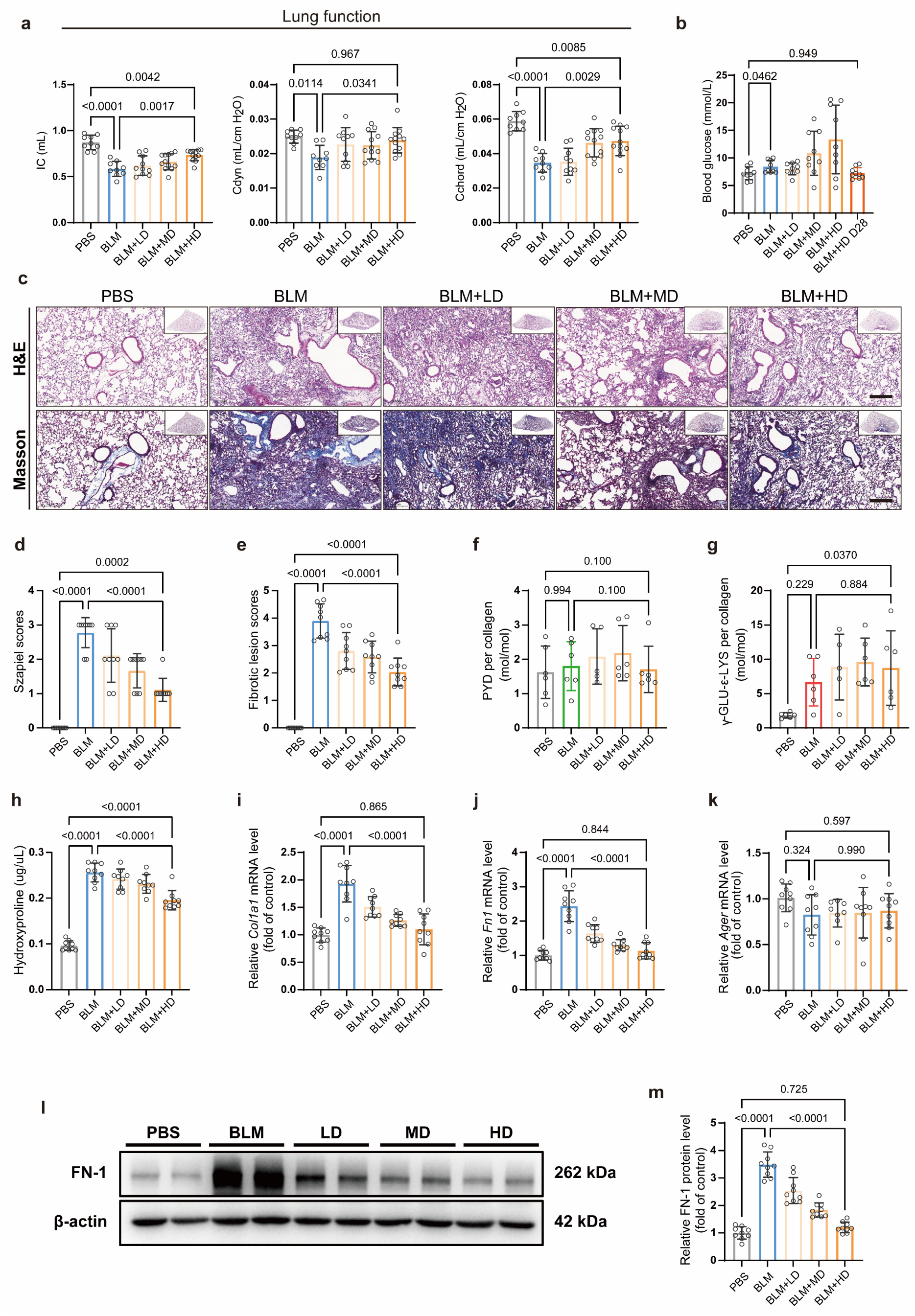
Molecular analysis confirms broad anti-fibrotic effects of rosmarinic acid treatment, establishing AGE crosslinking inhibition as a therapeutic strategy for pulmonary fibrosis. (**a**) Lung function was adversely affected by BLM treatment, but effects reduced to levels that were not significant statistically with high doses of RA. Factors studied were inspiratory capacity (IC), dynamic compliance (Cdyn), and chord compliance (Cchord). These functional improvements demonstrate that targeting AGE crosslinking can restore not only tissue structure but also physiological respiratory mechanics. (**b**) Scatter for blood glucose was large at intermediate timepoints, but resolved by day 28 to the level of control experiments, suggesting that RA treatment may also address the metabolic dysregulation associated with fibrosis and AGE formation. (**c**) Representative images of collagen deposition stained by H&E and Masson staining from lungs treated with vehicle and different doses of RA (*n* = 9 biologically independent mice per group). Scale bars, 200 μm. (**d**, **e**) Histological quantification of fibrosis severity based up (**d**) H&E and (**e**) Masson’s trichrome staining of lungs treated with vehicle or varying RA doses (*n* = 9 mice per group), providing quantitative confirmation of dose-dependent therapeutic effects. (**f**, **g**) Quantification of (**f**) LOX and (**g**) TGM crosslinking in decellularized lung ECM following RA treatment of BLM-induced fibrosis (*n* ≥ 5 ECM pieces per group), demonstrating that RA selectively targets AGE crosslinking without affecting enzymatic crosslinking mechanisms and confirming the specificity of this therapeutic approach. (**h**) Tissue hydroxyproline content as a measure of total collagen (*n* = 9 mice per group), showing significant reduction in collagen accumulation with RA treatment. (**i**-**k**) Relative mRNA expression in lung tissue of fibrosis-associated genes *Acta2*, *Col1a1*, and *Ager*, confirming molecular reversal of the fibrotic phenotype and restoration of normal RAGE expression. (**l**, **m**) Western blot analysis of fibronectin (FN-1) expression, with (**l**) densitometric quantification normalized to β-actin and (**m**) representative blots. This molecular analysis demonstrates that inhibiting AGE crosslinking with RA effectively breaks the mechanobiological feedback loop driving pulmonary fibrosis. Statistical analysis performed using one-way ANOVA with Tukey’s post-hoc test, exact *p* values labeled. Results presented as mean ± S.D.

**Extended Data Fig. 13.**
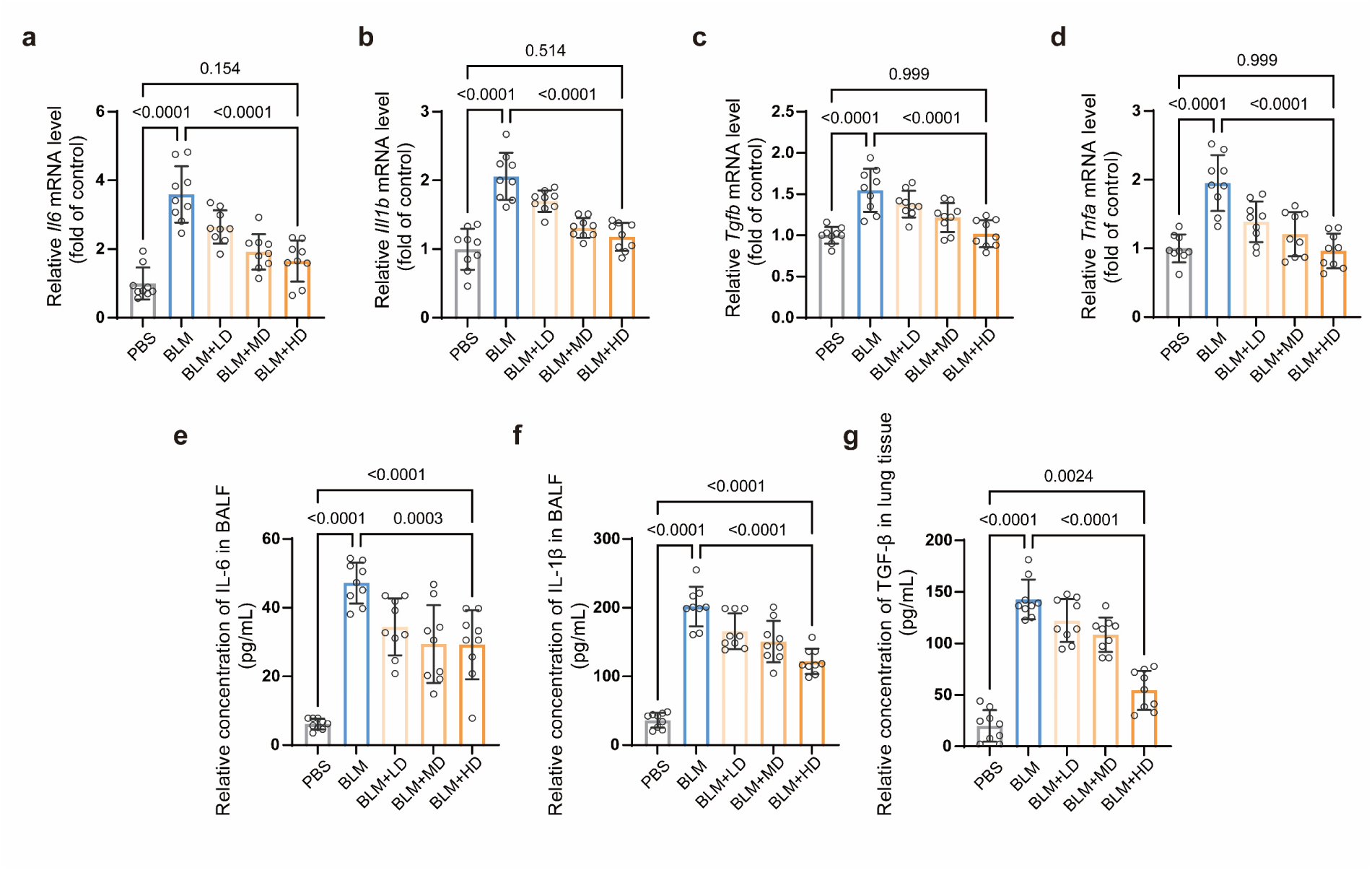
Molecular analysis confirms broad anti-inflammatory effects of rosmarinic acid treatment, revealing dual therapeutic mechanisms in pulmonary fibrosis. (**a**-**d**) Relative mRNA expression of inflammatory mediators in lung tissues *Il6*, *Il1b*, *Tgfb*, and *Tnfa*. These significant reductions in pro-inflammatory cytokine expression demonstrate that RA treatment addresses not only the mechanical and structural aspects of fibrosis but also the inflammatory component of disease initiation and progression. (**e**-**g**) Protein levels in bronchoalveolar lavage fluid (BALF) and lung tissue of inflammatory cytokines IL-1β, IL-6, and TGF-β. The decrease in these inflammatory mediators reveals how AGE inhibition impacts multiple aspects of the disease process, including the TGF-β pathway that drives fibroblast activation and matrix production. This dual action on both the mechanobiological feedback loop and the inflammatory environment may underlie the efficacy of RA in reversing established fibrosis. Statistical analysis performed using one-way ANOVA with Tukey’s post-hoc test, exact *p* values labeled. Results presented as mean ± S.D. (*n* = 9, biologically independent mice per group).

**Extended Data Fig. 14.**
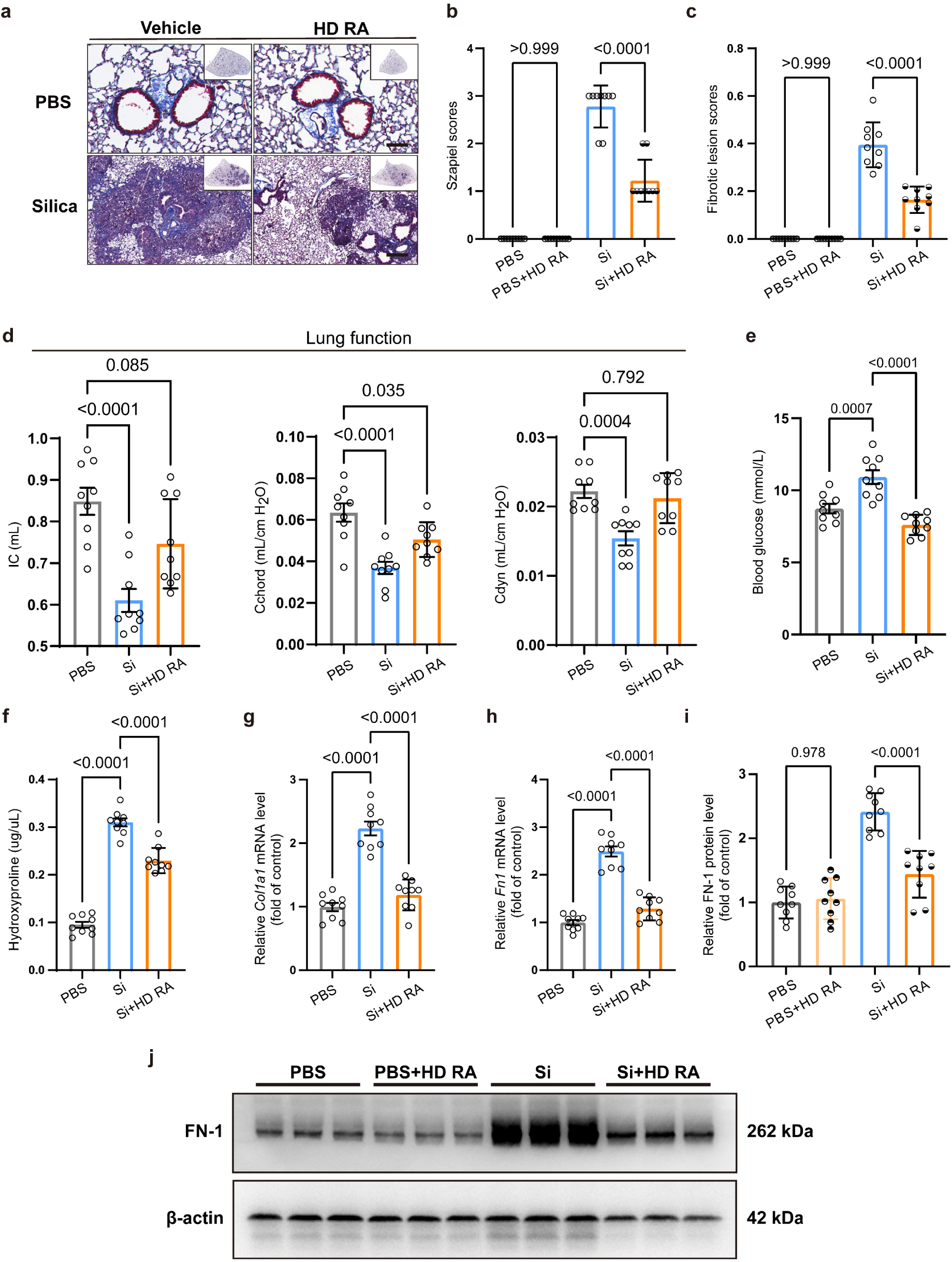
Rosmarinic acid effectively treats silica-induced pulmonary fibrosis, demonstrating generalizability of the AGE-targeting approach across different fibrosis etiologies. (**a**) Representative images of lung tissue sections stained with Masson’s trichrome from silicosis mice treated with vehicle or HD RA, demonstrating reduced fibrotic changes. Scale bars, 100 μm. (**b**, **c**) Histological quantification of fibrosis severity based up (**b**) H&E and (**c**) Masson’s trichrome staining of lungs treated with vehicle or varying RA doses (*n* = 8 mice per group, related to Fig. 6h). These quantitative assessments confirm the robust anti-fibrotic effects of RA in a mechanistically distinct model of pulmonary fibrosis, suggesting broad applicability of AGE inhibition. (**d, e**) Functional improvements following high-dose (HD) RA treatment. (**d**) Restoration of lung function parameters: inspiratory capacity (IC), dynamic compliance (Cdyn), and chord compliance (Cchord). (**e**) Normalization of blood glucose levels. These functional improvements parallel those observed in the bleomycin model, indicating that AGE-mediated crosslinking drives pathology across diverse forms of pulmonary fibrosis. (**f**) Quantification of hydroxyproline content in lung tissue as a measure of collagen deposition. (**g, h**) Relative mRNA expression in lung tissues of key genes showing fibrosis- associated markers *Col1a1*, *Fn1*. (**i**, **j**) Western blot analysis and densitometric quantification of fibronectin (FN-1) expression normalized to β-actin, showing reduced fibrotic protein expression following RA treatment. The efficacy of RA in both bleomycin and silica models provides evidence that the mechanobiological feedback loop involving AGE crosslinking is fundamental to fibrosis progression. Statistical analysis performed using one-way ANOVA with Tukey’s post-hoc test, exact *p* values labeled. Results presented as mean ± S.D. (*n* = 9, biologically independent mice per group).

**Extended Data Fig. 15.**
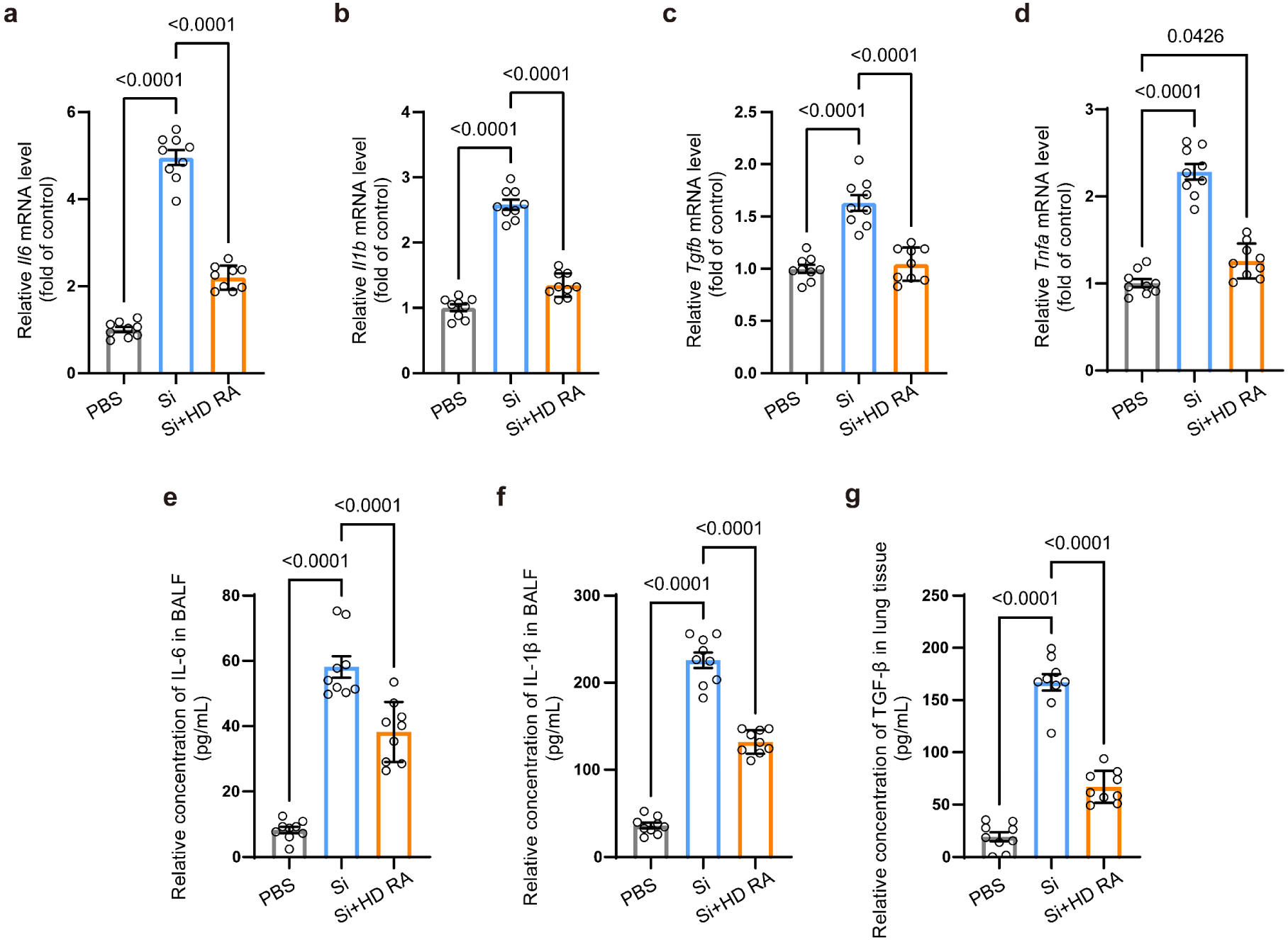
Detailed molecular analysis confirms rosmarinic acid efficacy in silica-induced pulmonary fibrosis through modulation of inflammatory pathways. (**a-d**) Relative mRNA expression in lung tissues of key genes showing inflammatory mediators *Il6*, *Il1b*, *Tgfb*, and *Tnfa*. These significant reductions in pro-inflammatory cytokine expression across multiple fibrosis models suggest that AGE inhibition addresses core inflammatory mechanisms. (**e-g**) Protein levels of inflammatory cytokines IL-6, IL-1β, and TGF-β in bronchoalveolar lavage fluid (BALF) or lung tissues, demonstrating reduced inflammation following RA treatment. The ability of RA to normalize TGF-β levels in silicosis parallels its effects in bleomycin-induced fibrosis, reinforcing the concept that AGE-mediated crosslinking represents a convergent mechanism across different forms of pulmonary fibrosis. Statistical analysis performed using one-way ANOVA with Tukey’s post-hoc test, exact *p* values labeled. Results presented as mean ± S.D. (*n* = 9 biologically independent mice per group).

**Extended Data Fig. 16.**
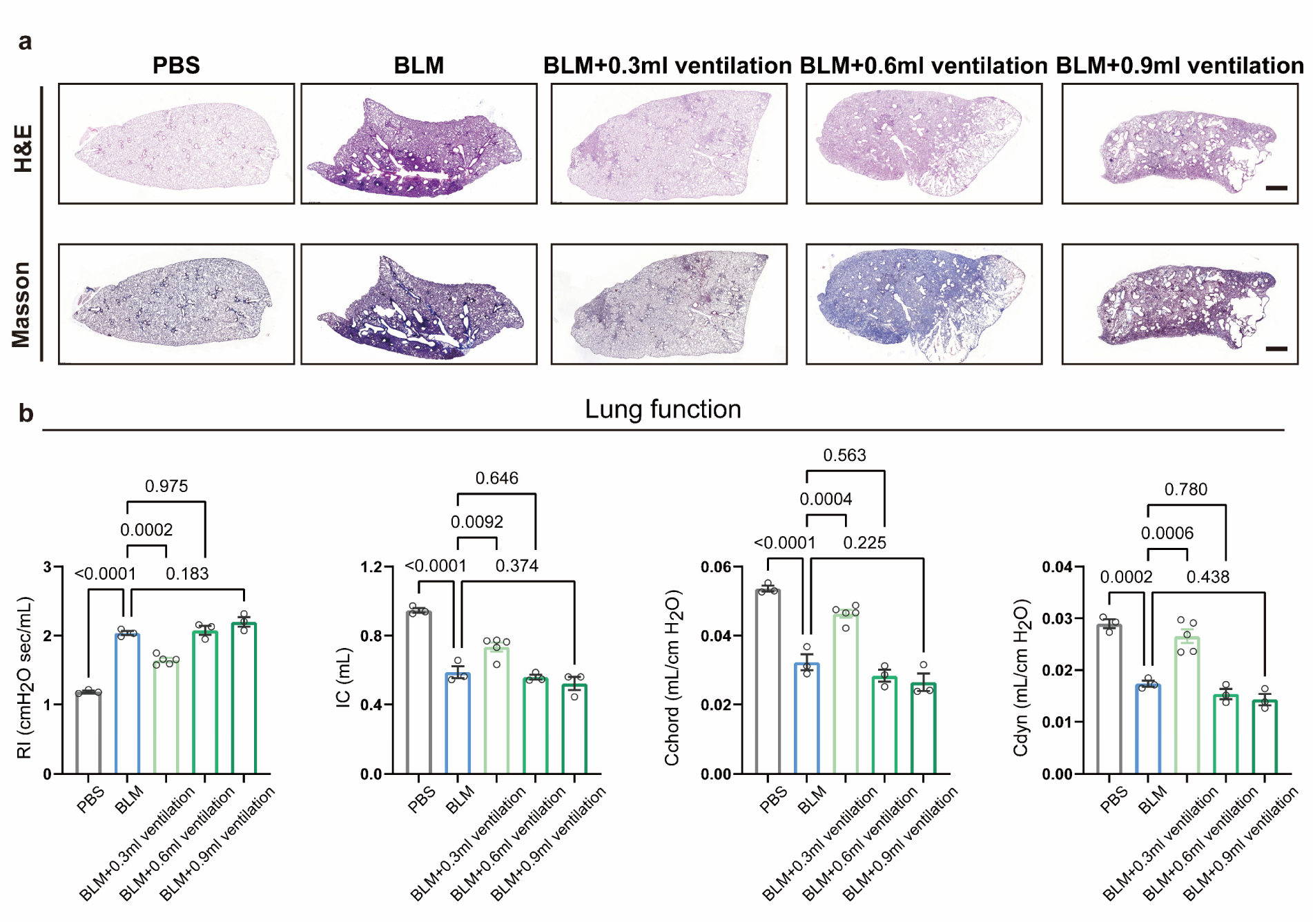
Examination of the therapeutic effects of different gas ventilation volumes in treatment reveals optimal mechanical parameters for mechanotherapy. (**a**) Representative images of collagen deposition stained by H&E and Masson staining from lungs treated with vehicle and different gas volumes of ventilation (0.3ml, 0.6ml, 0.9ml). Scale bars, 1000 μm. Data demonstrate that mechanotherapy requires calibration, with lower ventilation volumes providing therapeutic benefit but higher volumes possibly causing injury. (**b**) Lung function was adversely affected by BLM treatment, but effects reduced to levels that were not significant statistically with low ventilation volume. Factors studied were resistance index (RI), inspiratory capacity (IC), dynamic compliance (Cdyn), and chord compliance (Cchord) (*n* ≥ 3 biologically independent mice per group). These results confirm that calibrated mechanotherapy can restore normal respiratory mechanics, but that excessive mechanical strain may counteract these benefits. Statistical analysis performed using one-way ANOVA with Tukey’s post-hoc test, exact *p* values labeled. Results presented as mean ± S.D.

**Extended Data Fig. 17.**
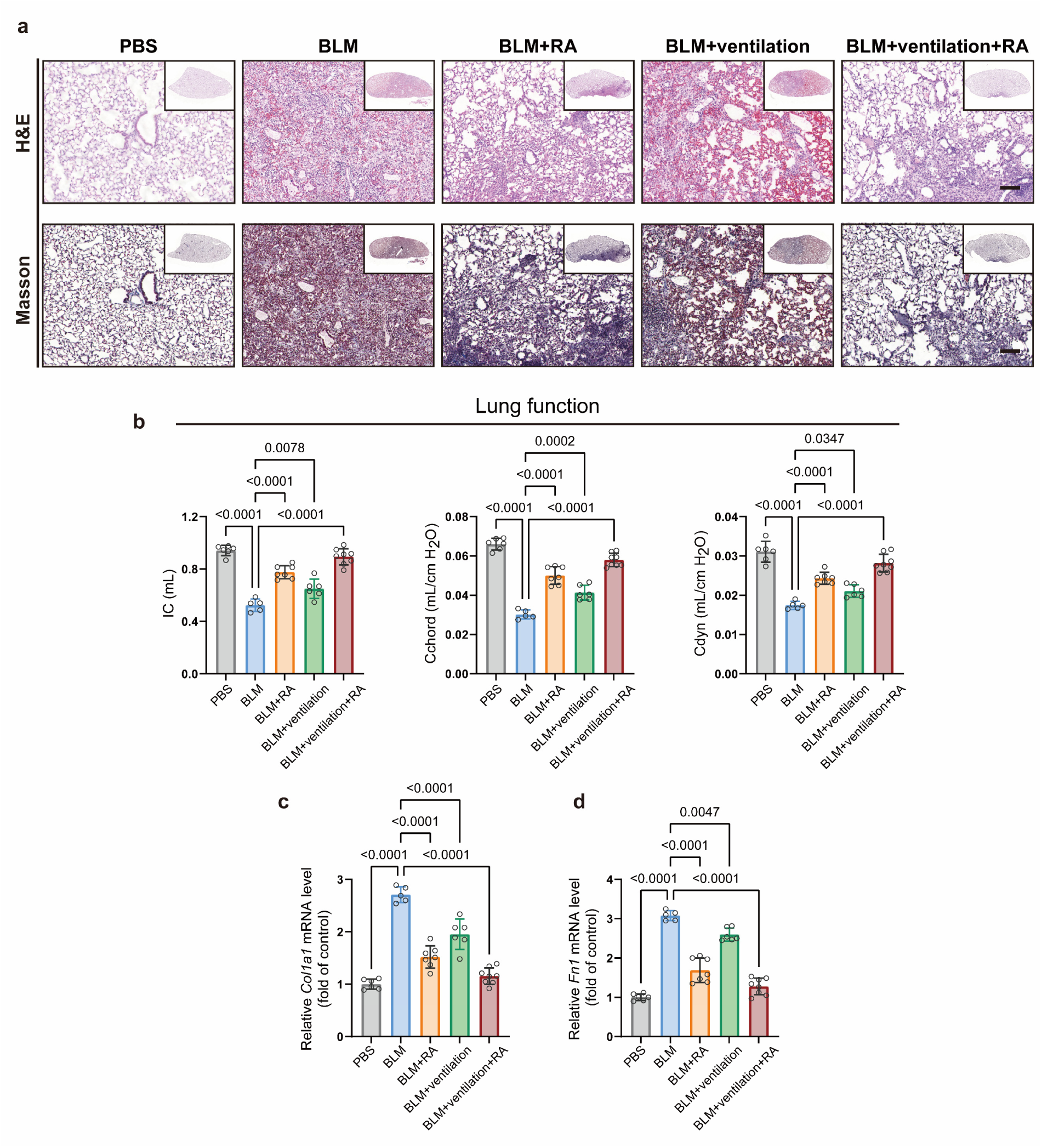
Molecular analysis confirms broad anti-fibrotic effects of mechanical ventilation treatment, establishing mechanotherapy as a therapeutic approach for pulmonary fibrosis. (**a**) Representative lung sections stained with H&E (top) and Masson’s trichrome (bottom) showing reduced fibrotic remodeling in ventilation and combination therapy groups. Scale bars, 100 μm. These data confirm that controlled mechanotherapy can reverse established fibrosis, with maximal benefit observed when combined with AGE inhibition. (**b**) Pulmonary function parameters showing restoration of lung mechanics with combination therapy: inspiratory capacity (IC), dynamic compliance (Cdyn), and chord compliance (Cchord). These functional improvements validate that structural changes induced by mechanotherapy translate to meaningful physiological benefit, particularly with combined therapy. (**c**, **d**) RT-qPCR analysis of fibrosis-associated genes *Acta2* and *Col1a1* in lung tissue showing suppression of fibrotic markers with treatment (*n* ≥ 5 biologically independent mice per group), again confirming that mechanotherapy modulates the same core pathways as RA-based pharmacological AGE inhibition. Statistical analysis performed using one-way ANOVA with Tukey’s post-hoc test. Exact *p* values are indicated. Data presented as mean ± S.D.

**Extended Data Fig. 18.**
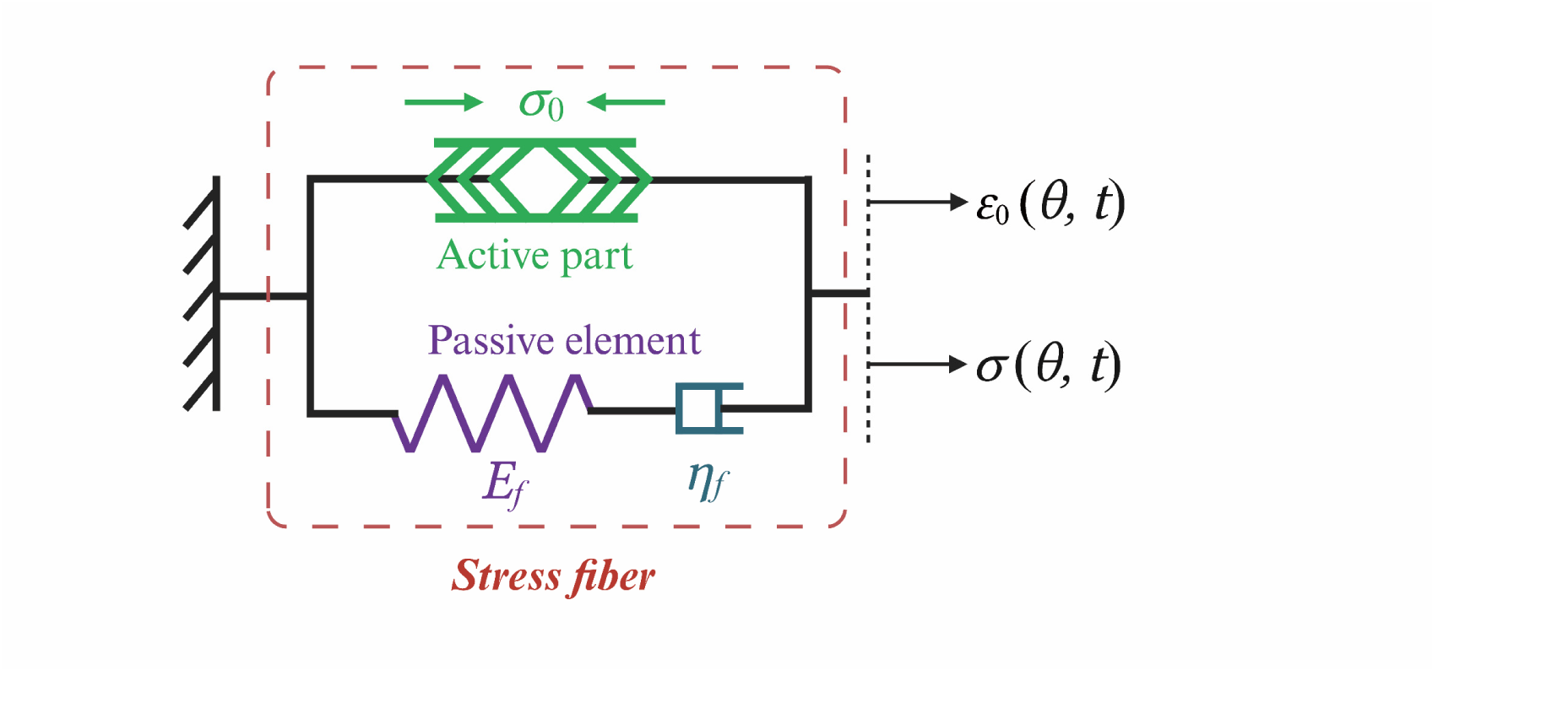
Theoretical model of actomyosin stress fiber dynamics under cyclic mechanical stretching. Schematic of the Hill-type model used to analyze stress fiber and focal adhesion response to cyclic stretch. The model incorporates both active contractile elements (𝜎_0_) and passive viscoelastic components (𝐸_ƒ_, 5_ƒ_), enabling prediction of how cells respond to different cyclical stretch amplitudes. This mechanical framework was applied to explain the biphasic cellular response observed experimentally, where moderate cyclical stretch (5%) enhances activation while higher physiologic cyclical stretch (20%) leads to cytoskeletal remodeling and reduced activation.

