## Supplementary information including Sup Note, Sup Figures 1-2, Sup Tables 1-4 for "Chemo-mechanotherapy of fibrosis: dynamic control of biological materials through tissue-architecture-dependent crosslink disruption"

**Contents**

Supplementary Note
Supplementary Figures 1-2
Supplementary Tables 1-4
Supplementary Videos 1-3 Legends

**Other supplementary information for this manuscript includes the following:**

Supplementary Videos 1-3

### Supplementary Note

**Pore size variations do not affect fibroblast activation**. Comparison of fibroblast behavior on non-crosslinked (NS) and AGE-crosslinked (AS) scaffolds revealed distinct differences in cell activation. A potential confounding variable was the difference in pore size between these scaffolds (36 µm for NS versus 52 µm for AS), which raised the question of whether architectural differences rather than crosslinking status might drive the observed cellular responses.

To isolate the effects of pore size from those of crosslinking and matrix elasticity, we engineered a series of non-crosslinked scaffolds with controlled pore architectures. Through modulation of DMSO concentration during freeze-drying, we regulated ice crystal nucleation and growth to generate scaffolds with pore sizes ranging from 20-40 µm. This architectural control was validated through both SEM imaging and quantitative morphometric analysis (**Supplementary Figures. 1a-b**).

Primary pulmonary fibroblasts (pPFBs) cultured on these scaffolds, showed consistent cellular phenotypes across all pore size conditions. Neither fibroblast activation markers nor RAGE expression showed statistically significant variation despite the substantial differences in scaffold architecture (**Supplementary Figures. 1c-d**). This control experiment establishes that the cellular responses documented in our main studies can be specifically attributed to AGE crosslinking status rather than to differences in the physical architecture of the scaffold microenvironment.

**Matrix architecture determines cellular response to dynamic stretching.** To dissect the specific influence of matrix architecture on mechanotransduction, we first isolated the direct effects of dynamic stretching from matrix-mediated effects. We developed a simplified 2D experimental system where primary pulmonary fibroblasts (pPFBs) were cultured on stretching chambers coated with either non-crosslinked or AGE-crosslinked collagen fibrils (**Supplementary Figures. 2a**). This approach preserved the cell-collagen biochemical interface while eliminating 3D architectural variables present in both scaffold and hydrogel systems.

In this reduced-complexity 2D environment, AGE-crosslinked collagen consistently promoted fibroblast activation, as demonstrated by significant upregulation of canonical activation markers (*Acta2*, *Fn1*, *Col1a1*) and adhesion molecules (*Itga1*, *Itga11*) (**Supplementary Figures. 2b-f**). Application of dynamic stretching (20% strain) counteracted these pro-fibrotic effects, reducing activation marker expression while simultaneously increasing *Ager* expression (**Supplementary Figures. 2g**). These results establish that dynamic mechanical stimulation directly modulates cell behavior independent of the surrounding matrix architecture.

However, comparison across different model systems revealed that matrix architecture significantly modulates the efficiency of this mechanical signaling. In porous scaffolds that mimic lung ECM, dynamic stretching effectively attenuated fibroblast activation, consistent with our 2D findings. In contrast, fibrous hydrogels that model liver-like ECM showed minimal response to the same mechanical stimulation, suggesting fundamentally different mechanotransduction in these environments. This architectural dependence likely stems from differential force transmission mechanisms: porous scaffolds maintain their structural integrity under strain, enabling consistent mechanotransduction, whereas fibrous hydrogels undergo substantial fiber realignment and reorganization (**Fig. 2g-h**) that may dissipate or redirect mechanical signals.

Furthermore, dynamic stretching of porous scaffolds conferred the additional benefit of simultaneously reducing both AGE crosslinking density and matrix elasticity (**Fig. 4f-g**). This dual-action mechanism, combining direct cellular mechanotransduction with beneficial matrix remodeling, explains the superior therapeutic response observed in porous scaffolds compared to fibrous hydrogels, where only direct mechanical effects are possible without significant matrix modification. These findings highlight the importance of tissue-specific ECM architecture in determining therapeutic responses to potential mechanical interventions in fibrotic disease.

### Supplementary Figures

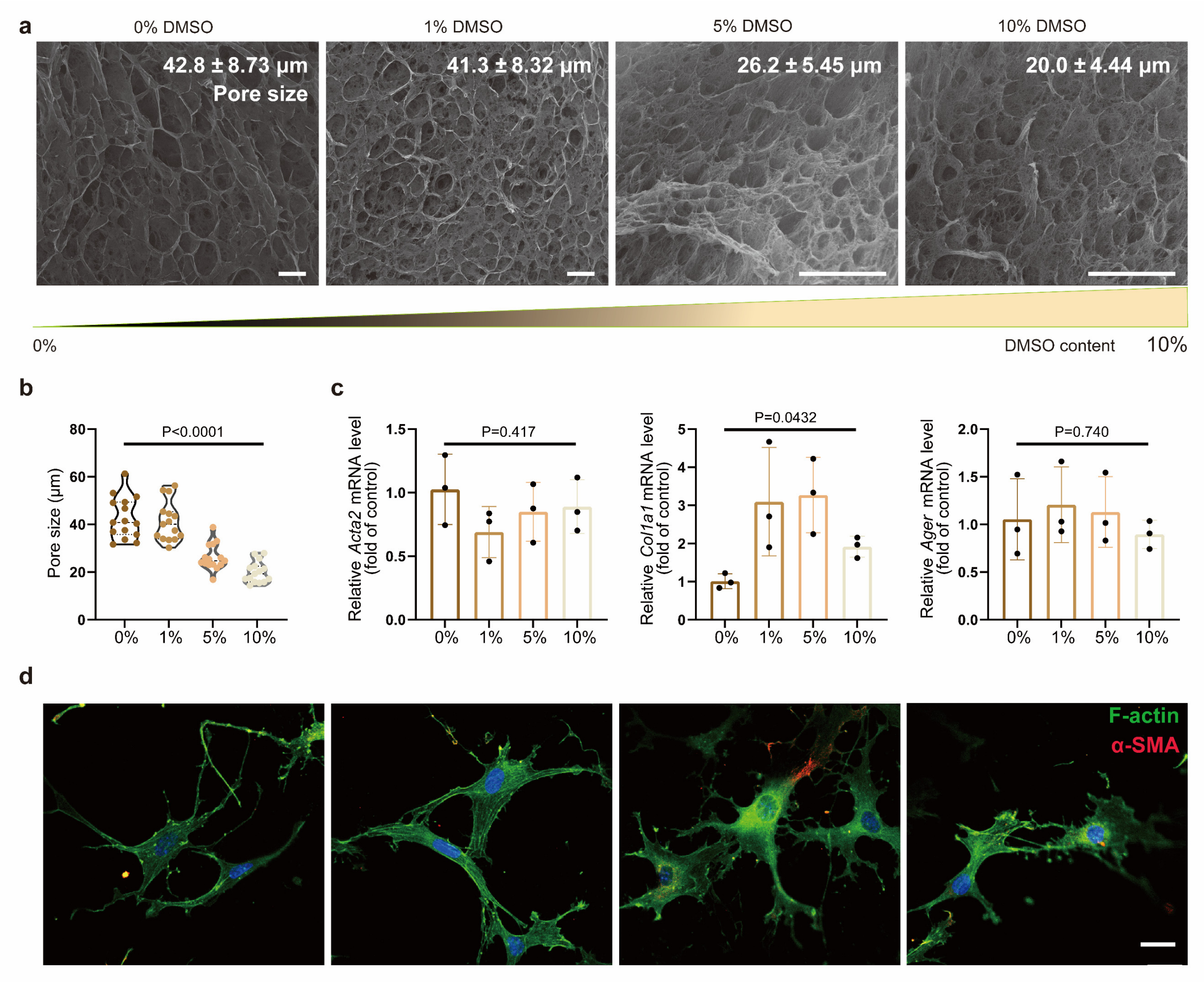

**Supplementary Figure 1. Pore size variation does not affect pulmonary fibroblast activation status, confirming that observed cellular responses are due to AGE crosslinking.** (a) SEM images demonstrating controlled modulation of scaffold pore size through varied DMSO content during fabrication. Scale bars, 100 μm. (**b**) Quantitative analysis of pore sizes achieved by different DMSO concentrations during scaffold synthesis ($n$ = 15 measurements from ≥3 SEM fields). (**c**) Quantification of mRNA expression for key fibrosis-associated genes (*Acta2*, *Col1a1*, *Ager*) in pulmonary fibroblasts cultured on scaffolds with different pore sizes, showing no significant variation despite architectural differences. (**d**) Representative immunofluorescence images of pulmonary fibroblasts on scaffolds with different pore sizes, showing consistent expression patterns of α-SMA (red), F-actin (green), and nuclei (DAPI, blue) regardless of pore size. Scale bar, 20 μm. This control experiment confirms that the cellular activation observed in the main experiments (Figure. 4) can be specifically attributed to AGE crosslinking rather than to differences in scaffold architecture, validating our mechanobiological model of pulmonary fibrosis progression. Statistical analysis performed using one-way ANOVA with Tukey's post-hoc test, exact $p$ values labeled. Results presented as mean ± S.D.

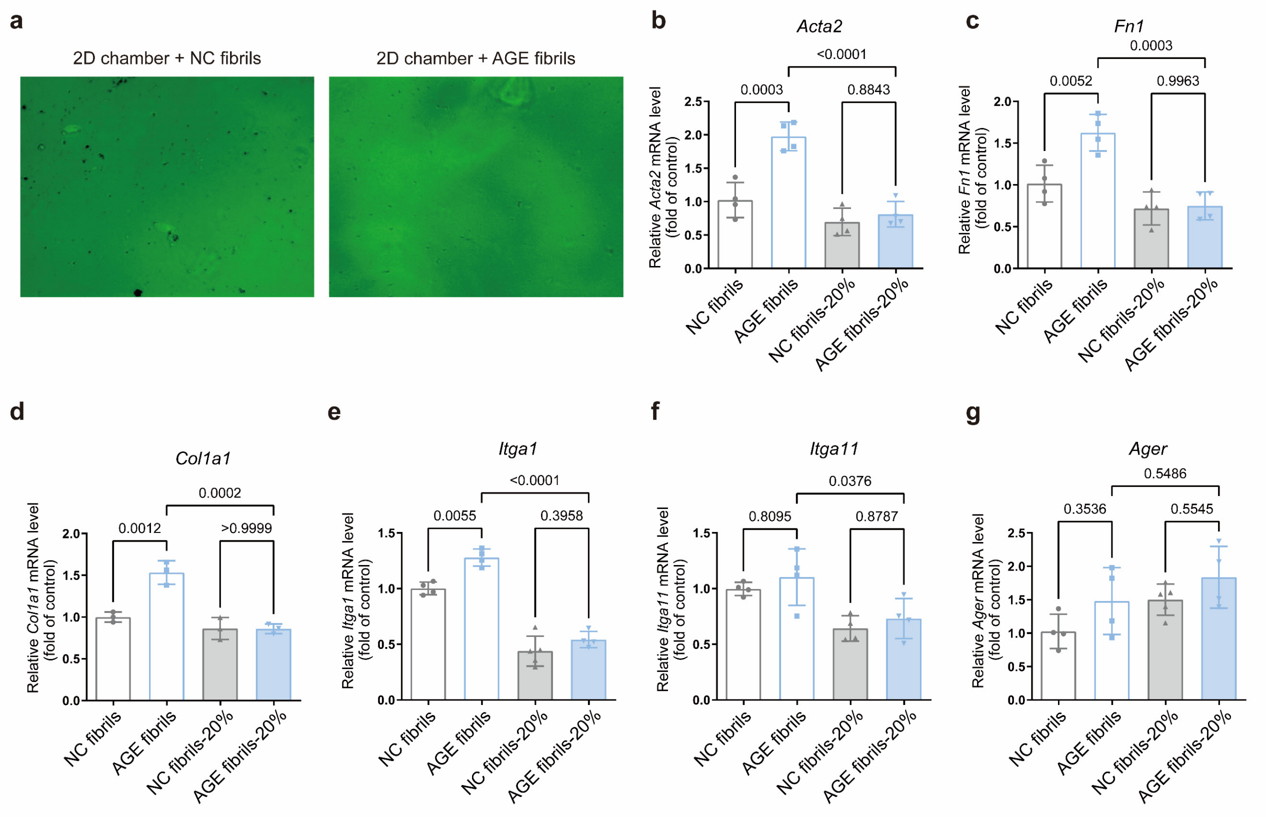

**Supplementary Figure 2. Matrix architecture determines mechanotransduction efficiency, with dynamic stretching directing modulates fibroblast activation in 2D substrates independent of 3D architecture.** (**a**) Representative images showing 2D stretching chamber substrates uniformly coated with 100 μg/mL of either non-crosslinked (NC) or AGE-crosslinked collagen fibrils (green), providing a simplified *in vitro* system to isolate direct mechanical effects from matrix architectural influences. (**b-g**) Quantification of relative mRNA expression in pulmonary fibroblasts (pPFBs) cultured on coated 2D substrates with or without 20% dynamic stretching: (**c**) *Acta2*, (**c**) *Fn1*, (**d**) *Col1a1* (activation marker), (**e**) *Itga1*, (**f**) *Itga11* (adhesion molecule), and (**g**) *Ager* (AGE receptor). Results demonstrate that AGE-crosslinked fibrils promote fibroblast activation even in a 2D context, while dynamic stretching directly counteracts these pro-fibrotic effects by suppressing activation markers and increasing RAGE expression. These findings complement the 3D scaffold results in Figure. 4 by confirming that mechanical signaling occurs independently of matrix architecture, while the magnitude of response depends on matrix structure (porous vs. fibrous) as shown in Supplementary Figure 7. Statistical analysis performed using one-way ANOVA with Tukey's post-hoc test, exact $p$ values labeled. Results presented as mean ± S.D.

**Supplementary Tables**

**Supplementary Table 1. Information of clinical human lung samples.**

| Group | ID | Age | Sex | Case diagnosis | Complications and concomitant diseases |
|---|---|---|---|---|---|
| IPF cases | 1 | 64 | M | Interstitial lung disease, likely IPF | Coronary atherosclerotic heart disease, coronary stent placement; carotid atherosclerosis with plaque formation, stiffening of the aortic valve |
| | 2 | 68 | M | Interstitial lung disease, IPF, Type I respiratory failure, invasive aspergillosis, CMV pneumonia, | Mediastinal emphysema; atherosclerosis; reflux esophagitis; steroid diabetes; sleep disorders; osteoporosis |
| | 3 | 58 | M | IPF | None |
| | 4 | 51 | M | Idiopathic interstitial lung disease | Type 2 diabetes |
| | 5 | 62 | M | IPF | Pulmonary hypertension secondary to hypoxia; New York Heart Association Class IV heart failure |
| | 6 | 58 | M | Idiopathic pulmonary interstitial fibrosis, right pulmonary bulla | Elevated tumor markers |
| | 7 | 69 | M | IPF, Type I respiratory failure | Steroid diabetes mellitus; hyperlipemia; chronic gastritis; allergic rhinitis; reflux esophagitis; severe osteoporosis; coronary atherosclerosis; carotid atherosclerosis with plaque formation |
| | 8 | 65 | M | Interstitial lung disease, IPF, Type I respiratory failure | Congenital heart disease; ventricular septal defect (perimembranous), ventricular septal membranous tumor; coronary atherosclerosis; arrhythmia (sinus tachycardia); hyperlipidemia; fatty liver |

| | | | | | |
|---|---|---|---|---|---|
| | 9 | 68 | M | Idiopathic pulmonary interstitial fibrosis, Type I respiratory failure | Type 2 diabetes; chronic gastritis duodenal ulcer |
| | 10 | 72 | M | Interstitial lung disease, IPF, Type I respiratory failure) | fatty liver; abnormal liver function; atherosclerosis of the carotid artery |
| | 11 | 64 | M | Interstitial lung disease, high likelihood of IPF, pulmonary hypertension, Type I respiratory failure | Coronary atherosclerotic heart disease, post-PTCA, hyperlipidemia |
| | 12 | 72 | M | Interstitial lung disease, IPF, Type I respiratory failure, pulmonary hypertension (moderate) | Chronic pulmonary heart disease; coronary atherosclerosis; carotid atherosclerosis; elevated tumor markers |
| | 13 | 67 | M | Interstitial lung disease, IPF, Type I respiratory failure | Coronary atherosclerotic heart disease; right coronary artery dissection, right coronary artery stent implantation; old cerebral infarction; carotid plaque formation; hyperlipidemia; left femur fracture after plate fixation |
| | 14 | 61 | M | IPF | None |
| | 15 | 58 | M | IPF | Liver cirrhosis |
| | 16 | 59 | M | Idiopathic nonspecific interstitial pneumonia (fibrotic type), Type I respiratory failure, lung infection (cytomegalovirus, respiratory syncytial virus), pulmonary embolism (low risk group), pulmonary hypertension | Abnormal liver function; electrolyte disturbance; hypokalemia; reflux esophagitis; gastric ulcer |
| | 17 | 65 | M | Idiopathic interstitial lung disease | Status post-appendectomy |
| | 18 | 63 | M | IPF | None |

| Group | ID | Age | Sex | Case diagnosis | Complications and concomitant diseases |
|---|---|---|---|---|---|
| **Normal controls** | 19 | 58 | M | Lung space occupying lesion (peripheral healthy tissue) | |
| | 20 | 72 | M | Donor | |
| | 21 | 65 | M | Lung space occupying lesion (peripheral healthy tissue) | |
| | 22 | 65 | M | Donor (smoker) | |
| | 23 | 59 | M | Donor | |
| | 24 | 65 | F | Lung space occupying lesion (peripheral healthy tissue) | |
| | 25 | 40 | M | Normal tissue adjacent to tumor | |
| | 26 | 69 | F | Normal tissue adjacent to tumor | |
| | 27 | 48 | M | Donor | |
| | 28 | 34 | M | Donor | |
| | 29 | 62 | M | Lung space occupying lesion (peripheral healthy tissue) | |
| | 30 | 45 | F | Lung space occupying lesion (peripheral healthy tissue) | |

| | | | | |
|---|---|---|---|---|
| | 31 | 63 | F | Lung space occupying lesion (peripheral healthy tissue) |
| | 32 | 56 | F | Lung space occupying lesion (peripheral healthy tissue) |
| | 33 | 64 | M | Lung space occupying lesion (peripheral healthy tissue) |

**Supplementary Table 2. MS2 fragmentation spectra of the crosslinking products used for detection.**

| Compound | Q1 (m/z) | Q3 (m/z) | RT (min) | DP (V) | CE (V) |
|---|---|---|---|---|---|
| Glucosepane | 429.2 | 384.3 | 5.85 | 180 | 45 |
| CML | 205.0 | 130.1 | 6.97 | 80 | 18 |
| PYD | 429.1 | 412.2 | 7.46 | 180 | 36 |
| CEL | 219.1 | 130.1 | 6.75 | 95 | 19 |
| Pentosidine | 379.1 | 187.1 | 5.64 | 150 | 46 |
| HYP | 132.1 | 86.1 | 3.87 | 65 | 16 |
| γ-GLU-ε-LYS | 276.3 | 147.2 | 6.87 | 70 | 46 |

*Q1, Q3: the first and third quadrupoles act as mass filters; *RT: retention time; *DP: declustering potential; *CE: collision energy

**Supplementary Table 3. Primer set for gene expression analysis using real-time qPCR.**

| Species | Gene | primer |
|---|---|---|
| Mouse | *bactin*-F | TAGGCACCAGGGTGTGAT |
| | *bactin*-R | CTCCTCAGGGGCCACA |
| | *Col1a1*-F | CCTGGTCCCTCTGGAAATG |
| | *Col1a1*-R | GGAAGCCTCTTTCTCCTCTC |
| | *Fn1*-F | CCCTATCTCTGATACCGTTGTCC |
| | *Fn1*-R | TGCCGCAACTACTGTGATTCGG |
| | *Acta2*-F | TGCTGACAGAGGCACCACTGAA |
| | *Acta2*-R | CAGTTGTACGTCCAGAGGCATAG |
| | *Ager*-F | CCACTGGAATTGTCGATGAGG |
| | *Ager*-R | CTCGGACTCGGTAGTTGGACT |
| | *Itga1*-F | TGGCTTCTCACCGTTATCCTA |
| | *Itga1*-R | CACACAAGGCATTGATCTCTCT |
| | *Itga11*-F | TGCCCCAATGGAAACCAATG |
| | *Itga11*-R | CATGCCAGTGGTGTAGTAGGA |
| | *Il6*-F | TTCCTCTCTGCAAGAGACTTC |
| | *Il6*-R | GTTGGGAGTGGTATCCTCTG |
| | *Il1b*-F | CAAGCTTCCTTGTGCAAGTGTC |
| | *Il1b*-R | TTCATCTTTTGGGGTCCGTCA |
| | *Tnfa*-F | AGACACCATGAGCACAGAAA |
| | *Tnfa*-R | CACTTGGTGGTTTGTGAGTG |
| | *Tgfb*-F | CGTGGAAATCAACGGGATCA |
| | *Tgfb*-R | TCCAAATATAGGGGCAGGGT |

**Supplementary Table 4. Parameters used in the analysis and simulations.**

| Symbol | Physical meaning | Value |
|---|---|---|
| $k_+^b$ | Association rate of integrin-RGD bonds | $2\pi\ \mathrm{s}^{-1}$ |
| $k_+^f$ | Association rate of contractile units | $0.4\pi\ \mathrm{s}^{-1}$ |
| $k_-^f$ | Disassociation rate of contractile units | $0.2\pi\ \mathrm{s}^{-1}$ |
| $U_b^*$ | Dimensionless energy release associated with a single integrin-RGD bond | 5 |
| $U_{int}^*$ | Dimensionless additional energy associated with actomyosin recruitment to the growing focal adhesion | 3.5 |
| $\varepsilon_c$ | Critical strain | 0.02 |
| $\nu$ | Poisson ratio | 0.5 |

**Supplementary Videos**

**Supplementary Video 1.** Dynamic stretching system.

**Supplementary Video 2.** Computational simulations of stretch-induced fibrous hydrogel fiber alignment.

**Supplementary Video 3.** Computational simulations of stretching did not alter porous scaffold pores.
